# Microbial production of the low-caloric sweetener D-allulose from D-glucose by evolutionary engineering

**DOI:** 10.1101/2024.12.16.628640

**Authors:** Alexander Lehnert, Rocco Gentile, Claudia Tahiraj, Astrid Wirtz, Meike Baumgart, Tino Polen, Holger Gohlke, Michael Bott

## Abstract

The low-calorie sugar D-allulose is a promising alternative to D-sucrose and high-fructose corn syrup, but its microbial production from D-glucose at mesophilic temperatures is limited by insufficient D-glucose isomerase (XylA) activity. Here, we overcome this bottleneck by evolving a *Corynebacterium glutamicum* selection strain whose growth strictly depends on XylA function. This strategy yielded a XylA variant with a nine-fold higher catalytic efficiency, sugar transporter variants (IolT1) with ten-fold increased activity for D-glucose and D-fructose, and hints for co-transport of these sugars by the D-sucrose transporter PtsS. Molecular dynamics simulations provided possible mechanistic explanations for the adaptive mutations. Combining the evolved enzymes with a suitable D-allulose 3-epimerase in a highly engineered chassis strain enabled whole-cell conversion of D-glucose to D-allulose with a 15% yield at 30 °C. This performance rivals immobilized enzyme processes performed at ∼60°C while avoiding enzyme purification and immobilization, offering an alternative for low-calorie sweetener production.

## Introduction

The overconsumption of food and beverages sweetened with high-caloric D-sucrose or high-fructose corn syrup (HFCS) causes numerous health problems, such as obesity, type 2 diabetes, and cardiovascular diseases [1,2]. In order to reduce the consumption of D-sucrose and HFCS, numerous low-caloric sweeteners have been introduced, such as saccharine, aspartame, xylitol, or stevioside, several of which have undesired traits [3]. D-Allulose, also named D-psicose, is a rare natural sugar, the C3 epimer of D-fructose, that gained strong interest as a dietary sweetener, as it has 70% of the sweetness of D-sucrose, but less than 10% of its caloric content (≤0.4 kcal/g) [4]. In 2024, the global D-allulose market was valued at approximately USD 148 million and was projected to grow at a compound annual growth rate (CAGR) of 14% from 2025 to 2034, expected to reach around USD 553 million by 2034 by Global Market Insights Inc. D-Allulose is generally recognized as safe by the FDA (GRAS Notice 400) and its consumption is associated with positive health effects, such as attenuation of blood glucose levels and reduction of body weight [5,6].

The industrial production of D-allulose is performed in enzyme reactors, either from D-fructose via a one-step conversion with D-allulose 3-epimerase or D-tagatose 3-epimerase, or from D-glucose in a two-step conversion including additionally a D-glucose isomerase (XylA) [7]. D-Glucose isomerase is in fact a D-xylose isomerase having the physiological function to convert D-xylose to D-xylulose, which after phosphorylation can be metabolized in the pentose phosphate pathway. As a side-reaction this enzyme also catalyzes the conversion of D-glucose to D-fructose and therefore is also called D-glucose isomerase, which is the term generally used in this study [8]. D-Glucose isomerases are used in the industrial production of HFCS, which is a mixture of D-glucose and D-fructose that is used as a sweetener for beverages and processed foods at a >10 million tons scale per year. HFCS is produced at 60-70°C to compensate for the poor activity of D-glucose isomerases towards D-glucose at mesophilic temperatures and to shift the thermodynamic equilibrium of the isomerization towards D-fructose [9]. D-Glucose isomerase research therefore focused on the isolation of more active and thermostable variants either by protein engineering or by using isoenzymes from hyperthermophilic microorganisms [10–12]. In contrast, no studies have been reported aiming at identifying D-glucose isomerases with improved activity at mesophilic temperatures. In the current industrial D-allulose production from D-glucose, the use of high temperatures of ≥50°C for the efficient isomerization of D-glucose to D-fructose poses a challenge, since D-allulose 3-epimerases or D-tagatose 3-epimerases generally reveal a low stability at this temperature range [13]. Hence, immobilization of the enzymes is used to increase their stability and reusability. Combined with immobilized D-glucose isomerases, the production of D-allulose from D-glucose via isomerization and epimerization achieves yields of 10-16% [14–16].

In this study, we engineered bacterial strains that efficiently produce D-allulose from D-glucose at 30°C via D-glucose isomerase and D-allulose 3-epimerase for applications in the food and beverage industry. Such an approach avoids the necessity of protein purification, protein immobilization, and high operating temperatures (≥50°C), thereby reducing energy and material demands while improving product stability. The major bottleneck was to enable a fast conversion of D-glucose to D-fructose within the cells by D-glucose isomerase. To solve this problem, we employed adaptive laboratory evolution (ALE). As host we chose *Corynebacterium glutamicum*, an industrially well established microbe that is used since decades for amino acid production in the million ton scale and whose products are generally regarded as safe (GRAS) [17,18]. Of particular importance in the context of our study was the previous detailed knowledge on sugar uptake and metabolism in *C. glutamicum* [19]. D-Glucose, D-fructose, and D-sucrose are taken up via phosphoenolpyruvate-dependent phosphotransferase systems (PTS) with the EII transporters PtsG, PtsF, and PtsS that form D-glucose 6-phosphate, D-fructose 1-phosphate, and D-sucrose 6-phosphate, respectively. A small fraction of D-fructose is imported by PtsG forming D-fructose 6-phosphate [20]. D-Sucrose 6-phosphate is converted by D-sucrose 6-phosphate hydrolase to D-glucose 6-phosphate and D-fructose [21]. As *C. glutamicum* lacks fructokinase activity, the imported D-fructose cannot be metabolized and is exported by an unknown transporter followed by reimport via PtsF and PtsG [20]. Besides PTS transport, uptake of certain sugars can also be catalyzed by the H^+^-coupled secondary transporters IolT1 and IolT2. Their native substrates are inositols [22,23], but due to their promiscuity, they can also transport D-fructose [24] and D-glucose [25,26]. In contrast to D-fructose, imported D-glucose can be metabolized by *C. glutamicum* after phosphorylation to D-glucose 6-phosphate by the endogenous glucokinases Glk and PpgK [27,28]. Based on this knowledge, we engineered a selection strain whose growth was strictly dependent on the isomerization of D-fructose to D-glucose by the provided D-glucose isomerase. In a few rounds of ALE, fast-growing clones were obtained and found to harbor mutations in only two targets, XylA and IolT1, causing increased D-glucose isomerase activity and an increased activity for D-fructose uptake into the cytoplasm. The combination of the evolved XylA and IolT1 proteins in a chassis strain unable to grow on D-glucose, D-fructose, and D-sucrose and equipped with a D-allulose 3-epimerase enabled the conversion of D-glucose to D-allulose at 30°C with yields of about 15% that are comparable to the industrial processes using immobilized enzymes at >50°C.

## Results

### Growth-based selection for high *in vivo* D-glucose isomerase activity at 30°C

D-Glucose isomerases have a very low affinity for D-glucose, with K_m_ values typically above 100 mM (Table S1 in the supplemental information online). Whereas in processes with immobilized D-glucose isomerases much higher D-glucose concentrations can be provided to enable high enzyme activity, the D-glucose concentration within microbial cells is usually much lower, as D-glucose is immediately metabolized in the central metabolism. In order to obtain microbial strains enabling a high conversion rate of D-glucose to D-fructose within the cells, we used ALE with a selection strain of *C. glutamicum* whose growth at 30°C was strictly dependent on the isomerization of D-fructose to D-glucose. As first step, the genes for the EII permeases PtsG and PtsF were deleted to prevent uptake of D-glucose and D-fructose as D-glucose 6-phosphate and D-fructose 1-phosphate, respectively, as these metabolites cannot serve as substrates for the D-glucose isomerase. To enable the import of unphosphorylated D-glucose and D-fructose, expression of the *iolT1* gene was made constitutive by relieving its repression by its native transcriptional regulator IolR [29] via promoter mutations [30]. Growth of the resulting Fru^neg^ strain (Figure 1A) was tested in a BioLector microcultivation system, which measures the cell density online as backscatter signal [31]. The strain showed negligible growth in D-fructose minimal medium, while growth in D-glucose minimal medium was only slightly reduced (µ = 0.34 ± 0.02 h^-1^) compared to the parental strain MB001 (µ = 0.45 ± 0.01 h^-1^) due to the lack of PtsG (Figure 1B-C). Based on these results, *C. glutamicum* Fru^neg^ was a suitable seletion strain for the isolation of improved D-glucose isomerase variants.

**Figure 1.**
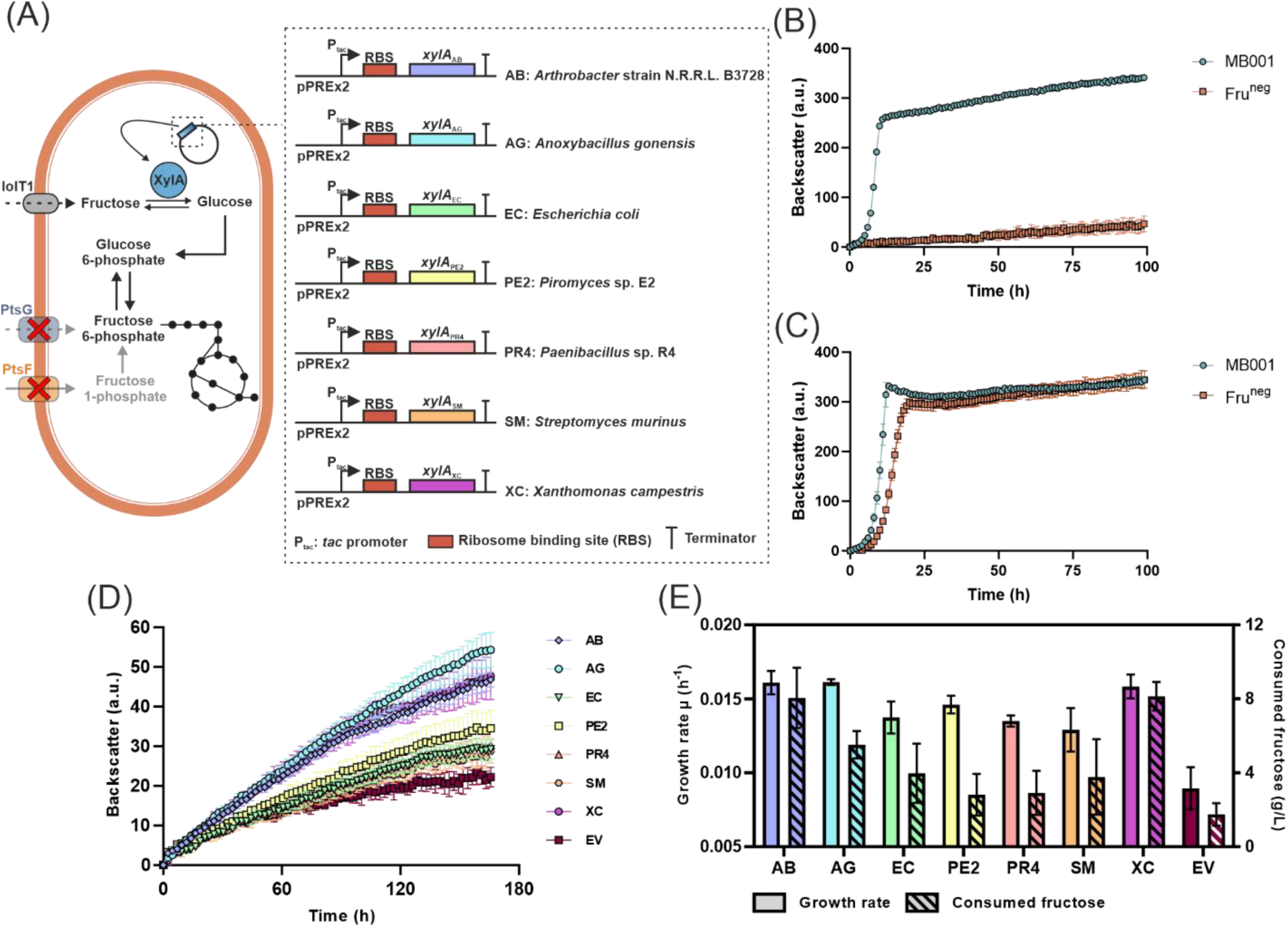
Growth-based selection system for the evaluation of D-glucose isomerase activity. (A) Coupling of D-glucose isomerase activity to the growth of the Fru^neg^ strain. The Fru^neg^ strain and its parental strain MB001 were cultivated in CGXII medium with 20 g/L D-fructose (B) or 20 g/L D-glucose (C) at 30°C and 1200 rpm for 100 h in a BioLector I microcultivation system. (D) Growth of Fru^neg^ transformed with either one of the seven *xylA* expression plasmids or the vector control pPREx2 (EV) in CGXII medium with 40 g/L D-fructose, 25 µg/mL kanamycin and 1 mM IPTG. The cultures were inoculated to an initial OD_600_ of 7.5. e Growth rates and consumed D-fructose titers of Fru^neg^ strains carrying different *xylA* variants shown in panel (d) after 160 h. All data points illustrate the mean ± S.D. from three biological replicates (*n* = 3).

Seven genes for D-glucose isomerases that were previously reported to show activity for D-glucose or D-xylose at 30°C (Table S1 in the supplemental information online) were cloned into plasmid pPREx2 under the control of an IPTG-inducible *tac* promoter and transferred into strain Fru^neg^ (Figure 1A). The resulting strains were then tested for growth on D-fructose in the BioLector. To observe small growth differences, inoculation was performed to a high optical density at 600 nm (OD_600_) of 7.5. The final OD_600_ of the cultures was measured spectrophotometrically and D-fructose titers in the culture supernatant were determined via HPLC. All seven selected D-glucose isomerases improved growth of the Fru^neg^ strain (Figure 1D, Figure S1 in the supplemtal information online), but differed with respect to the growth rate and D-fructose consumption (Figure 1E). The enzymes of *Anoxybacillus gonensis* (XylA_AG_), *Arthrobacter* strain N.R.R.L. B3728 (XylA_AB_) and *Xanthomonas campestris* (XylA_XC_) enabled the highest growth rates in D-fructose minimal medium, but only XylA_AB_ and XylA_XC_ also resulted in a high D-fructose consumption by the Fru^neg^ strain, supporting a direct link between growth and D-fructose isomerization. Hence, XylA_AB_ and XylA_XC_ were selected for the ALE experiments.

### Adaptive laboratory evolution of the Fru^neg^ strain carrying *xylA* expression plasmids

XylA_AB_ and XylA_XC_ slightly improved growth of the Fru^neg^ strain in D-fructose minimal medium, but both corresponding strains barely doubled their initial OD_600_ and consumed less than 20% of the provided D-fructose during 160 h of cultivation, confirming severe bottlenecks in the conversion of D-fructose to D-glucose. To eliminate these bottlenecks, the strains Fru^neg^ pPREx2-*xylA*_AB_ (AB) and Fru^neg^ pPREx2-*xylA*_XC_ (XC) were subjected to laboratory evolution by selecting for faster growth. The ALE experiment was performed with three biological replicates of each strain in 24-well plates (Figure 2A). After 3-5 serial cultivations in D-fructose minimal medium lasting for 5-15 days each over a total period of 45-55 days, the growth rate and the cell density of all six cultures were strongly increased. Then, aliquots of each culture were plated at a 1:100,000 dilution on D-fructose minimal medium agar plates. Small and large colonies appeared after 4-5 days of incubation at 30°C and one single large colony of each plate was picked and used for a growth experiment in the BioLector. All six evolved clones (called FAB evo 1-3 and FXC evo 1-3) grew much faster and reached a much higher OD_600_ than the parental and control strains and consumed the entire D-fructose, showing that evolution was successful (Figure 2B-C, Figure S2A-B in the supplemental information online).

**Figure 2.**
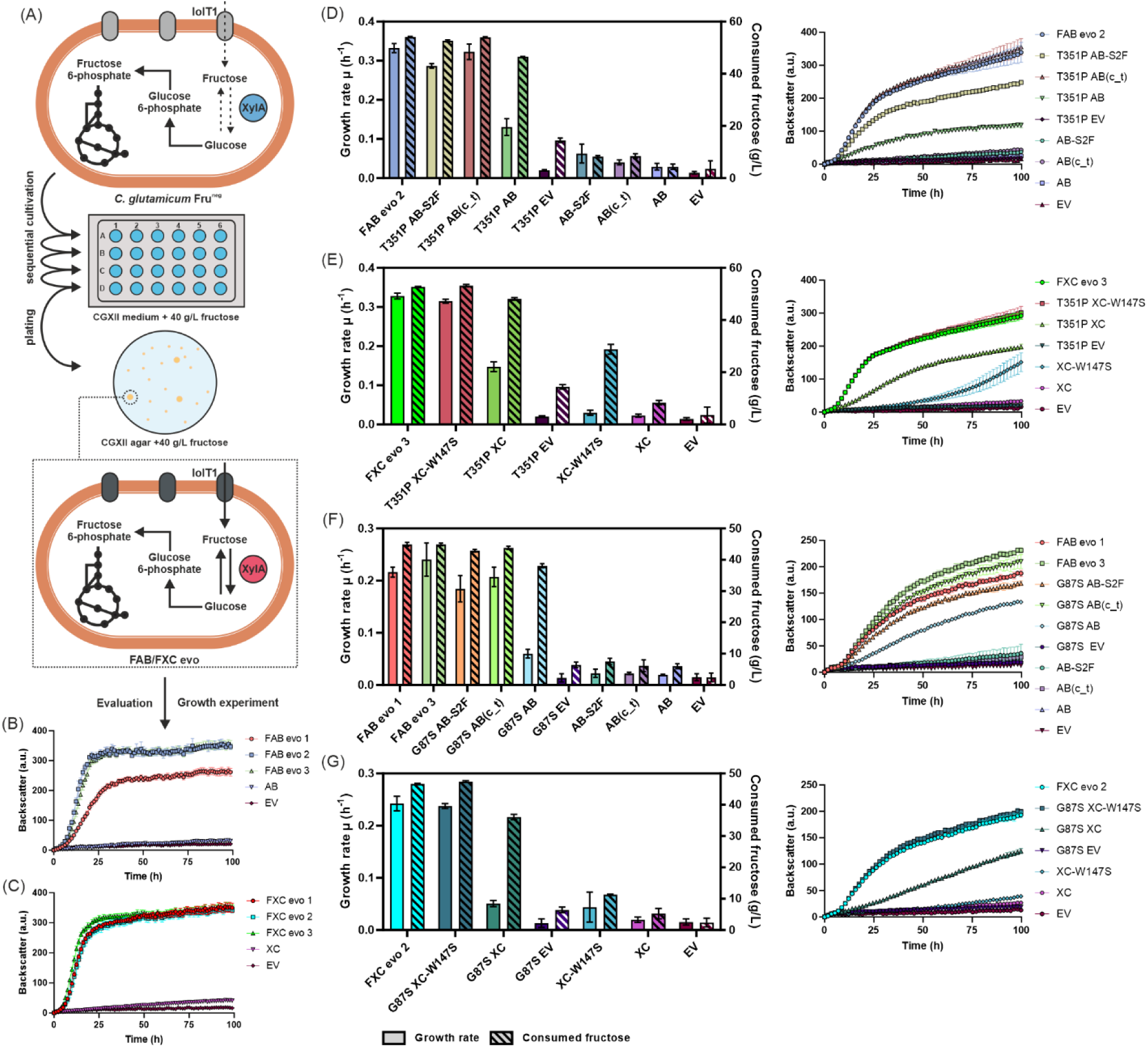
Evolution of Fru^neg^ strains with fast growth in D-fructose minimal medium. (A) Design of the adaptive laboratory evolution (ALE) experiment. Fru^neg^ strains carrying either pPREx2-*xylA*_AB_ (AB) or pPREx2-*xylA*_XC_ (XC) were sequentially cultivated in CGXII medium with 40 g/L D-fructose, 25 µg/mL kanamycin and 1 mM IPTG in 24-well plates at 30°C and 900 rpm. After the ALE was terminated each culture was diluted by a factor of 100,000 and plated on CGXII D-fructose agar plates to obtain single colonies. Comparatively large colonies were selected and used for a growth experiment in the same medium used for ALE (panels b and c). Each ALE was performed in biological triplicates. (B), (C) BioLector II growth experiments of the selected evolved AB and XC strains (FAB evo 1-3, FXC evo 1-3) in comparison to the unevolved parental strains (AB, XC) and the empty vector control (EV). (D), (E) Growth kinetics and D-fructose consumption of reverse engineered AB (D) and XC (E) strains containing *iolT1*-T351P (T351P) with pPREx2-*xylA*_AB_-S2F (AB-S2F), pPREx2-*xylA*_AB_(c_t) (AB(c_t)), pPREx2-*xylA*_XC_-W147S (XC-W147S) or pPREx2 (EV). (F), (G) Growth kinetics and D-fructose consumption of reverse engineered AB (F) and XC (G) strains containing *iolT1*-G87S (G87S) with AB-S2F, AB(c_t), XC-W147S or EV. All growth experiments were conducted in CGXII medium with 40 g/L of D-fructose, 25 µg/mL kanamycin and 1 mM IPTG at 30°C and 1200 rpm for ∼100 h either in BioLector I or II. The data shown represent the mean ± S.D. from three biological replicates (*n* = 3), except for XC (panel c) and EV (panel d-g) for which *n* = 2.

To elucidate the molecular basis for the improved growth of the evolved strains, first their plasmids were sequenced (Table 1). In pPREx2-*xylA*_AB_ of strain FAB evo 1 a mutation causing a S2F exchange in XylA was identified, whereas strains FAB evo 2 and FAB evo 3 both harbored a cytosine to thymine exchange 20 bp upstream of the *xylA*_AB_ start codon in pPREx2-*xylA*_AB_ (6 bp upstream of the ribosome binding site). The latter mutation might cause improved translation of the *xylA* mRNA leading to more XylA protein and higher enzyme activity. The XylA_AB_-S2F mutation might also enable an increased translation of the *xylA* mRNA. Remarkably, the plasmids of the three independently evolved strains with the *X. campestris xylA* gene all carried an identical mutation leading to a W147S exchange in XylA_XC_. We assume that this single mutation increases the enzymatic activity for D-fructose isomerization to D-glucose. Next the genomic DNA of the six evolved strains was sequenced. Interestingly, all genomes were found to carry only a single mutation which was located in the *iolT1* gene (Table 1). Four strains (FAB evo 1, FAB evo 3, FXC evo 1, and FXC evo 2) contained a G87S exchange in IolT1, whereas two strains (FAB evo 2 and FXC evo 3) harbored a T351P exchange in IolT1. The results suggested that the combination of two effects was responsible for enabling improved growth on D-fructose, namely an improved activity of D-glucose isomerase for D-fructose conversion to D-glucose and an improved D-fructose uptake to increase the cytoplasmic D-fructose concentration and thereby XylA activity.

**Table 1.** Plasmid and genome mutations of ALE-derived Fru^neg^ pPREx2-*xylA* strains.

| Strain | Location of mutation | Mutation |
| --- | --- | --- |
| **FAB evo 1** | *xylA*<sub>AB</sub> (plasmid) | S2F |
|  | *iolT1* (genome) | G87S |
| **FAB evo 2** | pPREx2 (plasmid) | c to t mutation 20 bp upstream of *xylA*<sub>AB</sub> start codon (c_t) |
|  | *iolT1* (genome) | T351P |
| **FAB evo 3** | pPREx2 (plasmid) | c to t mutation 20 bp upstream of *xylA*<sub>AB</sub> start codon (c_t) |
|  | *iolT1* (genome) | G87S |
| **FXC evo 1** | *xylA*<sub>XC</sub> (plasmid) | W147S |
|  | *iolT1* (genome) | G87S |
| **FXC evo 2** | *xylA*<sub>XC</sub> (plasmid) | W147S |
|  | *iolT1* (genome) | G87S |
| **FXC evo 3** | *xylA*<sub>XC</sub> (plasmid) | W147S |
|  | *iolT1* (genome) | T351P |

### Reverse engineering to confirm the relevance of the ALE-derived mutations

To confirm the relevance of the individual mutations and the assumption that a combination of *xylA* and *iolT1* mutations is required to achieve fast growth on D-fructose, we performed reverse engineering and introduced the mutations singly or in combination into the parent Fru^neg^ strain carrying either pPREx2-*xylA*_AB_ or pPREx2-*xylA*_XC_. In total, 18 strains carrying either the single mutations *iolT1*-T351P, *iolT1*-G87S, *xylA*_AB_-S2F, *xylA*_AB_(c_t), and *xylA*_XC_-W147S or various combinations were constructed by homologous recombination, checked for correctness, and tested for growth in D-fructose minimal medium. As shown in Figure 2D-G and Figure S2C-F in the supplemental information online, all introduced mutations showed a positive effect on growth, but the mutations in *iolT1*, i.e. T351P and G87S, stimulated growth and D-fructose consumption more strongly than the different *xylA* mutations. While pPREx2-*xylA* related mutations increased the growth rate maximally twofold, the G87S or T351P mutations in *iolT1* enhanced the growth rate by three- to sevenfold. However, the combination of *iolT1* and *xylA* mutations was necessary and sufficient to obtain strains with growth characteristics similar to the ALE strains containing the same mutations.

### Effect of the IolT1 mutations on the kinetics of D-fructose and D-glucose transport

The effects of the amino acid exchanges G87S and T351P found in IolT1 were analyzed by determining the growth rate of the Fru^neg^ strain with either wild-type IolT1, IolT1-G87S (IolT1^G87S^), IolT1-T351P (IolT1^T351P^), or IolT1-G87S-T351P (IolT1^G87S-T351P^) at varying concentrations of D-fructose. The concentration enabling the half-maximal growth rate represents the K_s_ value. For this experiment, the strains were transformed with plasmid pPREx2-*scrK* that encodes a fructokinase from *Clostridium acetobutylicum* known to be active in *C. glutamicum* [32,33]. With this enzyme, D-fructose taken up by IolT1 is phosphorylated to D-fructose 6-phosphate enabling growth of the cells in D-fructose minimal medium [20] independent of D-glucose isomerase activity.

Strain Fru^neg^ pPREx2-*scrK* with wild-type IolT1 grew poorly in D-fructose minimal medium with a maximal growth rate (µ_max_) of 0.032 ± 0.002 h^-1^ and a K_S_ value of 106.9 ± 7.6 mM (Figure 3, Figure S3 in the supplemental information online, Table 2). The strains with the variants IolT1^G87S^, IolT1^T351P^, and IolT1^G87S-T351P^ showed an about 10-fold increased growth rate with K_S_ values reduced by 10-32% (Table 2). These results confirmed that both mutations found in IolT1 after ALE strongly improve its ability for D-fructose transport. A combination of both mutations improved µ_max_ slightly, but concomitantly increased the K_S_ value. Hence, the double mutation did not show an additive effect and the two single mutations are sufficient to boost D-fructose uptake.

**Figure 3.**
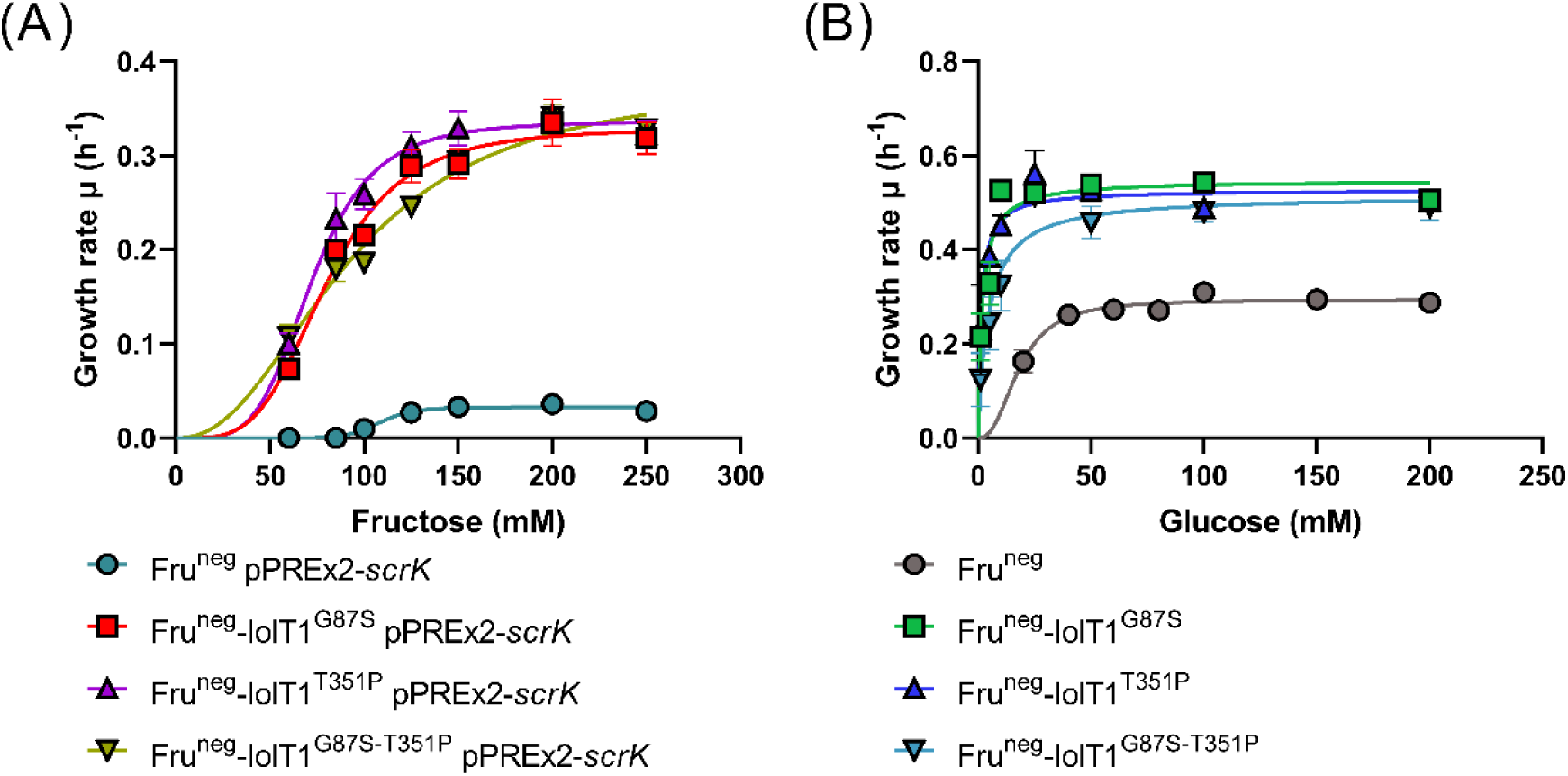
Effects of the mutations G87S and T351P in IolT1 on growth with D-fructose and D-glucose as substrates. (A) Growth experiments in D-fructose minimal medium were performed with Fru^neg^ strains containing the pPREx2-*scrK* plasmid, expressing fructokinase from *C. acetobutylicum*. The minimal medium was additionally supplemented with 1 mM IPTG and 25 µg/mL kanamycin. (B) Growth experiments in D-glucose minimal medium were performed with Fru^neg^ strains without a plasmid and minimal medium without IPTG and kanamycin. All experiments were performed in a BioLector XT at 30°C and 1200 rpm for 24-48 h. Inoculation was performed to an initial OD_600_ of 5. All data points given represent the mean ± S.D. of three biological replicates (*n* = 3). The data was fitted in GraphPad Prism using the default sigmoidal non-linear regression fit.

**Table 2.** Influence of the amino acid exchanges G87S and T351P in IolT1 on µ_max_ and K_s_.

| Substrate | Strain | $\mu_{\max}$ (h<sup>-1</sup>)<sup>a</sup> | $K_s$ (mM)<sup>a</sup> |
| --- | --- | --- | --- |
| **D-Fructose** | Fru<sup>neg</sup> pPREx2-*scrK* | 0.032 ± 0.002 | 106.9 ± 7.6 |
|  | Fru<sup>neg</sup>-IolT1<sup>G87S</sup> pPREx2-*scrK* | 0.329 ± 0.017 | 80.2 ± 1.8 |
|  | Fru<sup>neg</sup>-IolT1<sup>T351P</sup> pPREx2-*scrK* | 0.336 ± 0.013 | 72.6 ± 4.2 |
|  | Fru<sup>neg</sup>-IolT1<sup>G87S-T351P</sup> pPREx2-*scrK* | 0.380 ± 0.003 | 93.3 ± 3.3 |
| **D-Glucose** | Fru<sup>neg</sup> | 0.296 ± 0.010 | 18.9 ± 2.8 |
|  | Fru<sup>neg</sup>-IolT1<sup>G87S</sup> | 0.548 ± 0.017 | 1.9 ± 0.8 |
|  | Fru<sup>neg</sup>-IolT1<sup>T351P</sup> | 0.532 ± 0.010 | 1.4 ± 0.9 |
|  | Fru<sup>neg</sup>-IolT1<sup>G87S-T351P</sup> | 0.519 ± 0.039 | 4.9 ± 1.5 |

<sup>a</sup> Mean ± S.D. of the sigmoidal non-linear regression fits of three independent replicates

Besides D-fructose, we also tested the growth rate dependency on the D-glucose concentration. In this case, no plasmid was required due to the availability of the endogenous glucokinases Glk and PpgK [27,28]. Compared to D-fructose, the Fru^neg^ strain with wild-type IolT1 reached an almost ten-times higher µ_max_ of 0.296 ± 0.010 h^-1^ and a six-times lower K_S_ value of 18.9 ± 2.8 mM (Figure 3, Figure S3 in the supplemental information online, Table 2), revealing a clear preference of IolT1 for D-glucose over D-fructose. The strains with the variants IolT1^G87S^, IolT1^T351P^, and IolT1^G87S-T351P^ almost doubled their µ_max_ values, while revealing strongly reduced K_S_ values by factors of 4 - 14 (Table 2). As in the case of D-fructose, the combined mutations did not improve D-glucose transport. In summary, these experiments confirmed that the amino acid exchanges G87S and T351P in IolT1 have a strongly positive effect on D-fructose and D-glucose transport, presumably leading to increased cytoplasmic concentrations of these sugars, thereby stimulating their conversion by D-glucose isomerase.

### Molecular dynamics simulations of the IolT1 variants

To assess the influence of the G87S and T351P substitutions on the structural stability of IolT1, we performed all-atom unbiased MD simulations of structural models of IolT1 in four different configurations (outward-open, outward-occluded, inward-occluded, inward-open), in part bound to D-glucose (as β-D-glucopyranose, BGP) and D-fructose (as β-D-fructofuranose, BFF), inserted into the lipid bilayer according to the OPM (Orientations of Proteins in Membranes) database (Figure S4-10 and Table S2 in the supplemental information online) [34]. Starting structures of the IolT1 variants were generated using the FoldX suite [35]. In all cases, the change in the folding free energy ΔΔ*G* upon the G87S and T351 substitutions is <3 kcal/mol, indicating that they are not considered destabilizing [36]. The membrane composition used was phosphatidylglycerol 16:0-18:1, cardiolipin 16:0-18.1, and phosphatidylinositol 16:0-18:1 in a ratio of 6:2:1, resembling the composition of the inner membrane of *C. glutamicum* [37]. For each configuration, five replicas were simulated, each for 2.0 µs in length, of which the last 1.5 µs were used for analysis. We used the generated conformations to probe if substitutions G87S, T351P, and G87S-T351P change the structural stability of IolT1 by applying a model of dynamic allostery [38,39] previously used on receptors and transporters [40,41]. The model describes allosteric effects due to substitutions in terms of the per-residue (i, eq. 2, Supplemental Methods) or per-region (r, eq. 3, Supplemental Methods) rigidity index Δr_i/r,(Mutant – WT)_ [39]. We evaluated Δr_i/r,(Mutant – WT)_ for the helices 290-306 and 390-401, which are involved in the binding of the carbohydrate ligand in the outward-occluded and inward-occluded configurations and are critical for the carbohydrate transport to the inside of the cell, as found for the human GLUT transporter [42]. There, the helices are named TM7b and TM10b. Conformational changes of these helices during the transport cycle are common in the major facilitator superfamily (MFS) [43] and accompany the rocker-switch mechanism used by MFS transporters.

The helices 290-306 and 390-401 of IolT1 are more than 20 Å and 12 Å away from the substitutions G87S and T351P, respectively (Figure 4A). The substitutions are also not located near other parts of the transport route, so that a trivial effect on the transport process is unlikely. The model of dynamic allostery revealed a configuration-, substitution type-, and carbohydrate-dependent impact of the amino acid substitutions on the structural stability of both helices 290-306 and 390-401. First, the substitutions destabilize the helices in the outward-open configuration, whereas they stabilize them in the inward-open configuration (Figure 4A-B). In between, in the outward-occluded configuration, helix 290-306 becomes destabilized and helix 390-401 stabilized; in turn, in the inward-occluded configuration, helix 290-306 is stabilized and helix 390-401 is destabilized. In either case, the destabilization or stabilization is less than in the outward-open or inward-open configurations, respectively. Second, these effects decrease in the order of the substitutions T351P > G87S > G87S-T351P, i.e., in correlation with the substitution effects on the growth rate (Figure 3). Note that substitution effects on structural stability are not necessarily additive, which can explain why the impact of the double substitution is lower than that of the single ones [44]. Finally, the (de-)stabilizing effects for the two occluded states are generally weaker in the case of bound D-fructose than D-glucose (Figure 4B), again in line with the difference in growth rates between the two carbohydrates (Figure 3). Compared to wild-type IolT1, this leads to a succession of configurations with increasing structural stability along the transport cycle (Figure 4B), which favors the inward-open configuration as the most structurally stable. These changes could result in the enhanced transport of the cargo from the outside to the inside of the cell.

**Figure 4.**
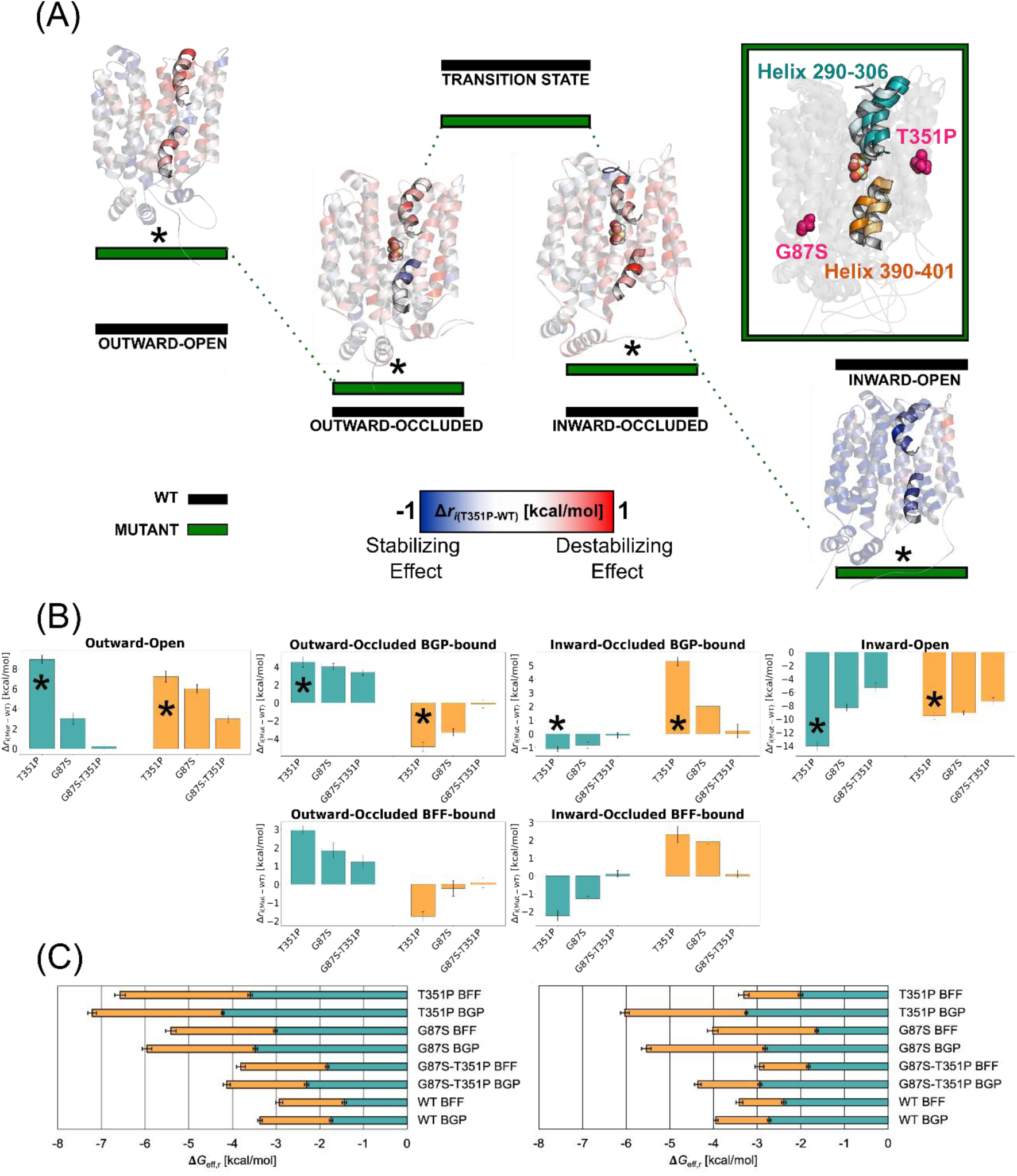
IolT1 mutations impacting D-glucose (β-D-glucopyranose, BGP) and D-fructose (β-D-fructofuranose, BFF) transport exert an allosteric effect on two helices critical to MSF transporter function. (A) IolT1-mediated transport is modeled using the induced transition fit (ITF) mechanism, where substrate binding supplies energy for conformational changes, resembling substrate destabilization in enzyme catalysis. The wild-type IolT1 energy profile (black bars, [42]) and T351P mutation effects (green bars) show how T351P destabilizes helices 290-306 and 390-401 in the outward-open state and stabilizes them in the inward-open state, facilitating substrate transport into the cell. The structural overlay of IolT1 highlights helix 290-306 (turquoise) and helix 390-401 (orange), with a gradient from outward- to inward-open states and the substrate (D-glucose), showing substitution sites G87S and T351P (magenta). (B) The difference Δr_r,(Mutant-WT)_ (eq. 3, Supplemental Methods) across residues of helices 290-306 and 390-401 reveals that T351P has the greatest stabilizing effect, followed by G87S and G87S-T351P, correlating directly with cell growth trends. This stabilization effect is more pronounced with D-glucose than D-fructose. The asterisks (*) indicate the values depicted in panel (A) as per-residue contributions. Error bars indicate the SEM. (C) Δ*G*_eff,r_ sums per-residue contributions to effective binding energy for helices 290-306 and 390-401, in outward- (left) and inward-occluded (right) states, averaged from five replicas per variant. Error bars represent the propagated SEM and, thus, indicate the precision, not the accuracy, of the method. For all variants, D-glucose binding shows more favorable binding, reinforcing the trend from panel (B).

While the gradient-driven transport is initiated by the binding of the cargo to the outward-open configuration [45], the reset of the transporter from the inward-open to the outward-open configuration is known for other carbohydrate transporters, such as the lactose permease of *E. coli* [46,47], to involve interactions of lipid head groups with charged amino acids to accommodate the energetic barrier imposed by salt bridges in the inward-open configuration. In IolT1, after the release of the carbohydrate on the intracellular side, D46 in TM1 and E400 in TM10b are thought to trigger the resetting between inward- and outward-facing conformations similarly to what has been suggested in other carbohydrate transporters [46,47]. MD simulations of IolT1 revealed that the PG lipid headgroups interact with D46 and E400, which form salt bridges to R126 (100.0% of total frames) and R339 (75.9% of total frames), respectively (Figure S11-12 in the supplemental information online). The substitutions do not affect such salt bridges (Figure S11-12 in the supplemental information online), thus, they should not influence the reset process.

To explain the difference in transport between D-glucose and D-fructose, we assessed the impact of helices 290-306 and 390-401 on carbohydrate binding in the outward-occluded and inward-occluded states by computing residue-wise effective binding energies. D-Glucose was found to bind stronger than D-fructose, in both the wild-type protein and the variants, and irrespective of the IolT1 configuration (Figure 4C). A less favorable binding of D-fructose likely impacts the transport rate of the carbohydrate. The IolT1 variants revealed stronger interactions with the carbohydrates than wild-type IolT1, suggesting that allosteric effects due to the substitutions can improve the binding of the carbohydrates in addition to impacting the structural stability of the transporter (Figure 4C). To conclude, substitutions T351P and G87S allosterically impact functionally relevant helices of IolT1, and the impacts correlate with the increase of the growth rates enabled by these substitutions.

### Effects of mutations in XylA on its kinetic properties

ALE-mediated selection for faster growth on D-fructose resulted in three different mutations with respect to the *xylA* gene, i.e. *xylA*_AB_-S2F, *xylA*_AB_(c_t), and *xylA*_XC_-W147S (Table 1). Mutations in the 5’-untranslated region (5’-UTR) and at the second codon of *xylA*_AB_ can influence gene expression levels, e.g. by stabilization of the mRNA or enhancing translation initiation [48–50]. To test the effect of the S2F and c_t mutations on the XylA protein level, Western blot analysis was performed. Both mutations led to a much higher protein content compared to wild-type XylA_AB_ (Figure S13A in the supplemental information online), suggesting that the improved growth of strains carrying these variants is caused by higher D-glucose isomerase activity due to strongly increased XylA protein levels.

In the case of strain Fru^neg^ carrying pPREx2-*xylA*_XC_, three independent ALE experiments all led to the amino acid exchange W147S in XylA_XC._ This mutation had no obvious effect on the protein level compared to the native enzyme when analyzed by Western blotting (Figure S13B in the supplemental information online). Since the presence of XylA_XC_-W147S in addition to IolT1^G87S^ or IolT1^T351P^ in strain Fru^neg^ led to accelerated growth in D-fructose minimal medium (Figure 2D-G, Figure 2 in the supplemental information online), the W147S mutation should have a positive effect on the D-glucose isomerase activity of XylA. To test this assumption, His-tagged variants of XylA_XC_ and XylA_XC_-W147S were overproduced in *C. glutamicum* MB001(DE3) using plasmids pPREx5-*xylA*_XC_ and pPREx5-*xylA*_XC_-W147S and purified by Ni-NTA and size exclusion chromatography. Both proteins were purified as homodimers (Figure S14 in the supplemental information online). Enzymatic activity was determined by a discontinuous colorimetric assay for D-glucose and D-xylose as substrates [51]. Native XylA_XC_ revealed a 65-fold higher activity for D-xylose compared to D-glucose with catalytic efficiencies (*k*_cat_/K_M_) of 45.2 ± 6.7 s^-1^M^-1^ and 0.7 ± 0.1 s^-1^M^-1^, respectively (Table 3). For the XylA_XC_-W147S enzyme, the *k*_cat_/K_M_ for D-glucose increased by 8.4-fold while the *k*_cat_/K_M_ for D-xylose decreased by 2.6-fold, resulting in a diminished activity for D-xylose, only about three times as high as the activity for D-glucose (Table 3). The results confirmed the strongly positive effect of the W147S mutation on the D-glucose isomerase activity of XylA_XC_.

**Table 3.**
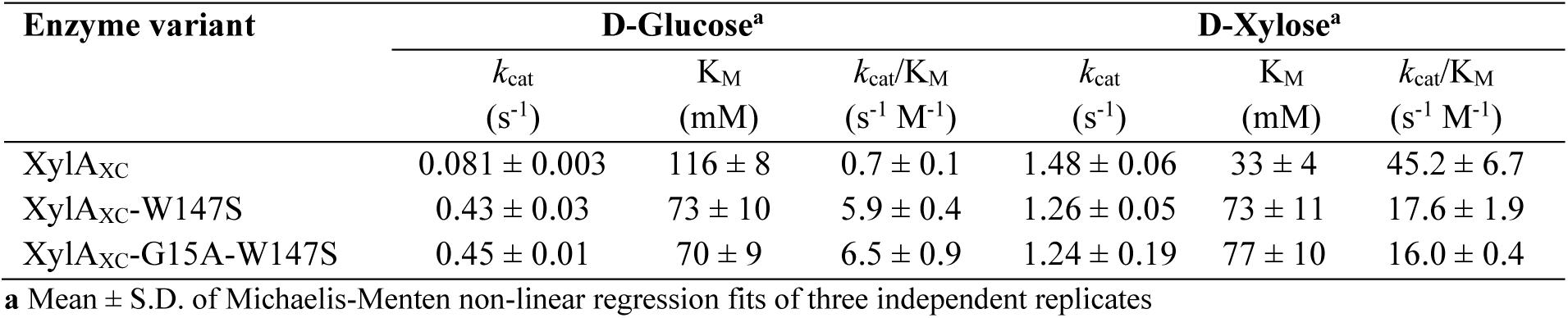
Kinetic parameters of XylA_XC_ variants for the conversion of D-glucose and D-xylose.

To assess whether the catalytic performance of XylA_XC_-W147S could be further improved, we established a modified selection system (Figure S15 in the supplemental information online). It employed the Fru^neg^ strain with the *iolT1*-G87S mutation, thereby alleviating the bottleneck in D-fructose uptake. To enable further optimization of XylA_XC_-W147S activity, expression of the corresponding gene was intentionally limited. For this, we examined the growth of Fru^neg^-IolT1^G87S^ harboring pPREx2-*xylA*_XC_-W147S in D-fructose minimal medium across a range of IPTG concentrations. Plotting the growth rate as a function of the IPTG concentration revealed a Michaelis-Menten-like relationship, with a K_s_ value of 19.7 ± 2.4 µM IPTG enabling a half-maximal growth rate (Figure S16 in the supplemental information online). Subsequently, *xylA*_XC_-W147S was subjected to random mutagenesis by error-prone PCR and the resulting mutant library was introduced into the Fru^neg^-IolT1^G87S^ strain (for details see Supplemental Note 1). In consecutive screenings (Figure S17 and Table S3 in the supplemental information online), a XylA_XC_-W147S variant with an additional G15A mutation was found to further increase the growth rate of Fru^neg^- IolT1^G87S^ in D-fructose minimal medium. Western blot analysis suggested somewhat higher protein levels of the XylA_XC_-G15A-W147S variant compared to the XylA_XC_-W147S variant (Figure S18 in the supplemental information online). Kinetic analysis of purified XylA_XC_-G15A-W147S revealed a slight improvement (10.2%) of catalytic efficiency for D-glucose compared to XylA_XC_-W147S (Table 3).

To explore the structural effects of the W147S and G15A-W147S mutations in XylA_XC_ on the binding of D-glucose, we performed all-atom unbiased MD simulations (for further details see Supplemental Note 2). Computations of effective binding energy revealed improved D-glucose binding caused by the W147S mutation in XylA_XC_ (Figure 5A-B), but did not show considerable changes with the additional G15A mutation (Figure 5C).

**Figure 5.**
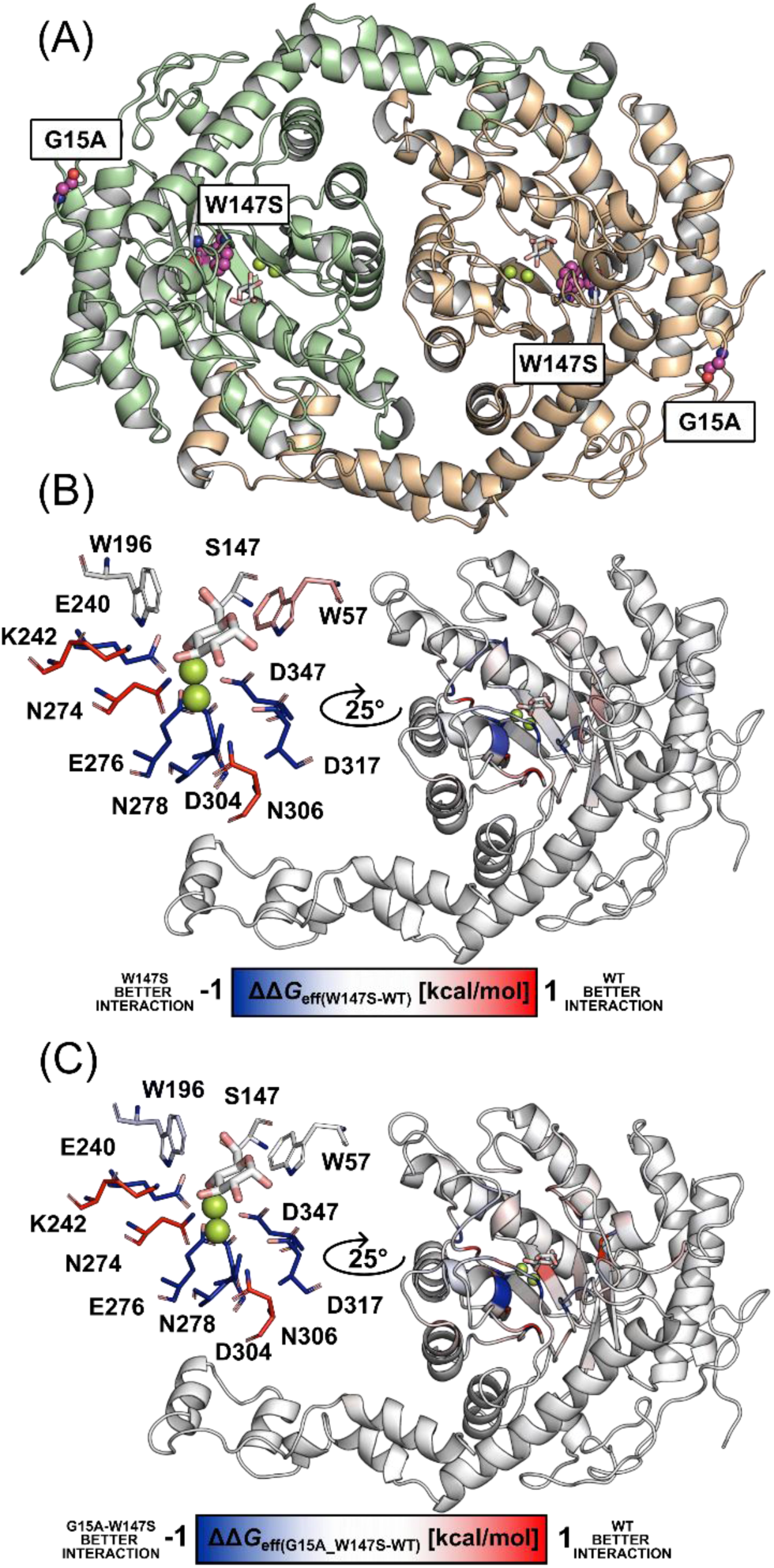
Mutation W147S impacts D-glucose (β-D-glucopyranose, BGP)-Mg^2+^ binding to the XylA homodimer. (A) The homodimeric structure of *X. campestris* XylA_XC_ was predicted using AlphaFold3 [113]. The structure contains D-glucose-Mg^2+^ in the respective binding site of each monomer (green and orange ribbons). D-Glucose is depicted as white-salmon sticks and Mg^2+^ as lemon spheres. The mutation sites G15A and W147S are shown as purple spheres. (B) Differential per-residue effective binding energy (ΔΔ*G*_eff(W147S-WT)_) mapped at the structural level. The energies were averaged across replicas. Larger changes in contributions to the effective binding energy occur near the D-glucose-Mg^2+^ binding site but encompass residues up to 15 Å away from position 147. Residues for which the largest differences were found are depicted as sticks and colored according to the color scale; red and blue colors indicate stronger binding by wild-type XylA_XC_ and the W147S variant, respectively. **c** Differential per-residue effective binding energy (ΔΔ*G*_eff(G15A-W147S-WT)_) mapped at the structural level. The energies were averaged across replicas. Residues for which the largest differences were found are depicted as sticks and colored according to the color scale; red and blue colors indicate stronger binding by wild-type XylA_XC_ and the G15A-W147S variant, respectively.

### Production of D-allulose from D-glucose with *C. glutamicum*

An efficient microbial conversion of D-glucose to D-allulose can only be realized if D-glucose utilization is abolished. For this purpose, the *C. glutamicum* Fru^neg^ strain was further engineered into a D-glucose-negative strain, which cannot utilize D-glucose as sole carbon source. This was achieved by preventing the phosphorylation of D-glucose *via* deletion of the glucokinase genes *glk*, *ppgK*, and *nanK* and by preventing the oxidation of D-glucose to D-gluconate by a side reaction of inositol dehydrogenases via deletion of the inositol-related gene clusters *iol1* and *iol2* and of *idhA3* (for further details see Supplemental Note 3). The resulting Glu^neg^ strain showed no growth in D-glucose minimal medium (Figure S19 in the supplemental information online).

To convert D-glucose to D-allulose based on the Glu^neg^ strain, a suitable D-allulose 3-epimerase was required for the epimerization of D-fructose to D-allulose. The D-allulose 3-epimerases of *Agrobacterium tumefaciens* (Dae_AT_) and *Arthrobacter globiformis* (Dae_AG_) [52,53] and a D-tagatose 3-epimerase of *Pseudomonas cichorii* (Dte_PC_) [54] were selected, since they showed a high residual activity for D-fructose at 30°C. The corresponding genes were cloned into the expression plasmid pPREx2. The enzymes were initially tested in the Fru^neg^ strain to check for D-allulose formation from D-fructose in a medium containing 20 g/L of D-fructose and 20 g/L of D-glucose, the latter as growth substrate. Dae_AT_ was the most efficient in producing D-allulose from D-fructose, giving a titer of 4.18 ± 0.14 g/L D-allulose corresponding to a yield of 22.3 ± 1.1 % (Figure S20 in the supplemental information online) and therefore was choosen for the subsequent studies.

For D-allulose synthesis from D-glucose, the plasmids pPREx2-*xylA*_XC_-*dae*_AT_, pPREx2-*xylA*_XC_-W147S-*dae*_AT_, and pPREx2-*xylA*_XC_-G15A-W147S-*dae*_AT_ were constructed, which enable simultaneous synthesis of D-glucose isomerase and D-allulose 3-epimerase. After transfer into the Glu^neg^ strain, the recombinants were cultivated in minimal medium with 10 g/L of D-gluconate for biomass formation and 30 g/L of D-glucose for D-allulose formation. With native XylA_XC_ and Dae_AT_ an D-allulose titer of 1.32 ± 0.07 g/L was obtained after 72 h, corresponding to a yield of 4.1 ± 0.3 % from D-glucose (Figure 6). With the improved variants XylA_XC_-W147S and XylA_XC_-G15A-W147S, the yield increased to 9.6 ± 0.4 % and 10.2 ± 0.2 %, respectively (Figure 6). In the next step, D-glucose uptake was improved by introducing the G87S mutation into the genomic *iolT1* gene of strain Glu^neg^ resulting in strain Glu^neg^-IolT1^G87S^. When transformed with pPREx2-*xylA*_XC_-G15A-W147S-*dae*_AT_, this strain produced 3.45 ± 0.14 g/L D-allulose, corresponding to a yield of 10.9 ± 0.4 % (Figure 6).

**Figure 6.**
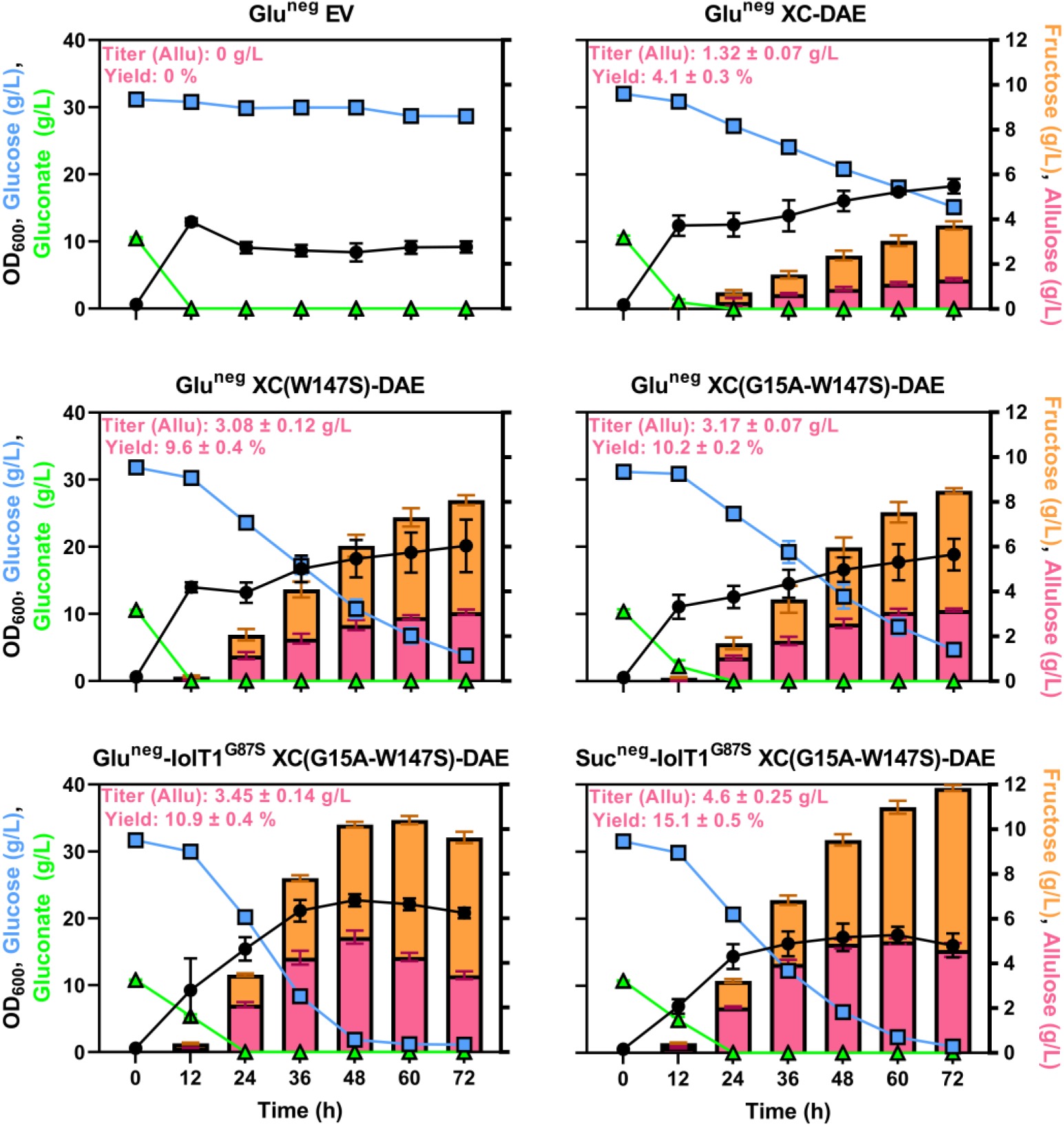
Production of D-allulose from D-glucose by *C. glutamicum* Glu^neg^ and Glu^neg^-IolT1^G87S^. The strains contained either pPREx2-*xylA*_XC_-*dae*_AT_ (XC-Dae_AT_), pPREx2-*xylA*_XC_-W147S-*dae*_AT_ (XC-W147S-Dae_AT_), pPREx2-*xylA*_XC_-G15A-W147S-*dae*_AT_ (XC-G15A-W147S-Dae_AT_) or pPREx2 (EV). The experiment was conducted in CGXII medium with 10 g/L D-gluconate and 30 g/L D-glucose, 25 µg/mL kanamycin and 1 mM IPTG at 30°C and 130 rpm. Samples of the supernatant were taken every 12 h for a total time of 72 h. Growth was measured via OD_600_ and sugar concentrations were determined via HPLC. The data points represent average values with standard deviations of biological triplicates. The data shown represents the mean ± S.D. from three biological replicates (*n* = 3) or from two biological replicates (*n* = 2) for the 0 h sugar concentrations of Suc^neg^-IolT1^G87S^.

Unexpectedly, during the D-allulose production experiment an increase of the OD_600_ concomitant with accelerated consumption of D-glucose was observed for the producer strains in comparison to the empty vector control (Figure 6), which required further attention. The presence of D-glucose isomerase in strain Glu^neg^ or Glu^neg^-IolT1^G87S^ was shown to be necessary for this behavior (Figure S21-22 in the supplemental information online), suggesting that the simultaneous presence of D-glucose and D-fructose could enable partial utilization for growth, despite the Glu^neg^ strain background. To elucidate the reason for this behavior, strain Glu^neg^-IolT1^G87S^ with pPREx2-*xylA*_XC_-W147S was subjected to ALE to obtain clones showing faster growth on D-glucose. In the ALE experiments, the cultures were initially grown with 10 g/L D-gluconate as second carbon source besides 30 g/L D-glucose. Later the D-gluconate content was reduced to 2 g/L and eventually completely omitted (Figure S23A in the supplemental information online). Fast growing clones were obtained in each of two biological replicates (Figure S23B-C in the supplemental information online). Genome sequencing of one clone from each experiment revealed that both carried an identical mutation in the *ptsS* gene, causing the amino acid exchange A129G in the EII transporter PtsS for D-sucrose (Table S4 in the supplemental information online). As *C. glutamicum* lacks a gene for D-sucrose synthase that might have formed D-sucrose from NDP-glucose and D-fructose, we speculated that the A129G mutation might reinforce the ability of PtsS to co-transport D-glucose and D-fructose with concomitant formation of D-glucose 6-phosphate which could be used as a growth substrate. Further studies on PtsS-A129G are reported in Supplemental Note 4.

To prevent D-glucose utilization by PtsS in D-allulose production, we deleted the *ptsS* gene in Glu^neg^-IolT1^G87S^, resulting in the strain Suc^neg^-IolT1^G87S^. When transformed with pPREx2-*xylA*_XC_-G15A-W147S-dae_AT_, the absence of PtsS increased the D-allulose titer by 33% compared to the PtsS-positive strain to 4.6 ± 0.25 g/L and the yield by 39% to 15.1 ± 0.5 % (Figure 6). In high-cell-density biotransformations of the Suc^neg^-IolT1^G87S^ strain with pPREx2-*xylA*_XC_-G15A-W147S-dae_AT_ (for details see Supplemental Methods), where 58 g/L of D-glucose was added to the culture after biomass formation from 60 g/L of D-gluconate, a D-allulose yield of 11.4 ± 0.2 % was obtained (Figure S24 in the supplemental information online).

## Discussion

The microbial conversion of D-glucose to D-allulose in foods and beverages provides opportunities to significantly reduce their caloric content and thus the health problems associated with consumption of high-caloric sugars. In our study we created strains of *C. glutamicum* that catalyze the conversion of D-glucose to D-allulose in a two-step reaction involving D-glucose isomerase and D-allulose 3-epimerase. The major hurdle was an efficient conversion of D-glucose to D-fructose within the cytoplasm. D-Glucose isomerases usually have K_m_ values above 100 mM for D-glucose and D-fructose, but the cytoplasmic concentrations in bacteria are much lower since as these sugars are immediately metabolized after uptake. Furthermore, *C. glutamicum* as many other bacteria take up D-glucose and D-fructose via the PTS, which generates the phosphorylated forms of these sugars that do no serve as substrates for D-glucose isomerase. We prevented uptake via the PTS by deleting the EII enzymes for D-glucose, D-fructose, and D-sucrose, and at the same time enabled uptake of D-glucose and D-fructose via the promiscuous transporter IolT1. When expressing D-glucose isomerase genes in this selection strain, a small but distinct growth on D-fructose was observed due to its conversion to D-glucose followed by phosphorylation and degradation. Subjecting these strains to ALE allowed the selection of clones with a strongly enhanced growth rate on D-fructose.

Genomic analysis of six independent ALE clones revealed two features that were responsible for the improved growth rate. Two different single amino acid exchanges in IolT1, G87S and T351P, both strongly improved the uptake of D-glucose and D-fructose. This presumably leads to highly elevated cytoplasmic concentrations of these sugars, which in turn strongly increase D-glucose isomerase activity. Molecular dynamics simulations suggest that both amino acid exchanges allosterically affect two helices known to be functionally involved in transport, thereby enhancing D-glucose and D-fructose transport. The second feature was an increase in the D-glucose isomerase activity. In the case of the enzyme from *Arthrobacter* the elevated activity was due to a strongly increased protein level caused by a mutation in the non-coding mRNA region or by an S2F exchange in the XylA protein. In the case of D-glucose isomerase from *X. campestris*, the increased activity was caused by an W147S exchange, which raised the catalytic efficiency for D-glucose almost 9-fold while reducing it at the same time for D-xylose. Molecular dynamics simulations suggest that binding of D-glucose to XylA_XC_ is stabilized by the W147S substitution. The contribution of the IolT1 mutations to faster growth on D-fructose was higher than the contribution the XylA mutations, but the combination of both mutations was necessary to rebuild strains with the same growth rates as the ALE-derived strains.

Native IolT1 has been used in numerous studies for the development of strains producing lysine [26], serine [55], shikimate [56], 3-hydroxypropionate [57], noreugenin [58], or hyaluronic acid [59]. The identification of the variants IolT1^G87S^ and IolT1^T351P^ enabling about 10-fold increased uptake rates for D-glucose and D-fructose should be of high interest for future metabolic engineering studies using IolT1 as a tool to enhance the overall sugar uptake rate or to avoid using the PTS, which is disadvantageous for the production of aromatic compounds requiring PEP as precursor for the shikimate pathway. D-Xylose isomerase is used in yeast for bioethanol production from D-xylose [60], a major constituent of hemicellulose. To our knowledge, this enzyme has never been used before in microbial production strains for the conversion of D-glucose to D-fructose. The XylA_XC_-W147S variant identified in our work offers new options to microbially synthesize D-fructose-derived products from D-glucose, which include disacharide sweeteners (e.g. kojibiose, cellobiose) [61,62], dietary supplements (e.g. mannose) [63], or even furan-based chemicals (e.g. 5-hydroxymethylfurfural) [64].

Our approaches to engineer a D-glucose-negative strain of *C. glutamicum* turned out to be quite challenging, as suppressor mutants rapidly evolved from strains lacking PtsG and the glucokinases Glk and PpgK. We revealed that inactivating mutations in the genes for the transcriptional repressors IolR and IhtR led to the derepression of their target genes, which include several genes for inositol dehydrogenases[23] that can catalyze the oxidation of D-glucose to D-gluconate as a side-reaction and thereby enable growth. Only after deletion of the inositol dehydrogenase genes a D-glucose-negative strain for D-allulose production was obtained. Many bacteria use D-glucose dehydrogenases for D-glucose metabolization via D-gluconate, such as *Pseudomonas putida* [65] or *Gluconobacter oxydans* [66]. *C. glutamicum* does not possess such dedicated enzymes, but can easily obtain the corresponding activity by inducing inositol dehydrogenases, as shown in our work.

Another surprising observation during our study was that the D-glucose-negative strain regained the ability to grow with D-glucose when it expressed D-glucose isomerase. The molecular basis of this phenotype was disclosed by ALE experiments selecting for faster growth of the strain, which revealed the amino acid exchange A129G in PtsS, the EII transporter for D-sucrose. We therefore speculate that PtsS might also have a weak ability to co-transport of D-glucose and D-fructose instead of D-sucrose, forming D-glucose 6-phosphate which can be used for growth. This ability could be enhanced by the A129G mutation, as supported by molecular dynamics simulations. Further studies are needed to follow this feature of PtsS.

By deleting *ptsS* in the D-glucose-negative strain the utilization of D-glucose for growth in the presence of D-glucose isomerase could be largely avoided, which increased the D-allulose yield to 15%. This value, limited by the reversibility of the D-glucose isomerase and D-allulose 3-epimerase reactions, corresponds to the one obtained with immobilized enzymes at high temperatures of about 60°C, but avoids enzyme purification and immobilization. Furthermore, in high-cell-density biotransformations a comparable D-allulose yield of ∼12% was obtained.

In conclusion, we provide novel microbial catalysts for the efficient one-pot conversion of D-glucose to D-allulose at 30°C, offering new opportunities for reducing the caloric content of foods and beverages containing D-sucrose or HFCS. Future studies will focus on further engineering of our production strain to enable also the conversion of D-sucrose to D-allulose.

## Methods

### Bacterial strains, plasmids and growth conditions

All bacterial strains and plasmids used in this work are listed in Table 4. Cloning was performed with *Escherichia coli* DH5α as host. Cultivation of *E. coli* was performed in lysogeny broth (LB) medium or on LB agar plates at 37°C with 50 µg/mL kanamycin, if cells were transformed with a plasmid. *C. glutamicum* was cultivated in brain heart infusion (BHI) medium or on BHI agar plates at 30°C, supplemented with 25 µg/mL and 15 µg/mL kanamycin if carrying a pPREx2 and pK19*mobsacB* plasmid, respectively.

**Table 4.**
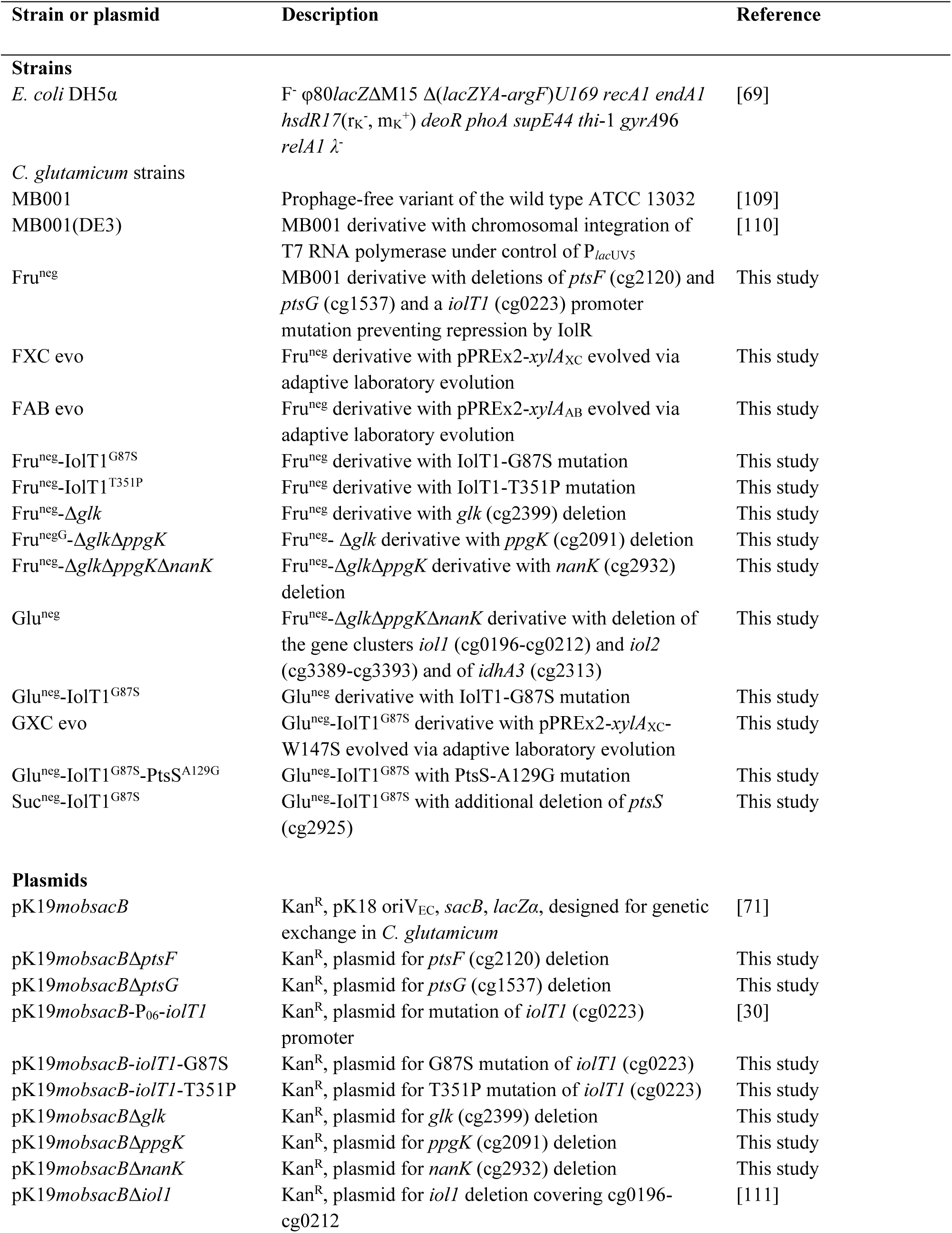
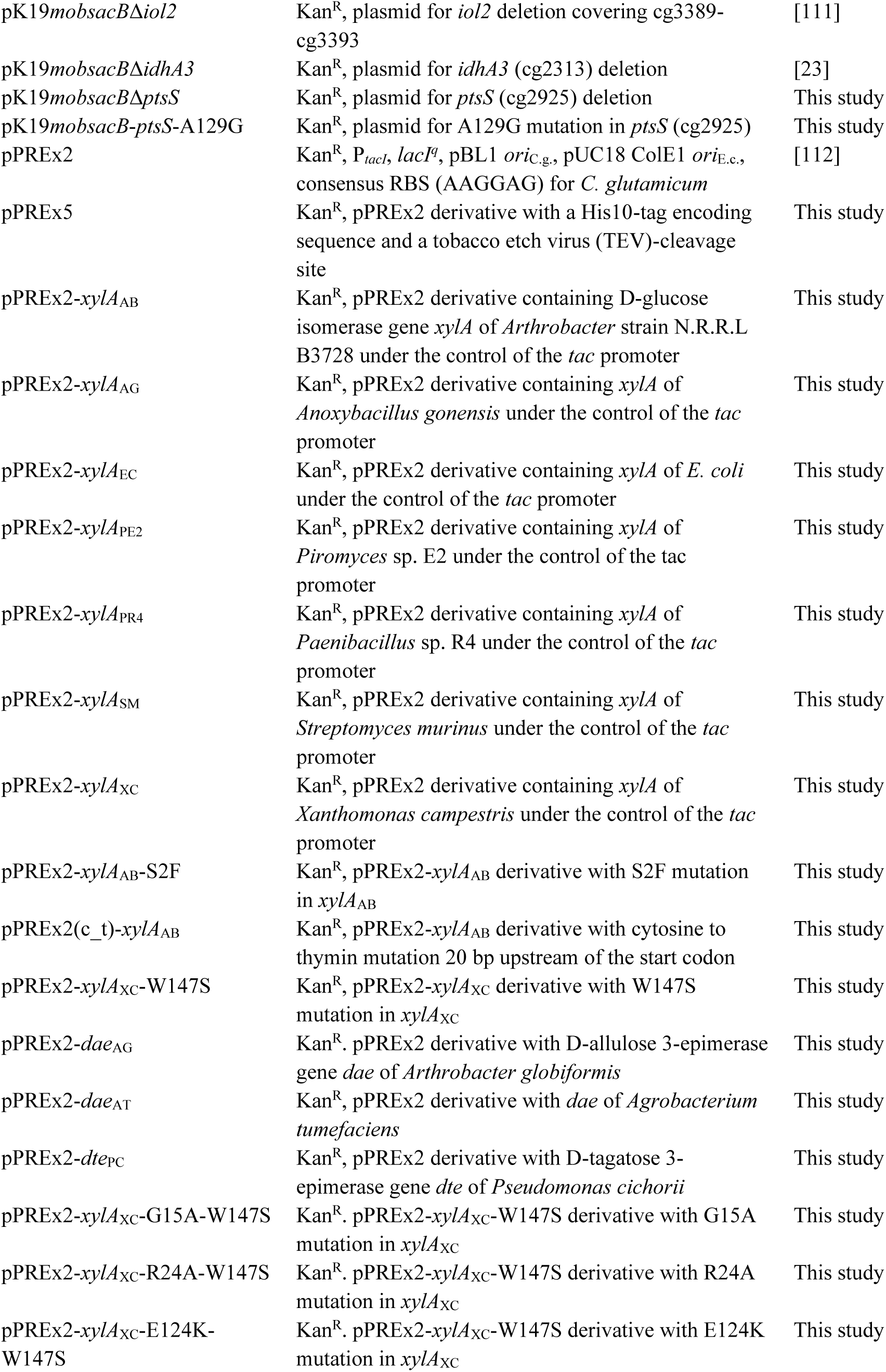
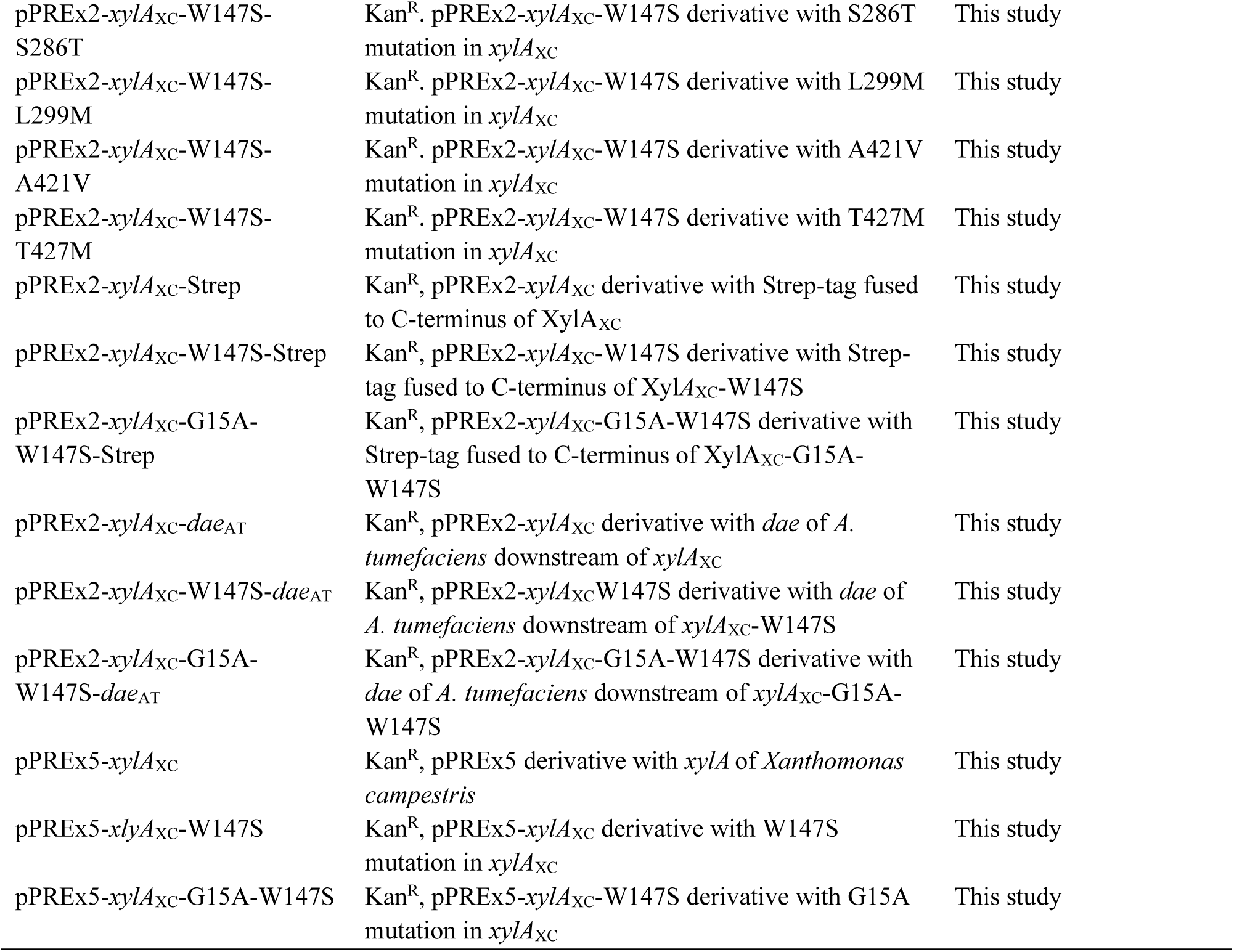
List of strains and plasmids used in this study.

### Recombinant DNA work and construction of deletion mutants

All plasmids and oligonucleotides used in this study are listed in Table 4 and Table S5 in the supplemental information online, respectively. All D-glucose isomerase (*xylA*), D-allulose 3-epimerase (*dae*), and D-tagatose 3-epimerase (*dte*) genes were ordered in a codon-optimized form for *C. glutamicum* from Life Technologies (Carlsbad, USA). Standard cloning procedures such as PCR, DNA restriction and Gibson assembly were performed according to established protocols [67,68]. Transformation of *E. coli* was performed according to a standard heat-shock protocol [69], while transformation of *C. glutamicum* was conducted via electroporation [70]. Deletion of genes or integration of mutations into the genome of *C. glutamicum* were performed via double homologous recombination using pK19*mobsacB*-based plasmids [71]. Confirmation of deletions or mutations was obtained via colony PCR, using oligonucleotides annealing up- and downstream of the targeted genomic area. To verify the successful integration of mutations, the amplified colony PCR product was additionally sequenced. Expression of selected genes in *C. glutamicum* was performed with the vectors pPREx2 or pPREx5 using a *tac* promoter. The gene of interest was cloned downstream of the consensus ribosome binding site (RBS) of *C. glutamicum* of the respective plasmid. Gene expression was induced by addition of 1 mM or 25 µM isopropyl-β-D-thiogalactoside (IPTG).

### Growth experiments in BioLector microcultivation systems

For micro-scale cultivations of *C. glutamicum*, the BioLector I, II or XT instruments (Beckman Coulter, Brea, USA) were used. Growth in this system was measured online as scattered light at 620 nm with a signal gain factor of 2, 4 or 10 with the BioLector I, II or XT, respectively [31]. Cultivations were performed in a 48-well FlowerPlate (Beckman Coulter, Brea, USA) by using defined CGXII medium [72] with 0.03 g/L protocatechuic acid and usually 40 g/L D-glucose, D-fructose or D-sucrose. If the strains carried expression plasmids, 1 mM IPTG and 25 µg/mL kanamycin were added to the medium. Inoculations were individually prepared to reach an initial OD_600_ of 0.5 or 5. BioLector cultivations were performed at 30°C, 1200 rpm and 85% humidity.

### Adaptive laboratory evolution (ALE)

ALE experiments for the evolution of D-glucose isomerases were prepared in 24-well plates (Porvair Sciences, King’s Lynn, UK) containing 5 ml CGXII medium with 40 g/L of D-fructose, 25 µg/mL kanamycin and 1 mM IPTG. The plates were shaken at 30°C and 900 rpm in a Multitron shaker (Infors HT, Einsbach, Germany). For evolution of the D-glucose isomerase of *Xanthomonas campestris* the first two transfers into fresh medium were performed after 5 days of cultivation with an initial OD_600_ of 7.5. Afterwards, the initial OD_600_ was lowered to 5 and the cultivation time was increased to 15 days. After three more consecutive cultivations, the experiment was stopped. For evolution of the D-glucose isomerase of *Arthrobacter* strain N.R.R.L B3728, the ALE was started at an OD_600_ of 5 with a cultivation time of 15 days. After three rounds of cultivation, the experiment was terminated. ALE experiments for the evolution of the Glu^neg^-IolT1-G87S strain were performed in 100 mL baffled shake flasks. Initially, CGXII medium containing 10 g/L D-gluconate and 30 g/L D-glucose was used. The initial OD_600_ was set to 0.5 and the cultivation time to 3 days. After three rounds of cultivation, the medium composition was changed to 2 g/L D-gluconate and 30 g/L D-glucose for one cultivation of 3 days, whereupon the CGXII composition was again changed to solely 30 g/L of D-glucose for the last 3 days of cultivation.

### Analysis of D-glucose isomerase protein levels

Investigation of intracellular protein levels was performed via semi-dry Western blot analysis. For this, C-terminally Strep-tagged protein variants of selected candidates were produced in the Fru^neg^ strain by using pPREx2 plasmid constructs (Table 4). Cultivation of strains was conducted in 100 mL baffled shake flasks containing 10 mL CGXII medium with 10 g/L of D-glucose, 25 µg/mL kanamycin and 1 mM IPTG for 24 h at 30°C and 130 rpm in a Minitron shaker (Infors HT, Einsbach, Germany). Cells were harvested at 4000 *g* for 30 min, resuspended in PBS and lysed via ceramic beads homogenization in the Precellys® 24 (Avantor, Radnor, USA). Cell debris of whole cell lysates was removed via centrifugation at 20.000 *g* for 10 min to obtain soluble protein fractions. 50 µg of each protein fraction was separated via SDS-PAGE and blotted on a nitrocellulose membrane according to standard procedures [67]. After blocking unspecific and biotinylated protein binding to the membrane via a skim milk and avidin solution, respectively, Strep-tagged proteins were then visualized by a Strep-Tactin-HRP conjugate-based assay (IBA Lifesciences, Göttingen, Germany).

### Production and purification of D-glucose isomerase variants

Production of XylA_XC_ variants was achieved by using *C. glutamicum* MB001(DE3) strains carrying pPREx5 derivatives with the desired *xylA*_XC_ variant. BHI medium was used for cultivation, which was inoculated to an OD_600_ of 0.5. Target gene expression was induced with 1 mM IPTG after 4 h of cultivation at 30°C and 130 rpm. After 24 h of cultivation, cells were harvested by centrifugation (4000*g*, 15 min, 4°C) and stored at -80°C for further use. For cell lysis, cells were resuspended in 4 mL lysis buffer (20 mM Tris-HCl pH 7.9, 500 mM NaCl, 5% (v/v) glycerol) per g cell wet weight containing cOmplete EDTA-free protease inhibitor (Roche, Basel, Switzerland) and lysed via the Multi Cycle Cell Disruptor (Constant Systems, Daventry, UK). Cell debris was removed via centrifugation (5000*g*, 30 min, 4°C) and filtration of the supernatant via Filtropur 0.2 µm filters (Sartorius, Göttingen, Germany). The soluble protein fraction was then loaded on a HisTrap HP column (GE Healthcare, Chicago, USA) and His-tagged proteins were eluted with 500 mM imidazole in elution buffer (20 mM Tris-HCl pH 7.9, 500 mM NaCl, 5% (v/v) glycerol, 500 mM imidazole). The resulting His-tagged XylA_XC_ proteins were further purified by size exclusion chromatography using a Superdex 200 10/300 GL column (GE Healthcare, Chicago, USA) via isocratic elution with enzyme assay buffer (200 mM NaPO_4_ pH 7.5, 10 mM MgCl_2_). Protein concentrations were determined with the Bradford assay [73].

### D-Glucose isomerase activity assay

D-Glucose isomerase activity was detected by measuring D-fructose or D-xylulose formed by isomerization of D-glucose or D-xylose, respectively. For this, a molybdosilicate-dependent colorimetric assay was used [51], where the reduction of D-fructose or D-xylulose by Mo(VI) species resulted in a color change from yellow to blue, which was quantified at 750 nm with a Tecan Infinite M200 Pro microplate reader (Männedorf, Switzerland). The staining reaction was initiated by mixing samples obtained from the isomerization reaction with the 20-fold volume of reaction solution (50 mM Na_2_SiO_3_, 600 mM Na_2_MoO_4_, 0.95 M HCl pH 4.5) followed by incubation for 30 min at 70°C. For the isomerization reaction, samples with varying D-glucose (25-1000 mM) and D-xylose (5-750 mM) concentrations were prepared in enzyme assay buffer (200 mM NaPO_4_ pH 7.5, 10 mM MgCl_2_). The reaction was started by adding 12.5-50 µg of purified D-glucose isomerase for D-glucose isomerization and 0.25-10 µg for D-xylose isomerization. Each reaction was conducted in a final volume of 500 µL at 30°C for 24-48 h. Samples were taken after various time points over this period and mixed with 1/5 volume of 1 M HCl to quench the reaction. After the complete reaction was terminated, quenched samples were used for the colorimetric assay.

### D-Allulose production experiments

Production experiments for the conversion of either D-fructose or D-glucose to D-allulose were performed with 25 mL CGXII medium in 500 mL baffled shake flasks, which were incubated at 30°C and 130 rpm over 72 h. For cultivation of Fru^neg^ strains, overnight precultures were prepared in test tubes in BHI and CGXII medium with 20 g/L D-glucose and 25 µg/mL kanamycin. The main culture was prepared in CGXII medium with 20 g/L of D-glucose and 20 g/L of D-fructose, 1 mM IPTG and 25 µg/mL kanamycin. For Glu^neg^ strains, preparation of precultures was similar to Fru^neg^ strains, except that 20 g/L D-glucose was exchanged by 20 g/L D-gluconate in BHI and CGXII medium. Main cultures were prepared in CGXII medium with 10 or 15 g/L of D-gluconate and 30 g/L of D-glucose, 1 mM IPTG and 25 µg/mL kanamycin. In each case, cultivations were started by inoculating the medium to an initial OD_600_ of 0.5. Samples of the supernatant were taken after 12-24 h, used for OD_600_ measurement, filtered with 0.2 µm Whatman Puradisc 13 (Cytiva, Marlborough, USA) and stored at -20 °C for further usage.

### Metabolite quantification via HPLC

Filtered samples taken from the production experiments were diluted 1:4 or 1:8 with deionized water and used for HPLC analysis. Sugars such as D-fructose, D-glucose, and D-allulose were separated and quantified using a Carbo-Pb Guard Cartridge (Phenomenex, Aschaffenburg, Germany) and a Metab-Pb 250 × 7.8 mm column (ISERA, Düren, Germany) in an Agilent LC-1100 system (Agilent Technologies, Santa Clara, USA). For each analysis, 5 µL of sample was injected into the system and separated at 80°C for 45 min with a flow rate of 0.6 mL/min with double-distilled water filtered through a 0.2 µm filter. A refraction index detector operating at 35°C was used for the detection of sugars. Sugar concentrations of samples were measured via a calibration curve, obtained from sugar standards with concentrations of 1 g/L, 2.5 g/L, 5 g/L, and 10 g/L of D-glucose, D-fructose and D-allulose. Organic acids such as D-gluconate, acetate or L-lactate were separated and quantified using a Carbo-H Guard Cartridge and a Rezex ROA – Organic acid H^+^ 300 × 7.8 mm column (Phenomenex, Aschaffenburg, Germany) in an Agilent LC-1260 system. 10 µL of sample were injected into the system for each analysis and separated at 70°C for 40 min with a flow rate of 0.5 mL/min with 5 mM of sulfuric acid. Organic acids were detected via an Agilent Diode Array Detector at 215 nm. Concentration of organic acids were quantified via a calibration curve, obtained from standards with concentrations of 1 g/L, 2.5 g/L, 5 g/L and 10 g/L of D-gluconate, acetate and L-lactate.

### Preparation of IolT1 starting structures

To our knowledge, there is no experimentally resolved 3D structure of IolT1 from *C. glutamicum* available. The starting structures for MD simulations of the different IolT1 states were thus modeled using TopModel [74] and the structures available in the Protein Data Bank [75] with PDB IDs 4ZWC, 4ZW9, 4JA3, and 4YB9 of the GLUT3 transporters in *H. sapiens*, the XylE transporter in *E. coli*, and the bovine GLUT5 transporter [42] as single templates for the outward-open, outward-occluded, inward-occluded, and inward-open conformations, respectively. The target sequence for IolT1 has Uniprot ID Q8NTX0 [76]. The quality of the models was assessed with TopScoreSingle (Table S2 in the supplemental information online) [77].

The substitutions G87S, T351P, and G87S-T351P were introduced into IolT1 using FoldX [35], and the stability of the variants was evaluated in terms of the change in the folding free energy (ΔΔ*G*) with respect to the wildtype [78], as done for other bacterial membrane proteins [36]. Single amino acid substitutions were performed 10 times for the respective residue and the results were averaged. If the average ΔΔ*G* > 3 kcal/mol, the substitution is considered destabilizing [79] and is not further pursued (Table S2 in the supplemental information online).

In an aqueous solution of 20°C, D-fructose exists as an equilibrium of ∼70% β-D-fructopyranose (BFP), ∼23% of β-D-fructofuranose (BFF), and smaller percentages of the open chain and cyclic α-anomers [80]. Considering the low growth rate of cells containing IolT1 for D-fructose (Figure 3), we hypothesized that this originates from the transport of BFF, which is less prevalent in solution. Accordingly, we prepared the BFF structure starting from its canonical SMILES using the fixpka option in OpenEye [81]. The most favorable conformer was generated using Omega 4.1.1.1 [82] and the flag -maxconfs = 1. BFF docking to IolT1 was performed using Autodock-3.0.5 [83] with the objective function DrugScore [84]. The selected pose represents the most populated cluster.

In an aqueous solution of 20°C, D-glucose exists as an equilibrium of ∼36% α-D-glucopyranose (AGP) and ∼64% β-D-glucopyranose (BGP) [85]. The more prevalent BGP structure is available in the Protein Data Bank (PDB ID 4ZW9) bound to human GLUT3 in the outward-occluded configuration [86]. After the superposition of the available protein structure with our generated model (RMSD = 0.3 Å), the bound pose of BGP was extracted and merged with the obtained model. Parallely, BGP was docked into the model, using the same protocol employed for BFF. The two BGP structures have a ligand dRMSD < 1.0 Å. We decided to use the BGP pose from the superpositioning and merging as the starting configuration.

The partial atomic charges of the two carbohydrate configurations were determined by first computing electrostatic potentials with Gaussian 16 [87] at the HF/6-31G level of theory and then performing a restrained fitting according to the RESP procedure [88,89]. The protonation states of protein residues were adjusted according to pH 7.4 using the Epik routine [90–92] in Maestro [93]. The two unbound (outward-open, inward-open) and four bound (outward-occluded and inward-occluded, both with BGP and BFF) IolT1 conformations were embedded into a membrane composed of phosphatidylglycerol 16:0-18:1, cardiolipin 16:0-18.1, and phosphatidylinositol 16:0-18:1 in a ratio of 6:2:1 using the PPM server [34] and solvated using PACKMOL-Memgen [94,95]. The membrane composition resembles that of the native inner membrane of *C. glutamicum* [37,96]. A distance of at least 15 Å between the protein or membrane in the z-direction and the solvent box boundaries was kept. To obtain a neutral system, counter ions were added that replaced solvent molecules (KCl 0.15 M). A total of 16 independent starting configurations were generated and are listed in Table S2 in the supplemental information online.

### Unbiased molecular dynamics (MD) simulations of IolT1 configurations, XylA_XC_ and PtsS

The GPU particle mesh Ewald implementation from the AMBER23 suite of molecular simulations programs [97] with the ff19SB [98], Lipid21 [99], and the GLYCAM-06j [100] forcefields for the protein, membrane lipids, and carbohydrate ligands, respectively, were used; water molecules and ions were parametrized using the OPC3POL model [101,102] and the Li and Merz 12-6 ions parameters [103–105]. For each configuration in Table S2 and Table S6 in the supplemental information online, five independent MD simulations of 2 µs length were performed. Covalent bonds to hydrogens were constrained with the SHAKE algorithm [106], and the hydrogen masses were repartitioned [107], allowing the use of a time step of 4 fs. Details of the thermalization of the simulation systems are given below. All unbiased MD simulations of IolT1 showed structurally rather invariant protein structures and membrane phases as evidenced by electron density calculations (Figure S4-9 in the supplemental information online). The overall nofit-RMSD of β-D-glucopyranose (BGP) and β-D-fructofuranose (BFF) revealed structurally stable binding poses across different replicas for the outward-occluded and inward-occluded configurations (Figure S10 in the supplemental information online). The unbiased MD simulations of XylA_XC_ yielded structurally rather invariant protein structures and stable binding poses of the ligands (Figure S25 in the supplemental information online). All unbiased MD simulations of PtsS showed structurally rather invariant protein structures and membrane phases as evidenced by electron density calculations (Figure S26-27 in the supplemental information online). The overall nofit-RMSD of BGP, BFF, D-sucrose and BGP/BFF revealed structurally stable binding poses across different replicas for the wild type and A129G mutant configurations (Figure S28 in the supplemental information online).

### MM-PBSA calculations to assess β-D-glucopyranose (BGP)/β-D-fructofuranose (BFF) binding to IolT1, BGP binding to XylA_XC_, and carbohydrate binding to PtsS

MM-PBSA (Molecular Mechanics Poisson-Boltzmann Surface Area) calculations were conducted to determine the effective binding energies (Δ*G*_eff_) of BGP/BFF binding to IolT1, BGP binding to XylA_XC_, and the binding of multiple carbohydrates to PtsS. Note that MM-PBSA computations were performed to provide qualitative insight into the relative effects of mutations on ligand binding and protein stability. For further details on the scope of MM-PBSA computations, see Supplemental Note 2. These calculations were performed for IolT1, XylA_XC_, and PtsS in the wildtype state and the proposed mutants, utilizing the last 1.5 µs of the trajectories from unbiased MD simulations. The first 500 ns were considered as part of the thermalization phase. Ligand binding poses were identified as those exhibiting an RMSD < 1.5 Å compared to the first frame of each respective production run. To compute the average effective binding energy of each ligand with the respective system within the ensemble of the unbiased MD frames, we employed MMPBSA.py [108]. This was done using dielectric constants of 1.0 for the protein and 80.0 for the solvent. For IolT1, the cumulative effective binding energy of two specific regions of interest (Δ*G*_eff,r_ for residues 290-306 and residues 390-401) was also calculated. For XylA_XC_ and PtsS, the effective binding energy computations were performed for the two monomers, and the results were averaged. The reported error bars represent the SEM and, therefore, the precision of the method across production trajectories and replica systems and not its accuracy. The computations were converged, as evidenced by the comparison between the first and second halves of the trajectories (Figure S29-30 in the supplemental information online).

## Data and code availability

The data generated in this study are provided in the manuscript, in the supplemental information, and in the repository researchdata.hhu.de at https://researchdata.hhu.de/handle/entry/205 (https://doi.org/10.25838/d5p-88). The manuscript does not report original code. Any additional data can be obtained from the corresponding author upon reasonable request.

## Supporting information

Supplemental figures, tables, notes, methods, and references

## Acknowledgments

This project was funded by the Bundesministerium für Bildung und Forschung (BMBF) within the project IMPRES-2 (FKZ 031B1054B), grant to M.B., and, in part, by the Deutsche Forschungsgemeinschaft (DFG, German Research Foundation), project 267205415/CRC1208, grant to H.G. (subproject A03), and, in part, by a grant from the Ministry of Innovation, Science, and Research within the framework of the NRW Strategieprojekt BioSC (No. 313/323-400-002 13). We are grateful for computational support from the “Zentrum für Informations und Medientechnologie” at the Heinrich-Heine-Universität Düsseldorf and the computing time provided by the John von Neumann Institute for Computing (NIC) to H.G. on the supercomputer JUWELS at the Jülich Supercomputing Centre (JSC) (user IDs: VSK33, T1SS). We would like to thank Susana Matamouros for critically reading the manuscript.

## Author contributions

A.L. and M.B. designed the study. A.L. generated the wet lab experimental data, while R.G. and H.G. performed the molecular dynamics simulations. C. T. supported the strain constructions and A.W. the HPLC measurements. T. P. evaluated the whole genome sequencing data. M. Ba. contributed to supervision. A.L. and R.G. performed the data work including data curation, formal analysis and statistics and prepared figures and tables. A.L., R.G., H.G., and M.B. drafted the manuscript and were responsible for the final review and editing.

