## Supplemental figures, tables, notes, methods, and references for "Microbial production of the low-caloric sweetener D-allulose from D-glucose by evolutionary engineering"

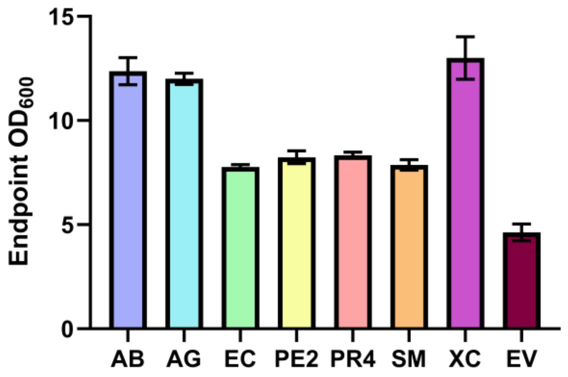

**Figure S1. Endpoint $OD_{600}$ after D-glucose isomerase-mediated growth of $Fru^{neg}$ in D-fructose minimal medium.** $Fru^{neg}$ strains carrying D-glucose isomerase genes from *Arthrobacter* strain N.R.R.L. B3728 (AB), *Anoxybacillus gonensis* (AG), *Escherichia coli* (EC), *Piromyces* sp. E2 (PE2), *Paenibacillus* sp. R4 (PR4), *Streptomyces murinus* (SM), or *Xanthomonas campestris* (XC) or the pPREx2 control (EV). The cultivation was performed in CGXII medium with 40 g/L D-fructose, 25 µg/mL kanamycin and 1 mM IPTG at 30°C and 1200 rpm for 160 h in the BioLector I, after which the endpoint $OD_{600}$ values were determined. All data points given represent the mean ± S.D. from three biological replicates (*n* = 3).

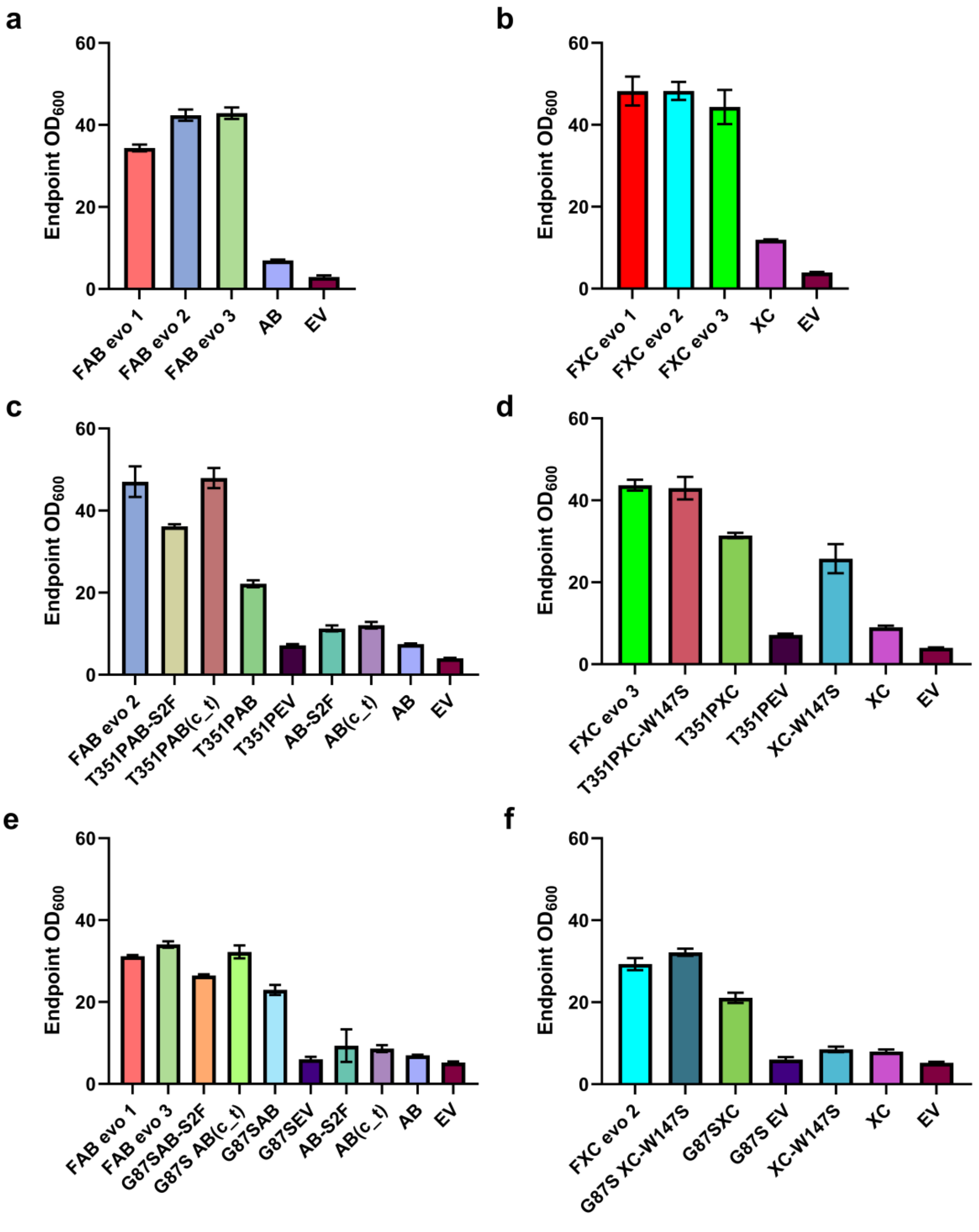

**Figure S2. Endpoint $OD_{600}$ values of reverse-engineered strains with the T351P or G87S mutation introduced into *iolT1*. a**, **b** $OD_{600}$ of $Fru^{neg}$ strains carrying pPREx2-*xylA*$_{AB}$ (FAB evo) or pPREx2-*xylA*$_{XC}$ (FXC evo) in comparison to their unevolved controls (AB, XC) and the pPREx2 control (EV). **c**, **d** $OD_{600}$ values of reverse engineered AB and XC strains, containing *iolT1*-T351P (T351P) with or without pPREx2-*xylA*$_{AB}$-S2F (AB-S2F), pPREx2-*xylA*$_{AB}$(c_t) (AB(c_t)) or pPREx2-*xylA*$_{XC}$-W147S (XC-W147S). **e**, **f** $OD_{600}$ values of reverse engineered AB and XC strains, containing *iolT1*-G87S (G87S) with or without AB-S2F, AB(c_t), XC-W147S. All growth experiments were conducted in CGXII medium with 40 g/L of D-fructose, 25 µg/mL kanamycin and 1 mM IPTG at 30°C and 1200 rpm for ~100 h either in BioLector I or II. The data shown represents the mean ± S.D. from three biological replicates (*n* = 3), except for XC (panel b) and EV (panel c-f) for which *n* = 2.

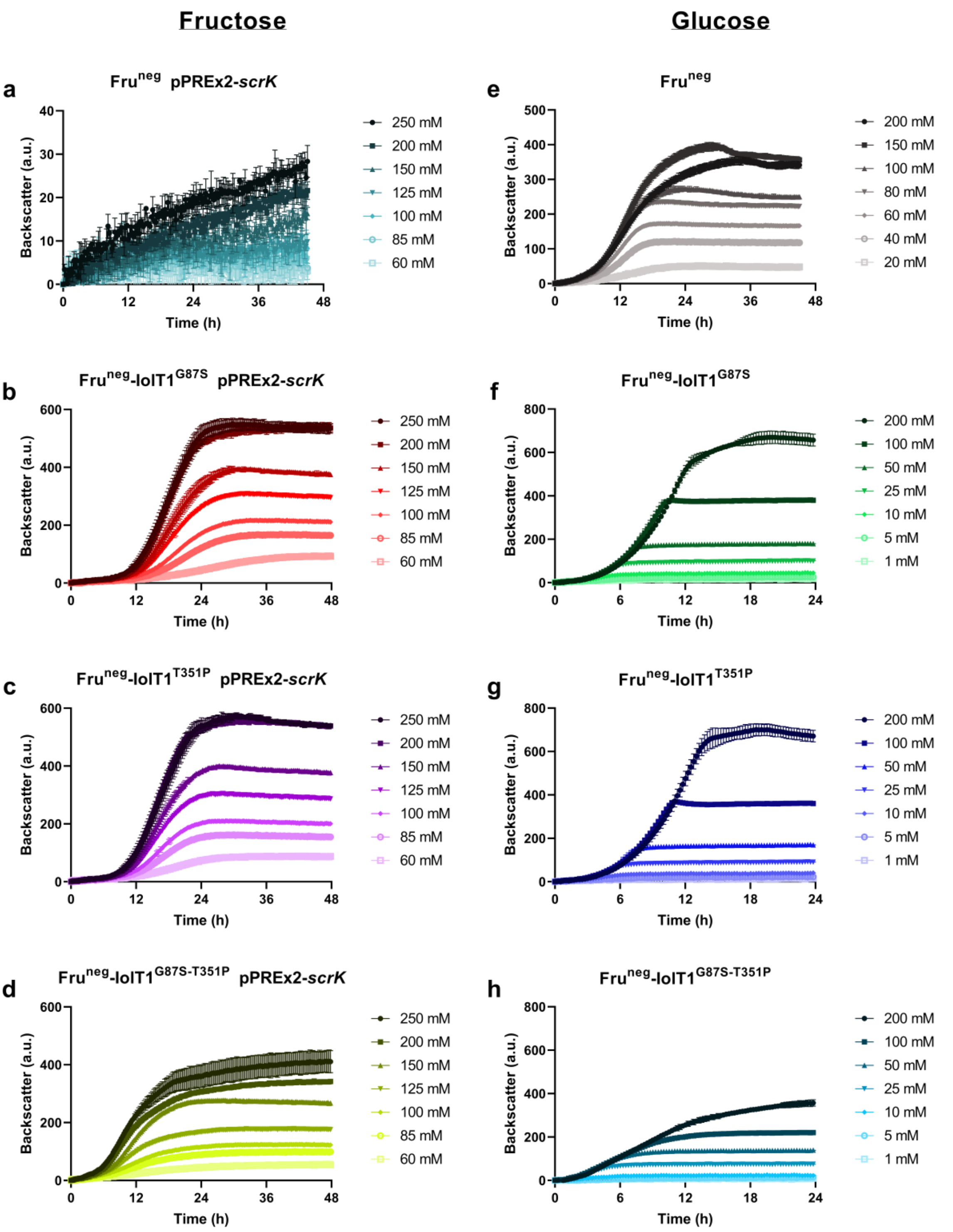

**Figure S3. Growth characteristics of *C. glutamicum* strains with wild-type IolT1, IolT1-G87S, IolT1-T351P and IolT1-G87S-T351P in minimal medium with varying concentrations of D-fructose and D-glucose. a-d** Growth experiments in CGXII medium with different D-fructose concentrations of the indicated strains. **e-h** Growth experiments in CGXII medium with different D-glucose concentrations of the indicated strains. All experiments were performed in a BioLector XT at 30°C and 1200 rpm. For each experiment, inoculation was performed to an $OD_{600}$ of 5. Strains with plasmid pPREx2-*scrK* were cultivated in CGXII medium supplemented with 1 mM IPTG and 25 µg/mL kanamycin. All data points given represent the mean ± S.D. of three biological replicates (*n* = 3).

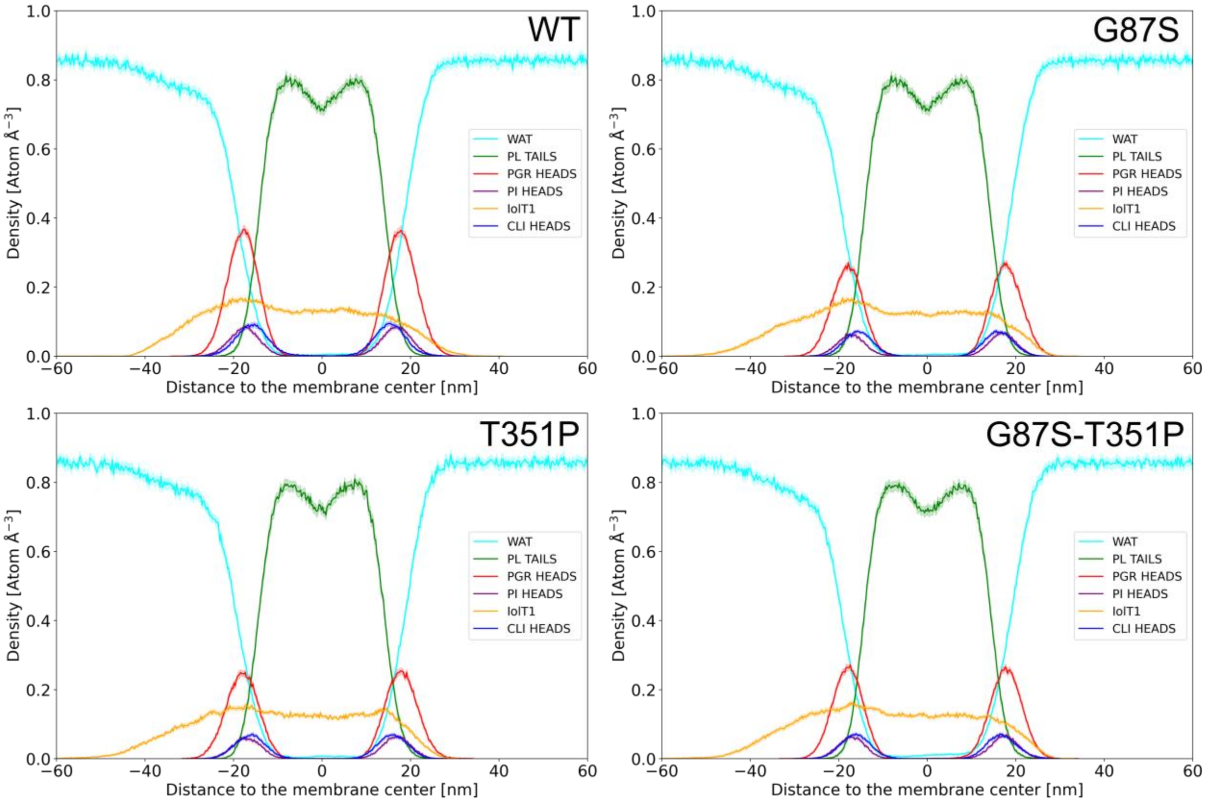

**Figure S4. Atom density profiles of membrane components averaged over five independent, unbiased MD simulations of outward-open configurations of wild-type IolT1 (WT), IolT1-G87S, IolT1-T351P, and IolT1-G87S-T351P for the phospholipid (PL) tails and head groups (PGR, phosphatidylglycerol; CLI, cardiolipin and PI, phosphatidylinositol).** The atom density profiles of water (WAT) are also depicted. The shaded area indicates the SEM over five independent replicas. The profiles correspond with those generally found by experiments and MD simulations for biological membranes [1-3]. Negative distances reflect the membrane leaflet oriented to the cytoplasm.

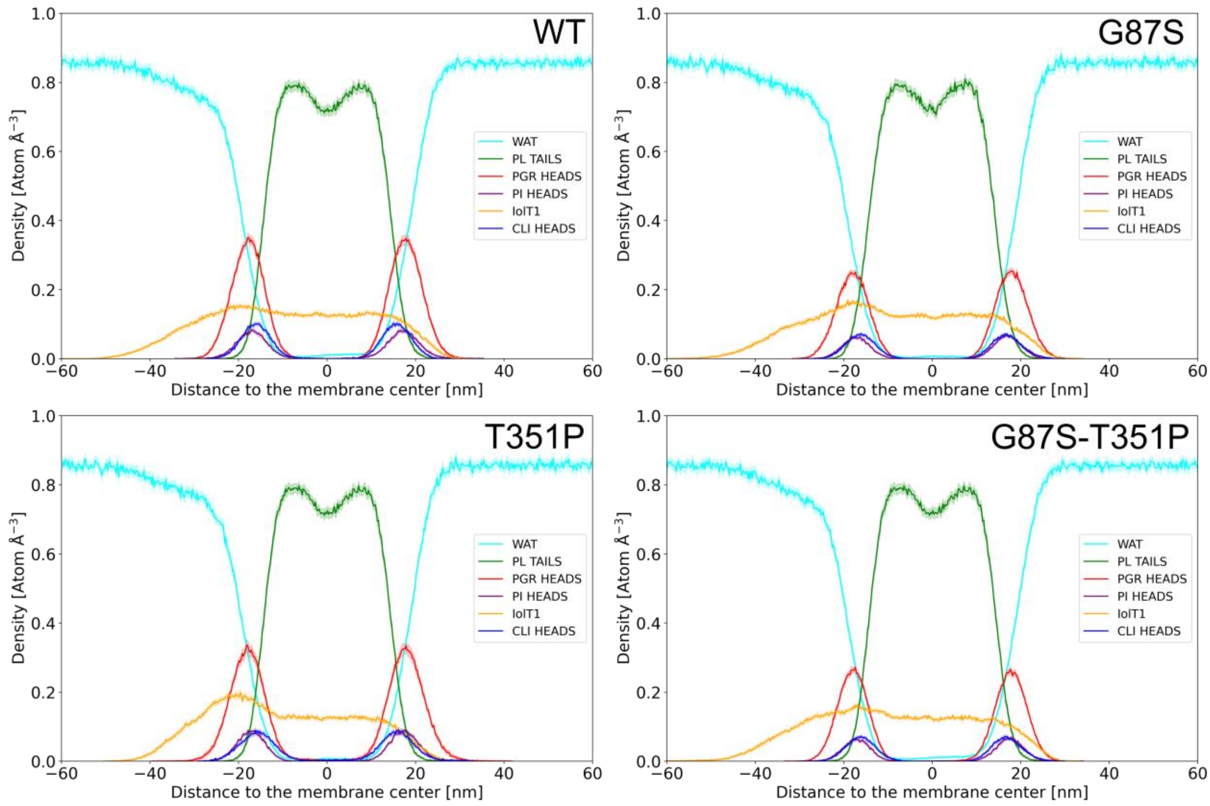

**Figure S5. Atom density profiles of membrane components averaged over five independent, unbiased MD simulations of outward-occluded β-D-glucopyranose-bound configurations of wild-type IolT1 (WT), IolT1-G87S, IolT1-T351P, and IolT1-G87S-T351P for the phospholipid (PL) tails and head groups (PGR, phosphatidylglycerol; CLI, cardiolipin and PI, phosphatidylinositol).** The atom density profiles of water (WAT) are also depicted. The shaded area indicates the SEM over five independent replicas. The profiles correspond with those generally found by experiments and MD simulations for biological membranes [1-3]. Negative distances reflect the membrane leaflet oriented to the cytoplasm.

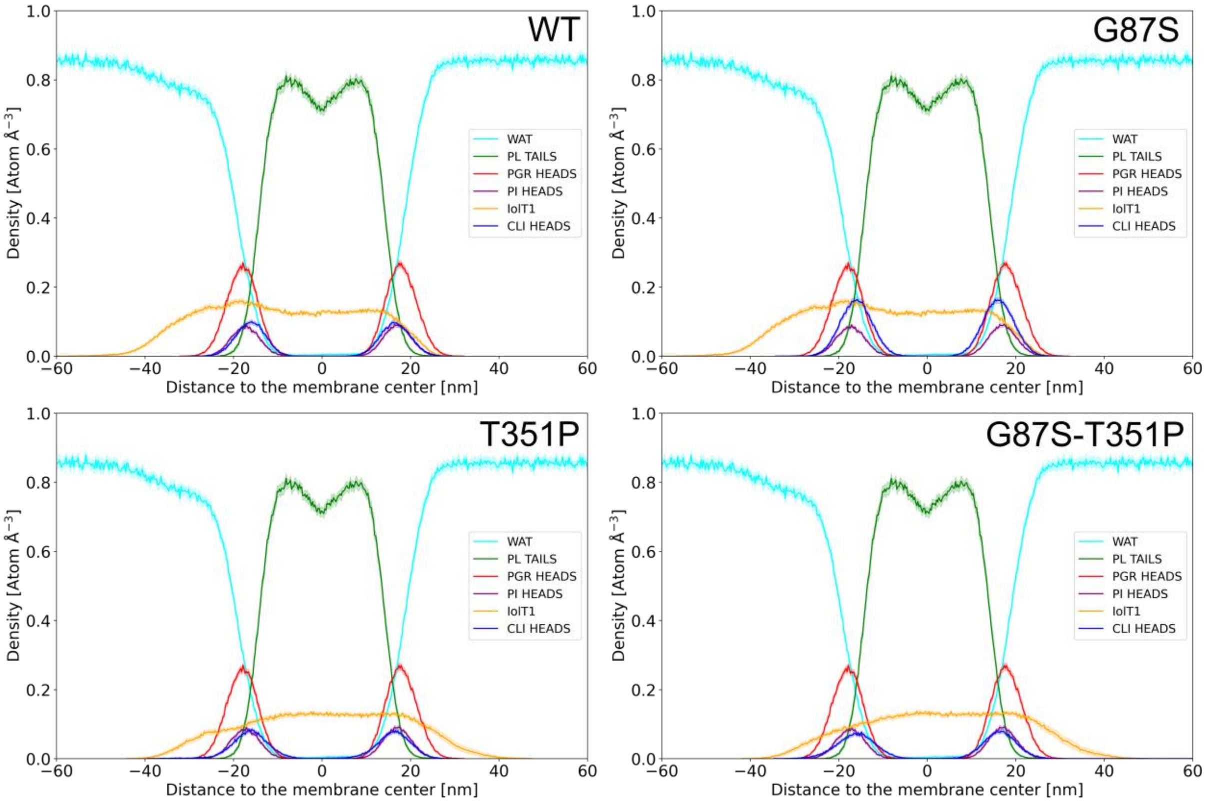

**Figure S6. Atom density profiles of membrane components averaged over five independent, unbiased MD simulations of outward-occluded β-D-fructofuranose-bound configurations of wild-type IolT1 (WT), IolT1-G87S, IolT1-T351P, and IolT1-G87S-T351P for the phospholipid (PL) tails and head groups (PGR, phosphatidylglycerol; CLI, cardiolipin and PI, phosphatidylinositol).** The atom density profiles of water (WAT) are also depicted. The shaded area indicates the SEM over five independent replicas. The profiles correspond with those generally found by experiments and MD simulations for biological membranes [1-3]. Negative distances reflect the membrane leaflet oriented to the cytoplasm.

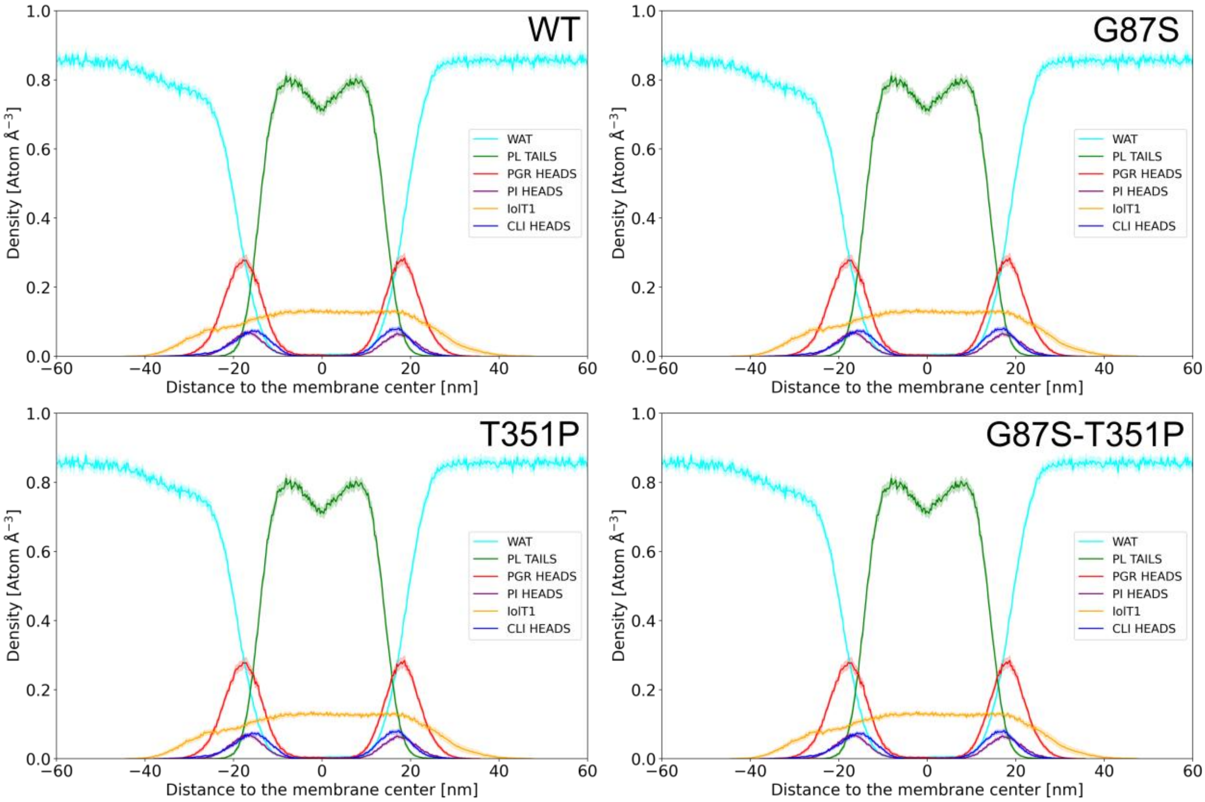

**Figure S7. Atom density profiles of membrane components averaged over five independent, unbiased MD simulations of inward-occluded β-D-glucopyranose-bound configurations of wild-type IolT1 (WT), IolT1-G87S, IolT1-T351P, and IolT1-G87S-T351P for the phospholipid (PL) tails and head groups (PGR, phosphatidylglycerol; CLI, cardiolipin and PI, phosphatidylinositol).** The atom density profiles of water (WAT) are also depicted. The shaded area indicates the SEM over five independent replicas. The profiles correspond with those generally found by experiments and MD simulations for biological membranes [1-3]. Negative distances reflect the membrane leaflet oriented to the cytoplasm.

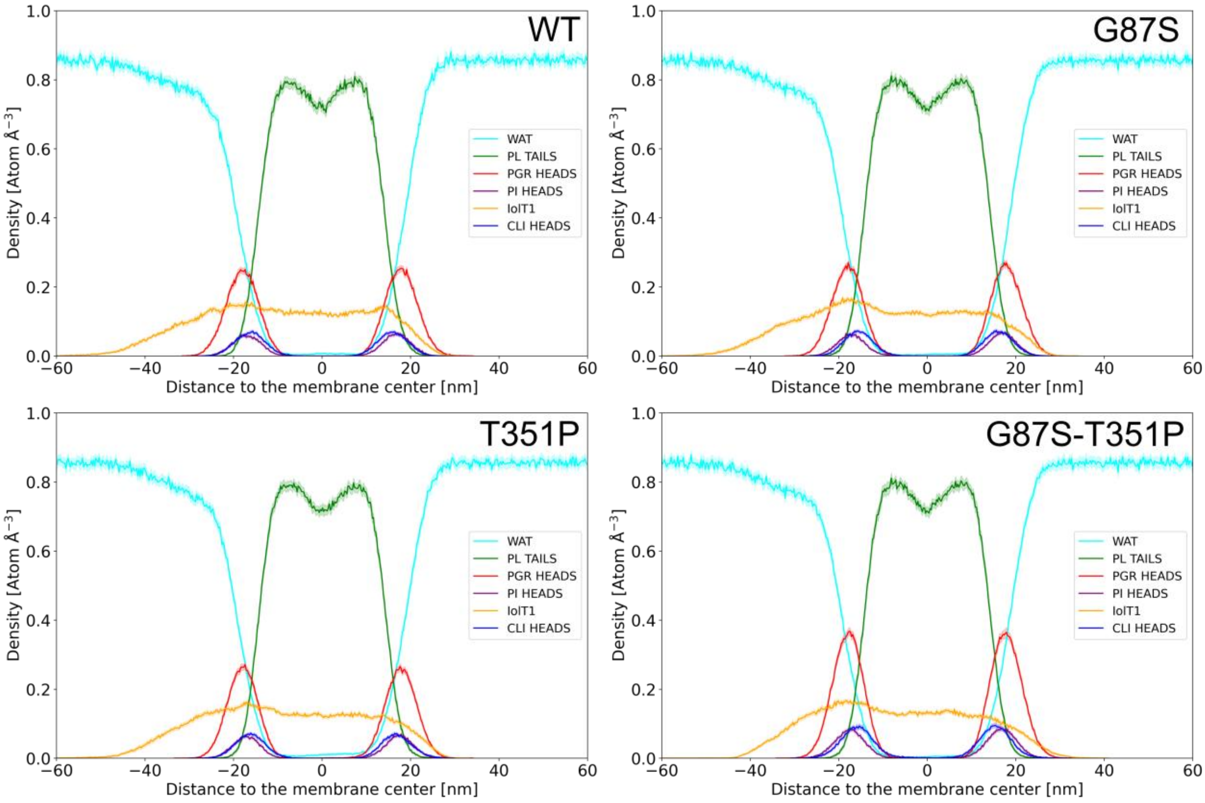

**Figure S8. Atom density profiles of membrane components averaged over five independent, unbiased MD simulations of inward-occluded β-D-fructofuranose-bound configurations of wild-type IolT1 (WT), IolT1-G87S, IolT1-T351P, and IolT1-G87S-T351P for the phospholipid (PL) tails and head groups (PGR, phosphatidylglycerol; CLI, cardiolipin and PI, phosphatidylinositol).** The shaded area indicates the SEM over five independent replicas. The profiles correspond with those generally found by experiments and MD simulations for biological membranes [1-3]. Negative distances reflect the membrane leaflet oriented to the cytoplasm.

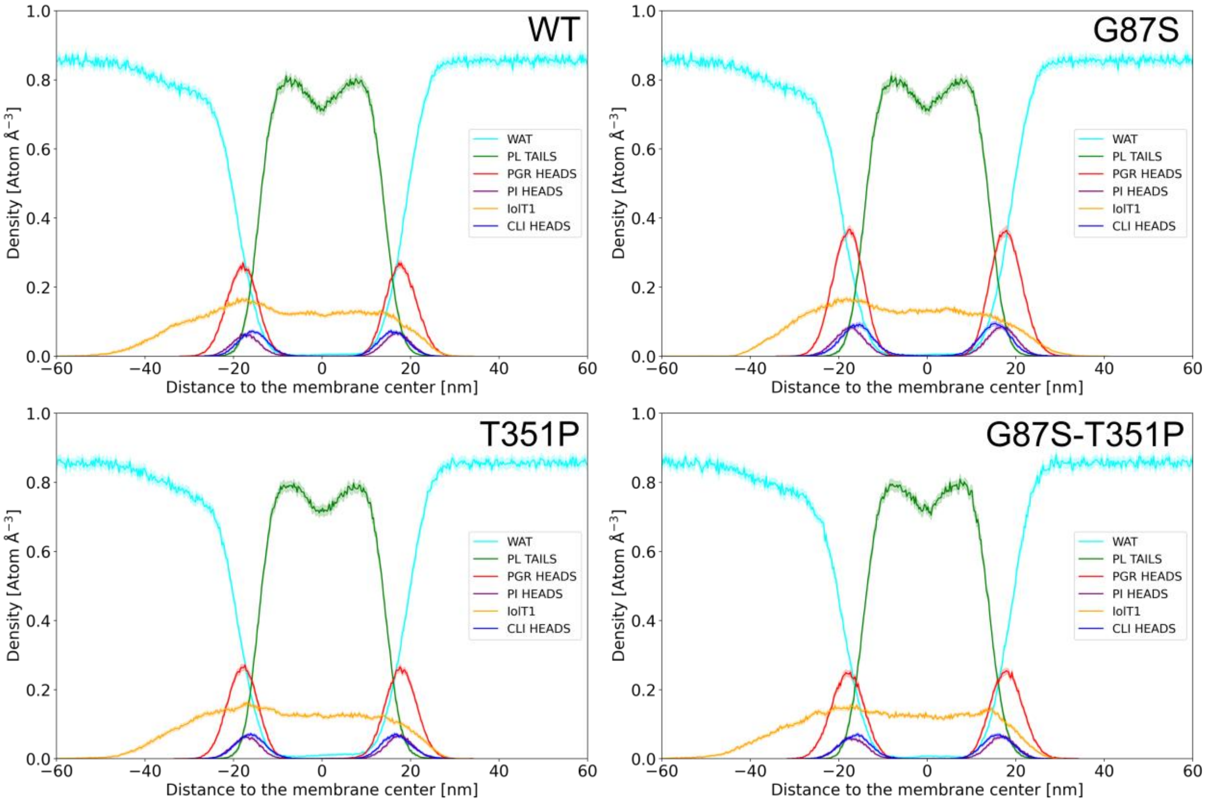

**Figure S9. Atom density profiles of membrane components averaged over five independent, unbiased MD simulations of inward-open configurations of wild-type IolT1 (WT), IolT1-G87S, IolT1-T351P, and IolT1-G87S-T351P for the phospholipid (PL) tails and head groups (PGR, phosphatidylglycerol; CLI, cardiolipin and PI, phosphatidylinositol).** The atom density profiles of water (WAT) are also depicted. The shaded area indicates the SEM over five independent replicas. The profiles correspond with those generally found by experiments and MD simulations for biological membranes [1-3]. Negative distances reflect the membrane leaflet oriented to the cytoplasm.

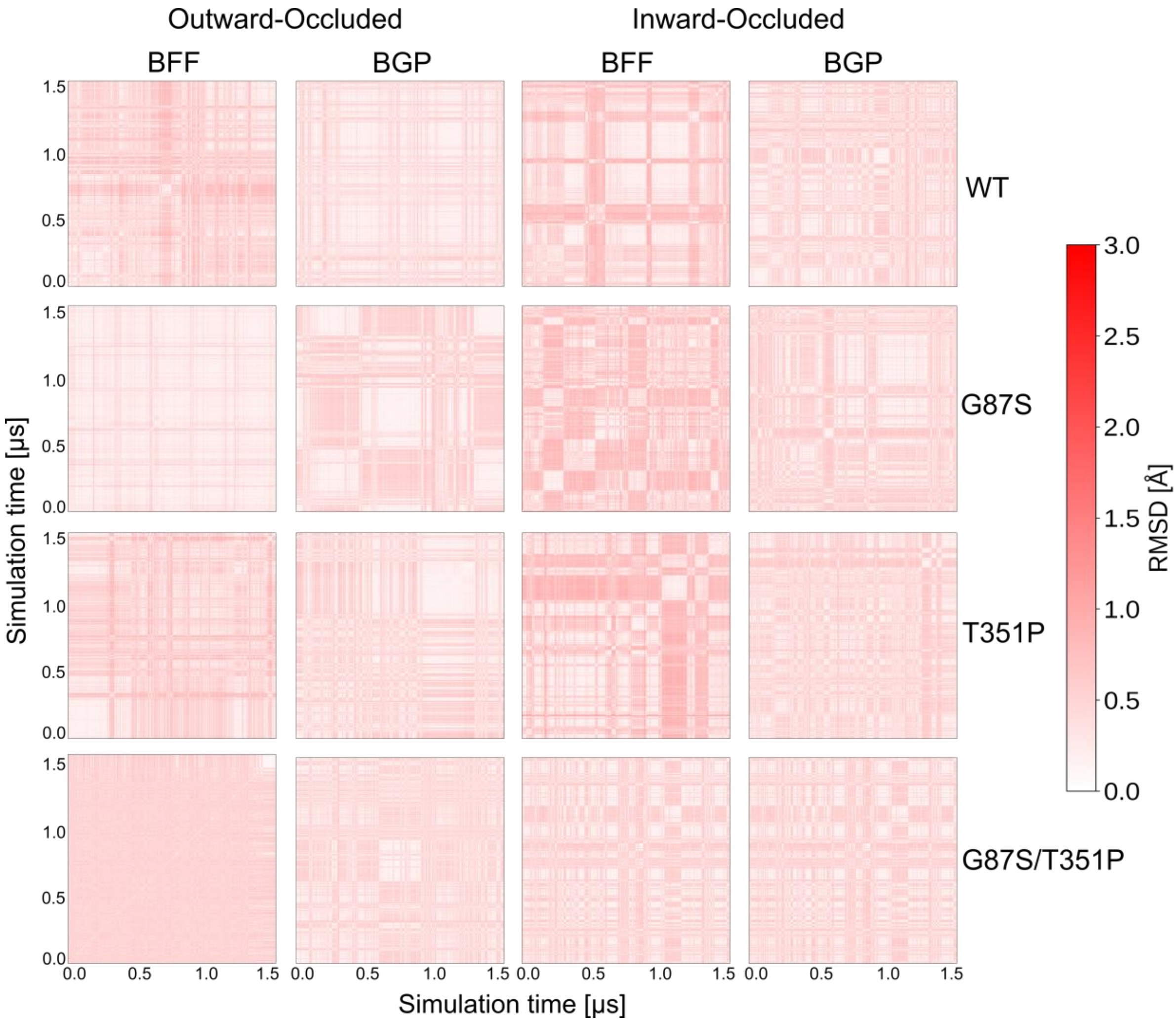

**Figure S10. Two-dimensional root-mean-square deviation (2D-RMSD) matrix comparing molecular dynamics simulation poses of β-D-glucopyranose (BGP) or β-D-fructofuranose (BFF) molecules in MD simulations of outward-occluded and inward-occluded IolT1 and the variants IolT1-G87S, IolT1-T351P, and IolT1-G87S-T351P.** In all frames, the BGP and BFF molecules preserve their position during 1.5 µs of simulation time with respect to the first frame (RMSD < 2 Å) across five different replicas. The 2D-RMSD of all no hydrogen ligand atoms was computed after superpositioning the Cα atoms of IolT1, with warmer colors denoting increased mobility.

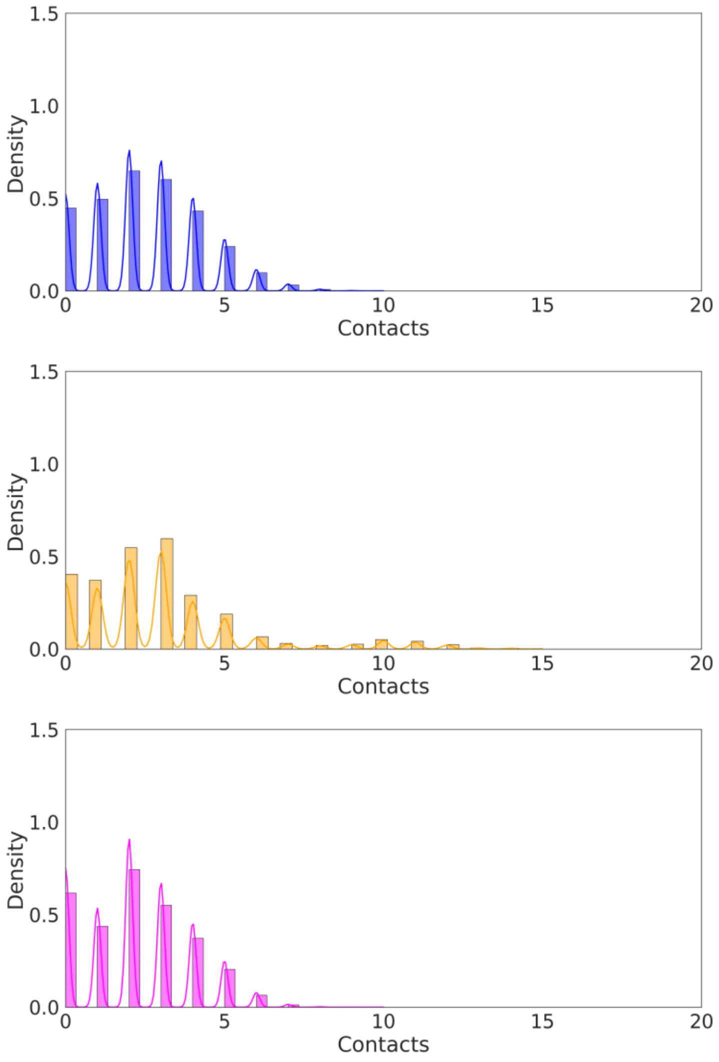

**Figure S11. Formation of the D46-R126 salt bridge in the inward-open conformation of wild-type IolT1 (blue), the G87S variant (orange), and the T351P variant (magenta).** Histogram showing the probability density of the number of contacts. A contact was counted when each of the two side-chain oxygens of D46 are within a distance < 4.0 Å from any side-chain nitrogen of R126. In all replicas, the salt bridge is overall maintained over 1.5 µs of production simulation time. The histogram is normalized to unit area, and the curve line represents the corresponding kernel density estimate (KDE).

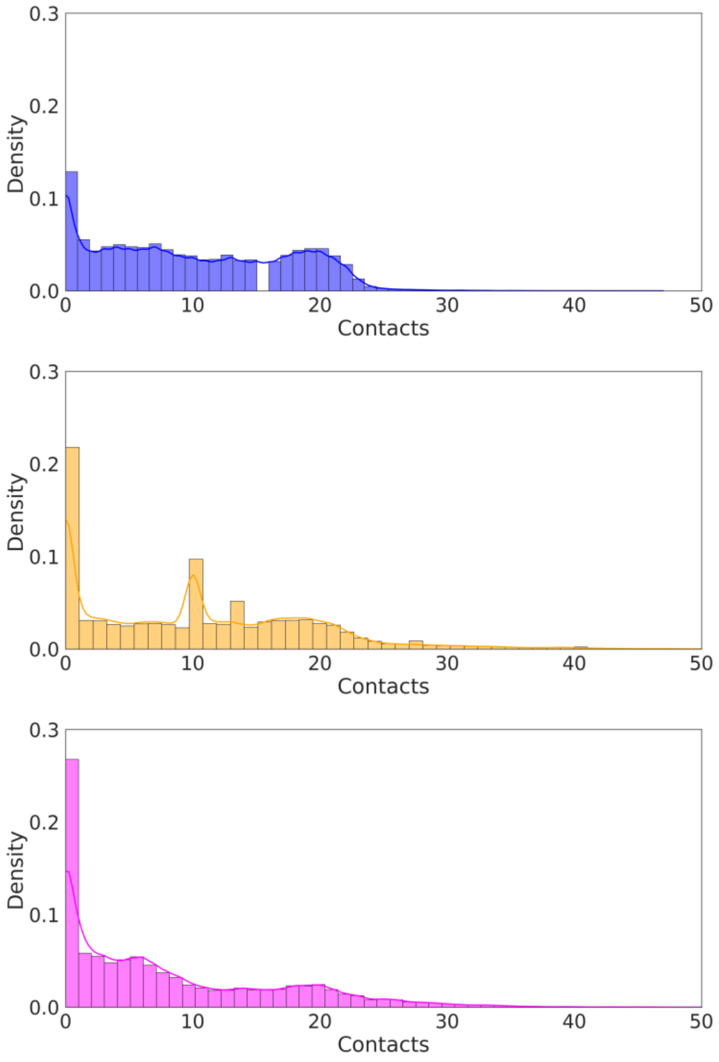

**Figure S12. Formation of the E400-R339 salt bridge in the inward-open conformation of wild-type IolT1 (blue), the G87S variant (orange), and the T351P variant (magenta).** Histogram showing the probability density of the number of contacts. A contact was counted when each of the two side-chain oxygens of E400 is within a distance < 4. 0 Å from any side-chain nitrogen of R339. In all replicas, the salt bridge is overall maintained over 1.5 μs of production simulation time. The histogram is normalized to unit area, and the curve line represents the corresponding kernel density estimate (KDE).

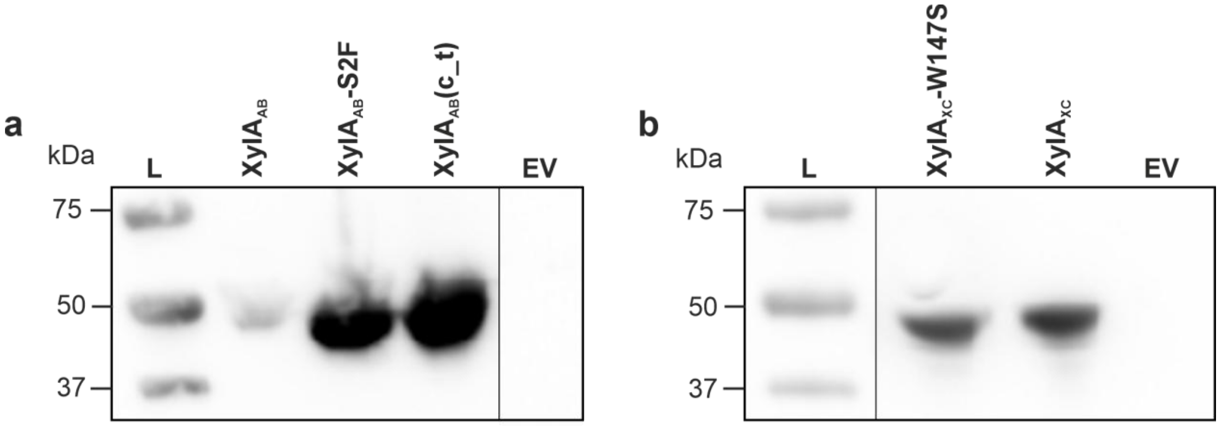

**Figure S13. Influence of the indicated mutations identified by ALE on XylA$_{AB}$ (a) and XylA$_{XC}$ (b) protein levels in *C. glutamicum* Fru$^{neg}$.** The strains were cultured in 10 mL of CGXII medium with 10 g/L of D-glucose, 25 µg/mL kanamycin and 1 mM IPTG for 24 h and then used for preparation of cell-free extracts. 50 µg protein of each strain was separated by SDS-PAGE and blotted onto a nitrocellulose membrane (GE Healthcare, Chicago, USA). The Strep-tagged XylA variants were visualized with Strep-Tactin-HRP conjugate. L: Precision Plus Protein$^{TM}$ Dual Xtra Prestain Protein Standard with molecular masses given in kDa, EV: Soluble protein fraction of the Fru$^{neg}$ strain with empty vector (pPREx2) control.

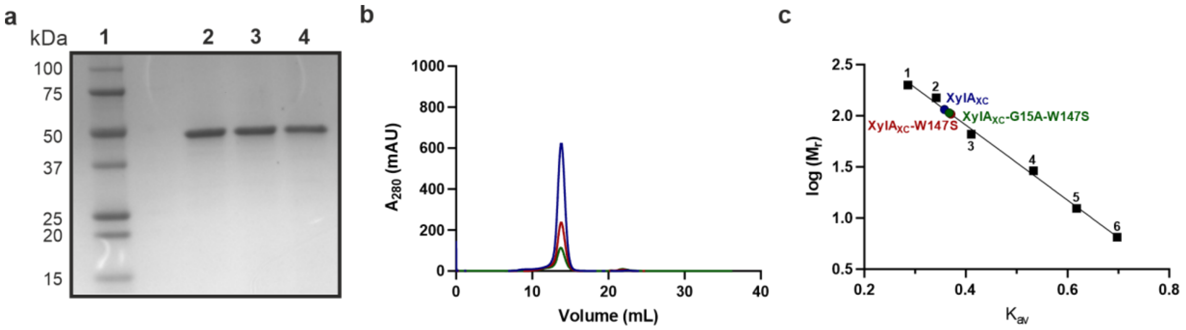

**Figure S14. Purification of XylA$_{XC}$ variants. a** SDS-PAGE analysis of XylA$_{XC}$ (lane 2), XylA$_{XC}$-W147S (lane 3) and XylA$_{XC}$-G15A-W147S (lane 4) after Ni-NTA affinity and size exclusion chromatography. The calculated molecular mass of the XylA$_{XC}$ variants is 48.8 kDa. Precision Plus Protein$^{TM}$ Dual Xtra Prestained Protein Standard was used as marker (lane 1). **b** Size exclusion chromatography profiles of XylA$_{XC}$ (blue), XylA$_{XC}$-W147S (red) and XylA$_{XC}$-G15A-W147S (green) via a Superdex 200 10-300GL column. Protein was detected via absorbance at 280 nm. **c** Molecular mass determination of XylA$_{XC}$ variants via a Superdex 200 10-300GL calibration curve: 1, amylase (200 kDa); 2, alcohol dehydrogenase (150 kDa); 3, bovine serum albumin (66 kDa); 4, carbonic anhydrase (29 kDa); 5, cytochrome *c* (12.4 kDa); 6, aprotinin (6.5 kDa). $K_{av}$ values of XylA$_{XC}$ variants were indicated as red, blue, green, purple and yellow dots on the calibration curve. The native molecular mass of the XylA$_{XC}$ variants was calculated to be 102-120 kDa, corresponding to a dimeric structure.

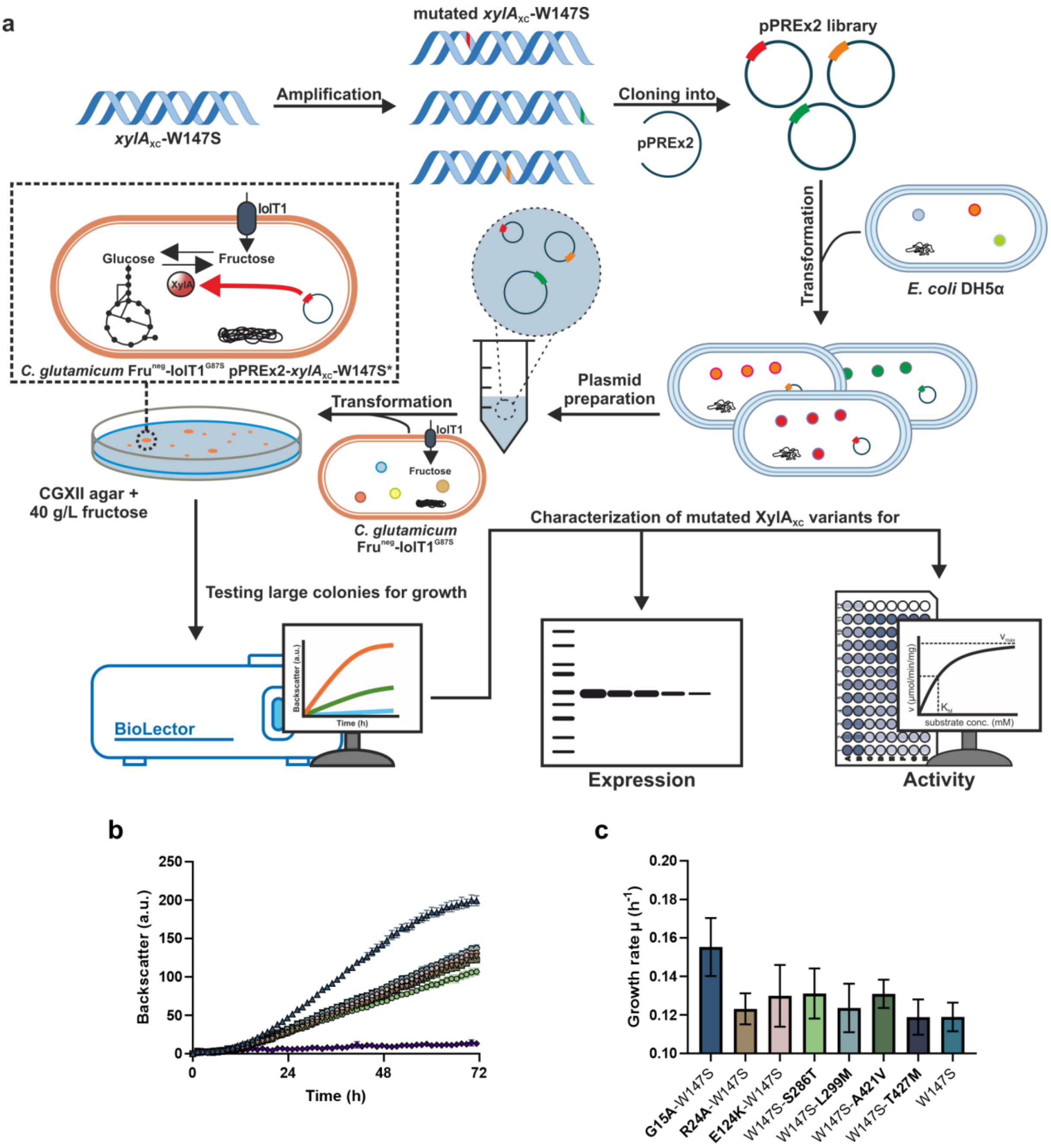

**Figure S15. Growth-based directed evolution of XylA$_{XC}$-W147S. a** Experimental workflow for random mutagenesis of *xylA*$_{XC}$-W147S. Mutations were introduced via the GeneMorph II random mutagenesis kit (Agilent Technologies, Santa Clara, USA). Mutated *xylA*$_{XC}$ variants were cloned into pPREx2 and propagated in *E. coli* DH5α. Isolation of assembled plasmids and transformation of *C. glutamicum* Fru$^{neg}$-IolT1$^{G87S}$ revealed faster growing clones on CGXII D-fructose agar plates, which were then evaluated for growth in liquid D-fructose minimal medium in the BioLector I system. Obtained mutated XylA$_{XC}$-W147S variants resulting in better growth were subsequently selected for further characterization via Western blot analysis for expression and/or D-glucose isomerase assays for protein activity determination. **b** Growth-based evaluation of novel *xylA*$_{XC}$ mutations obtained after random mutagenesis of *xylA*$_{XC}$-W147S. For this, novel *xylA*$_{XC}$ mutations were introduced separately into the pPREx2-*xylA*$_{XC}$-W147S plasmid. The newly constructed pPREx2 plasmids were then used for the transformation of *C. glutamicum* Fru$^{neg}$-IolT1$^{G87S}$. The growth experiment was performed in the BioLector XT at 30 °C and 1200 rpm in CGXII with 40 g/L of (w/v) D-fructose, 25 µg/mL kanamycin and 25 µM IPTG for 72 h. Inoculation was set to an $OD_{600}$ of 1. **c** Growth rates of the strains shown in panel (b). All data points represent mean ± S.D. from three biological replicates (*n* = 3).

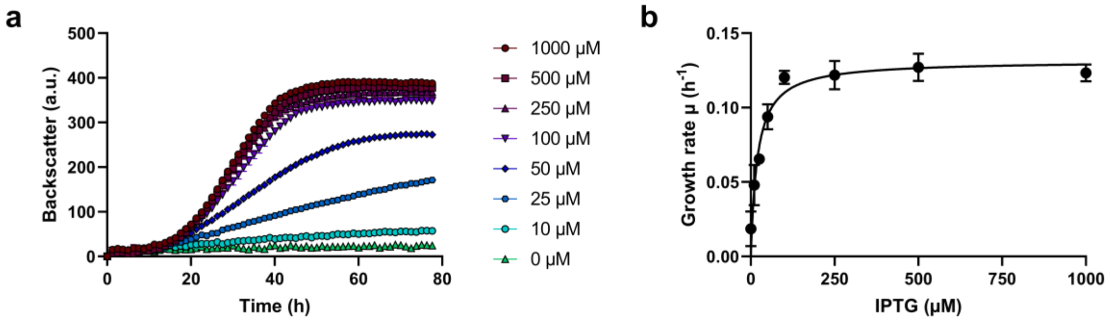

**Figure S16. Correlation between IPTG concentration and growth rate for *C. glutamicum* Fru$^{neg}$-IolT1$^{G87S}$ pPREx2-*xylA*$_{XC}$-W147S in D-fructose minimal medium. a** The growth experiment was performed in CGXII medium with 40 g/L D-fructose, 25 µg/mL kanamycin and varying IPTG concentrations (0 – 1000 µM) at 30°C and 1200 rpm for 78 h in a BioLector XT. Growth rates were determined in the period from 18-22 h for all IPTG concentrations. **b** Michaelis-Menten non-linear regression fit of growth rates plotted against IPTG concentration was performed to obtain growth parameters ($\mu_{max}$, $K_S$). All data points given represent average values with standard deviations of three biological replicates.

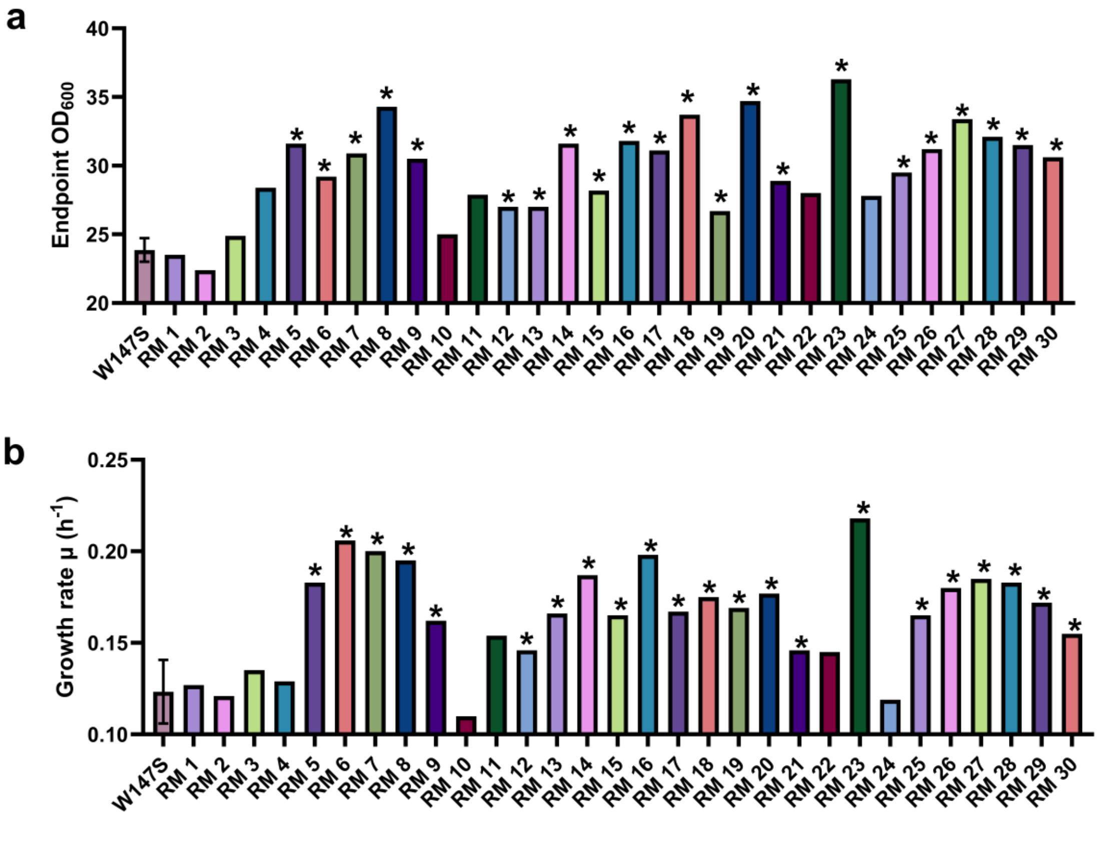

**Figure S17. Growth analysis of visually fast-growing Fru$^{neg}$-IolT1$^{G87S}$ pPREx2-*xylA*$_{XC}$-W147S clones obtained after two rounds of random mutagenesis. a** Endpoint $OD_{600}$ of clones obtained after random mutagenesis (RM1-30) and of a Fru$^{neg}$-IolT1$^{G87S}$ pPREx2-*xylA*$_{XC}$-W147S control strain (W147S). **b** Growth rate of tested clones and the W147S control. Plasmid sequencing was performed for the clones marked with an asterisk (*).

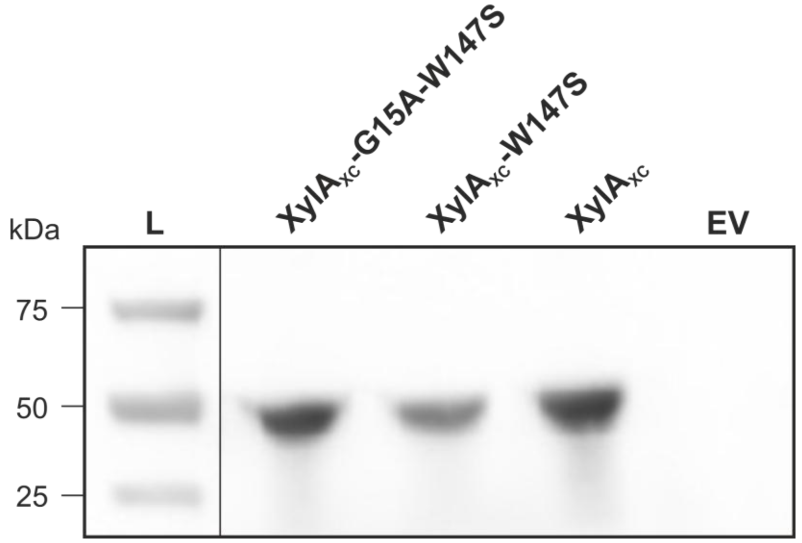

**Figure S18. Influence of the indicated XylA$_{XC}$ mutations on the corresponding protein levels in *C. glutamicum* Fru$^{neg}$.** The strains were cultured in CGXII medium with 10 g/L of D-glucose, 25 µg/mL kanamycin and 1 mM IPTG for 24 h and then used for preparation of cell-free extracts. 50 µg protein of each strain was separated by SDS-PAGE and blotted onto a nitrocellulose membrane (GE Healthcare, Chicago, USA). The Strep-tagged XylA$_{XC}$ variants were visualized with Strep-Tactin-HRP conjugate. L: Precision Plus Protein™ Dual Xtra Prestain Protein Standard with molecular masses given in kDa, EV: soluble protein fraction of the Fru$^{neg}$ strain with the empty vector (pPREx2) control.

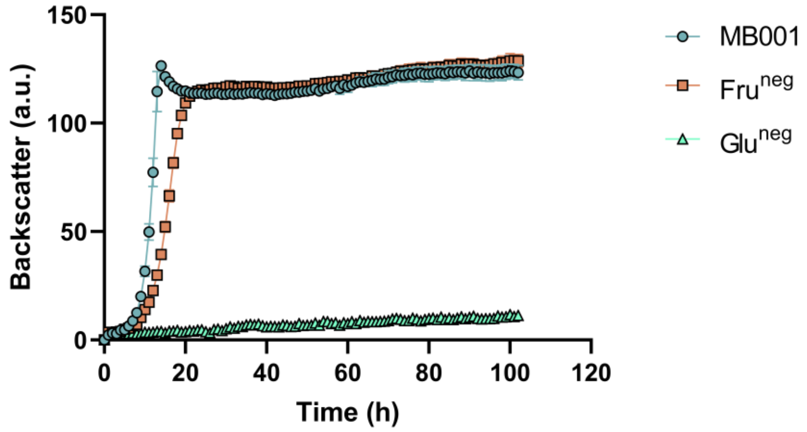

**Figure S19. Growth experiment of the *C. glutamicum* Glu$^{neg}$ strain in CGXII medium with 40 g/L D-glucose.** The strains MB001 and Fru$^{neg}$ were used as positive controls. The experiment was conducted in a BioLector II at 30°C and 1200 rpm for 102 h. All data points given represent mean ± S.D. from three biological replicates (*n* = 3).

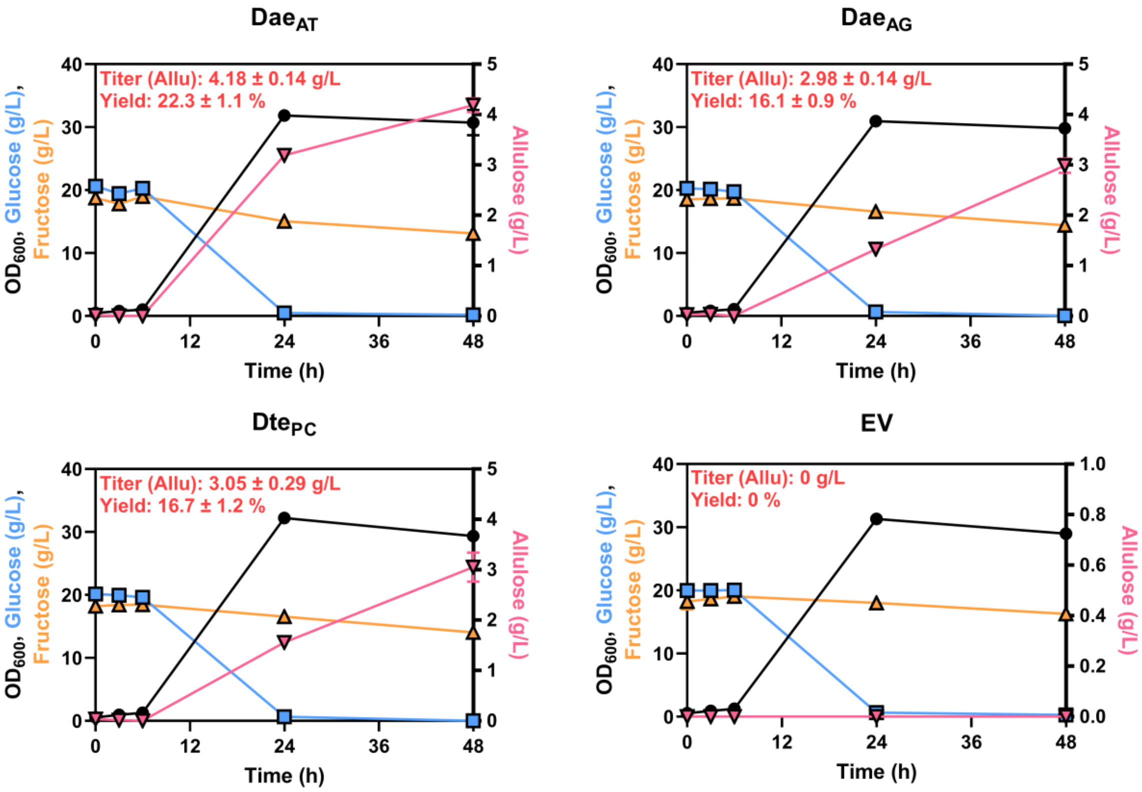

**Figure S20. Analysis of D-allulose formation from D-fructose by different D-allulose 3- and D-tagatose 3-epimerases in *C. glutamicum* Fru$^{neg}$.** The Fru$^{neg}$ strains carrying either pPREx2-*dae*$_{AT}$ (Dae$_{AT}$), pPREx2-*dae*$_{AG}$ (Dae$_{AG}$), pPREx2-*dte*$_{PC}$ (Dte$_{PC}$), or pPREx2 (EV) were used for the cultivation. The genes encode the D-allulose 3-epimerases of *Agrobacterium tumefaciens* (AG) and *Arthrobacter globiformis* (AG), as well as the D-tagatose 3-epimerase of *Pseudomonas cichorii* (PC). The experiment was conducted in baffled shake flasks with CGXII medium containing 20 g/L D-glucose, 20 g/L D-fructose, 25 µg/mL kanamycin and 1 mM IPTG at 30°C and 130 rpm. Cultures were inoculated to an initial OD$_{600}$ of 0.5. Samples of the supernatant were used for OD$_{600}$ measurement and HPLC sample preparation and taken after 0, 3, 6, 24 and 48 h. All data points represent mean ± S.D. from three biological replicates (*n* = 3).

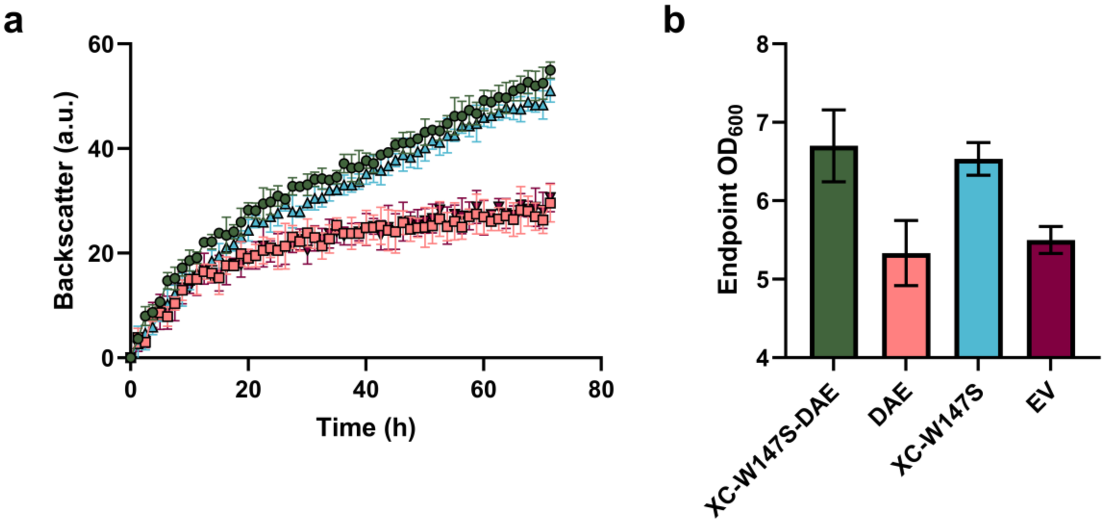

**Figure S21. Cultivation of *C. glutamicum* Glu$^{neg}$ carrying either pPREx2-*xylA*$_{XC}$-W147S-*dae*$_{AT}$ (XC-W147S-DAE), pPREx2-*dae*$_{AT}$ (DAE), pPREx2-*xylA*$_{XC}$W147S (XC-W147S), or pPREx2 (EV) plasmid in D-glucose minimal medium.** **a** Growth experiment performed in a BioLector XT at an initial $OD_{600}$ of 5 in CGXII medium with 40 g/L D-glucose, 25 µg/mL kanamycin and 1 mM IPTG. The cultivation was conducted at 30°C and 1200 rpm for 72 h. **b** Endpoint $OD_{600}$ values of cultures after the BioLector experiment was terminated. All data points represent mean ± S.D. from three biological replicates (*n* = 3).

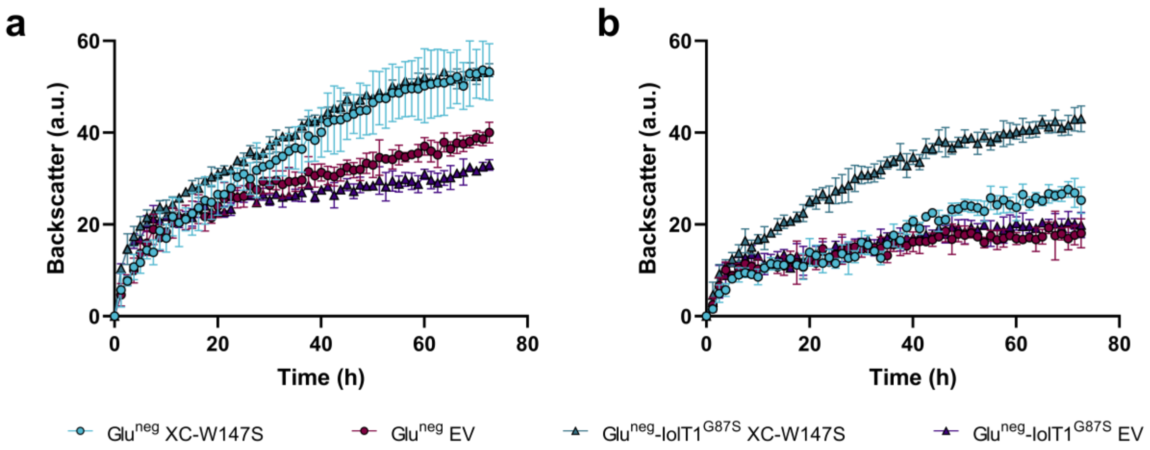

**Figure S22. Growth of *C. glutamicum* Glu$^{neg}$ and *C. glutamicum* Glu$^{neg}$-IolT1$^{G87S}$ with pPREx2-*xylA*$_{XC}$-W147S (XC-W147S) or pPREx2 (EV) in D-glucose minimal medium.** Cultivation experiment in CGXII medium with (**a**) 40 g/L D-glucose or (**b**) 40 g/L D-fructose, 25 µg/mL kanamycin and 1 mM IPTG. Inoculation was performed to an initial $OD_{600}$ of 5. The experiment was conducted at 30°C and 1200 rpm for 72 h in a BioLector XT. All data points represent mean ± S.D. from three biological replicates (*n* = 3).

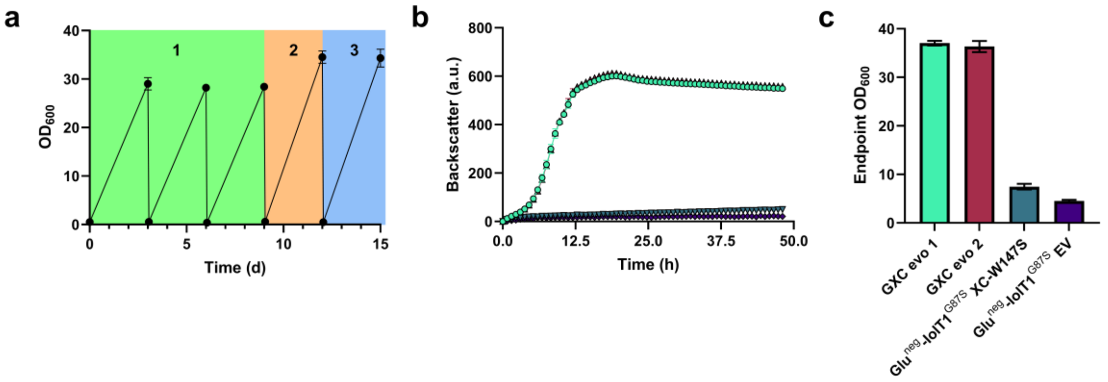

**Figure S23. Adaptive laboratory evolution of Glu$^{neg}$-IolT1$^{G87S}$ with pPREx2-*xylA*$_{XC}$-W147S in D-glucose minimal medium (a) and evaluation of obtained evolved strains (GXC evo) in a BioLector growth experiment (b, c). a** The ALE experiment was conducted in 100 mL baffled shake flasks with 10 mL CGXII medium containing 30 g/L of D-glucose, 25 µg/mL kanamycin, 1 mM IPTG and either 10 g/L (1, green), 2 g/L (2, orange), or 0 g/L (3, blue) of D-gluconate. Cultivation was performed at 30°C and 130 rpm for 3 days, after which a transfer to fresh medium was conducted. Evolved strains of Glu$^{neg}$-IolT1$^{G87S}$ with pPREx2-*xylA*$_{XC}$-W147S were obtained after a total cultivation time of 15 days, including four transfers. **b** Evolved and unevolved strains were evaluated in a BioLector growth experiment in CGXII medium with 40 g/L of D-glucose, 25 µg/mL kanamycin and 1 mM IPTG at 30°C and 1200 rpm for 48 h. **c** After termination of the cultivation, final $OD_{600}$ values of the respective cultures were determined. All data points represent mean ± S.D. from three biological replicates (*n* = 3), except for the ALE experiment (*n* = 2).

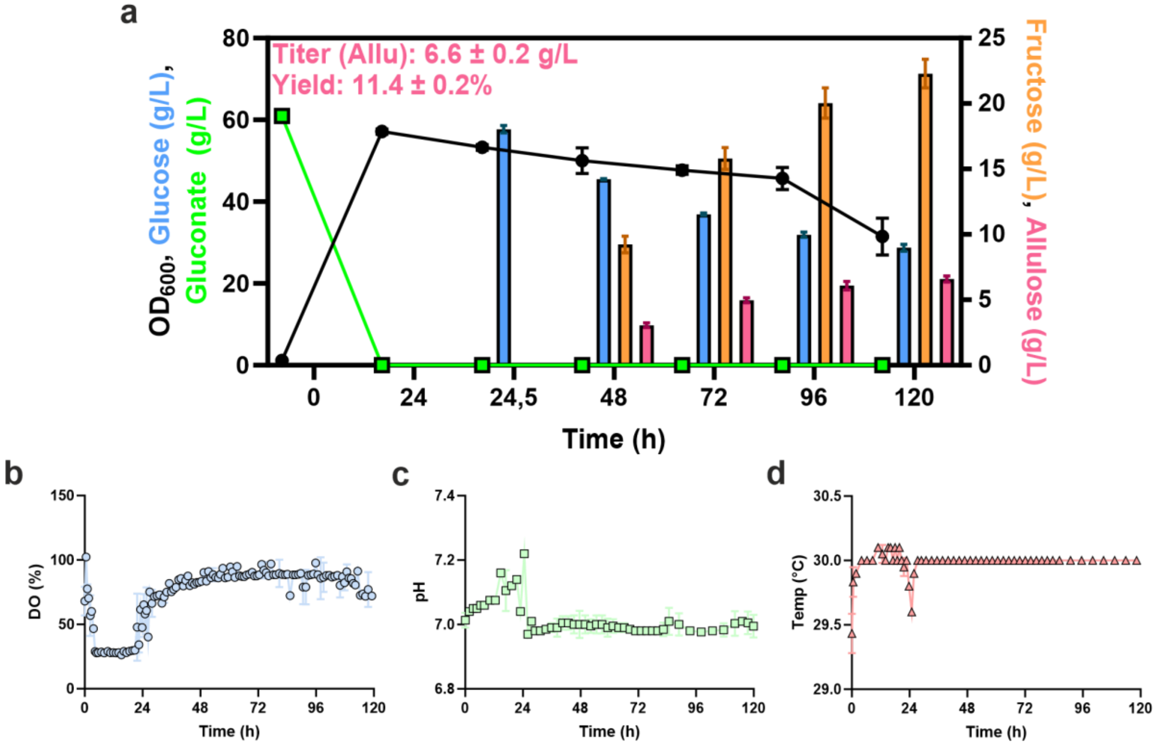

**Figure S24. High-cell-density biotransformation of *C. glutamicum* Suc$^{neg}$-IolT1$^{G87S}$ for the conversion of D-glucose to D-allulose. a** Cultivation of Suc$^{neg}$-IolT1$^{G87S}$ strain containing pPREx2-*xylA*$_{XC}$-G15A-W147S-*dae*$_{AT}$ in a DASGIP bioreactor with 1 L of CGXII medium supplemented with 60 g/L of D-gluconate, 25 µg/mL kanamycin and 1 mM IPTG. After D-gluconate was completely consumed, D-glucose was manually added to the medium up to a final concentration of 58 g/L. Samples of the supernatant were taken every 24 h for a total of 120 h. Growth was measured via $OD_{600}$ and sugar concentrations were determined via HPLC. Over the course of the fermentation, dissolved oxygen saturation (**b**), changes in pH values (**c**) and temperature changes (**d**) were measured. The data shown represents the mean ± S.D. from three biological replicates (*n* = 3).

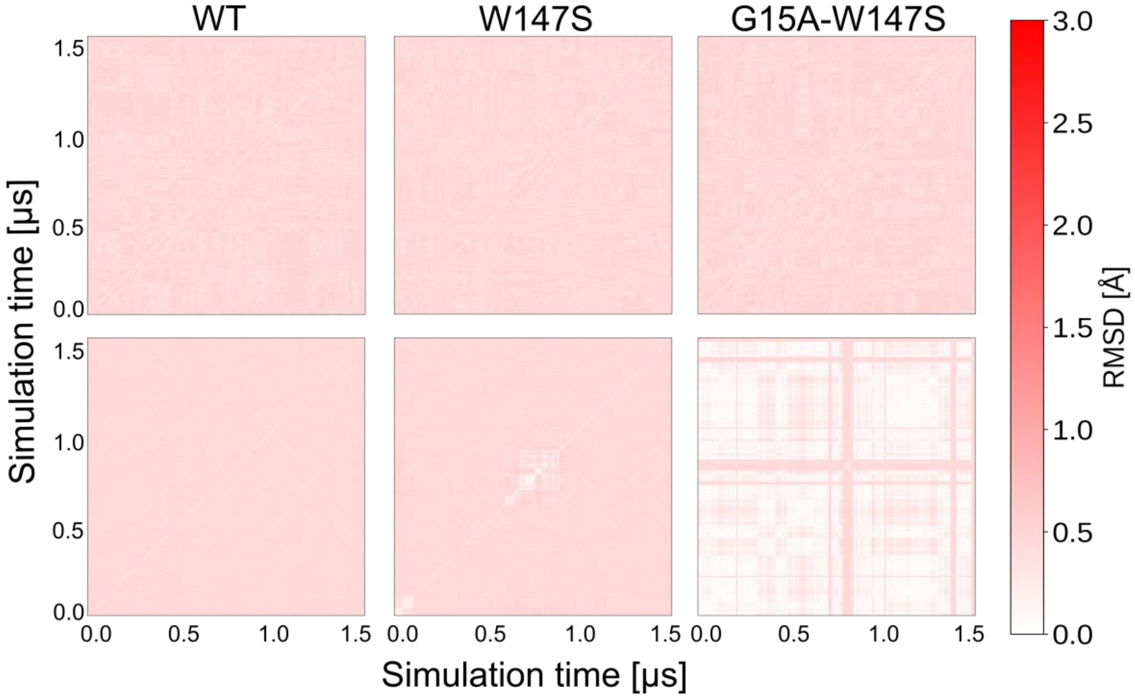

**Figure S25. Two-dimensional root-mean-square deviation (2D-RMSD) matrix comparing molecular dynamics simulation poses of (top) the XylA$_{XC}$ homodimer or (bottom) β-D-glucopyranose-$Mg^{2+}$ ligands in MD simulations of XylA$_{XC}$ WT, W147S, and G15A-W147S.** In all frames, the XylA$_{XC}$ homodimer and β-D-glucopyranose-$Mg^{2+}$ ligands retain their positions over 1.5 μs of production run with respect to the first frame (RMSD < 2 Å) across five independent replicas. The 2D-RMSD of the XylA$_{XC}$ protein backbone and the 2D-RMSD of all ligand non-hydrogen atoms (β-D-glucopyranose and $Mg^{2+}$ were considered in the ligand mask) were computed after superpositioning the Cα atoms of XylA$_{XC}$, with warmer colors denoting increased mobility.

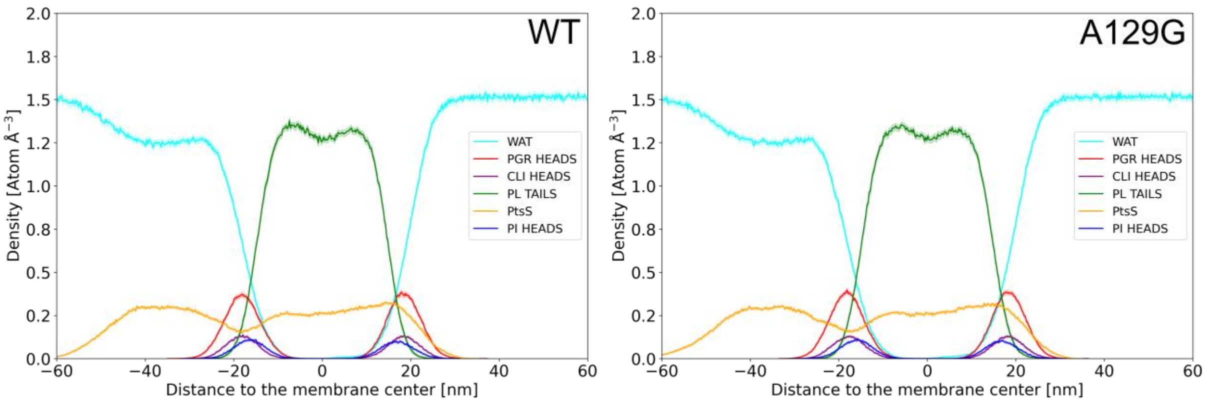

**Figure S26. Atom density profiles of membrane components averaged over five independent, unbiased MD simulations of PtsS configurations for the phospholipid (PL) tails and head groups (PGR, phosphatidylglycerol; CLI, cardiolipin and PI, phosphatidylinositol). The atom density profiles of water (WAT) are also depicted.** The shaded area indicates the SEM over five independent replicas. The profiles are concordant with those generally found by experiments and MD simulations for biological membranes [1-3]. Negative distances reflect the membrane leaflet oriented to the cytoplasm.

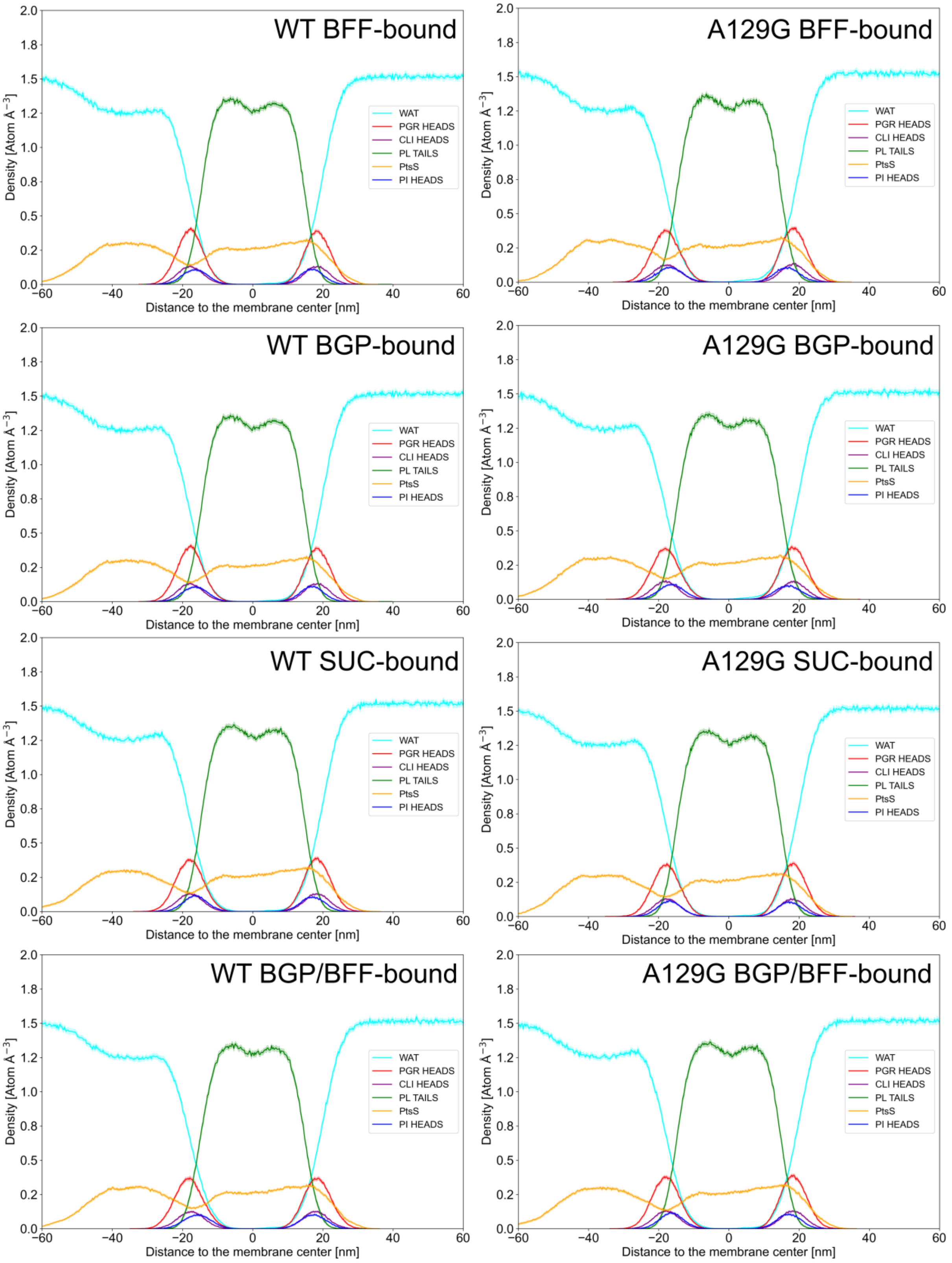

**Figure S27. Atom density profiles of membrane components averaged over five independent, unbiased MD simulations of PtsS configurations for the phospholipid (PL) tails and head groups (PGR, phosphatidylglycerol; CLI, cardiolipin and PI, phosphatidylinositol). The atom density profiles of water (WAT) are also depicted.** The shaded area indicates the SEM over five independent replicas. The profiles are concordant with those generally found by experiments and MD simulations for biological membranes [1-3]. BFF: β-D-fructofuranose, BGP: β-D-glucopyranose, SUC: D-sucrose. Negative distances reflect the membrane leaflet oriented to the cytoplasm.

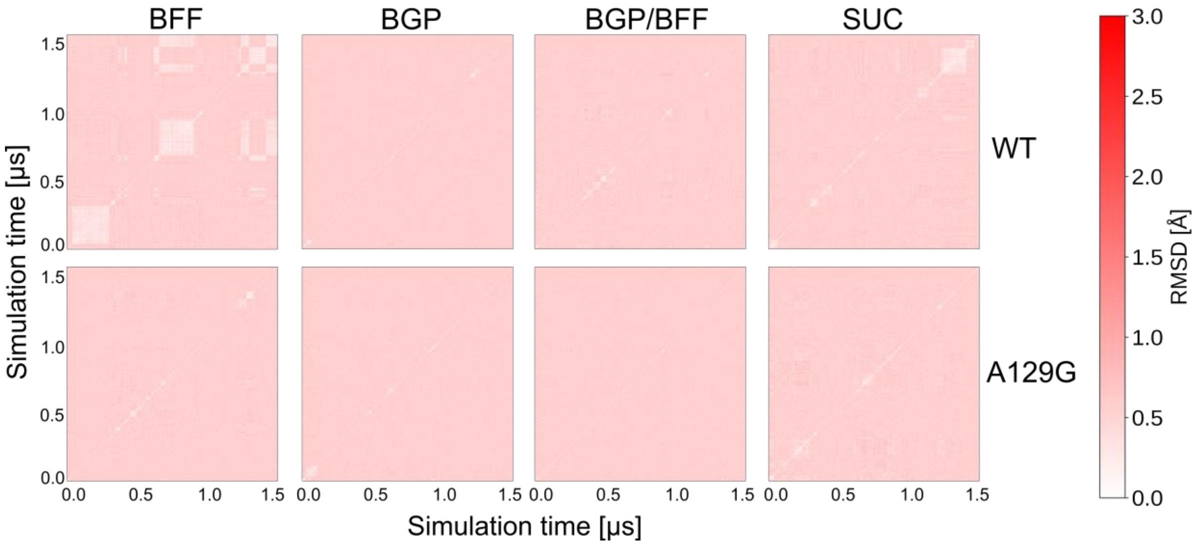

**Figure S28. Two-dimensional root-mean-square deviation (2D-RMSD) matrix comparing molecular dynamics simulation poses of β-D-fructofuranose (BFF), β-D-glucopyranose (BGP), D-sucrose (SUC), or BGP/BFF bound to the PtsS homodimer in MD simulations of PtsS WT (top) and PtsS-A129G (bottom).** In all frames, the ligands retain their position during 1.5 µs of simulation time with respect to the first frame (RMSD < 2 Å) across five different replicas. The 2D-RMSD of all ligand non-hydrogen atoms was computed after superpositioning the Cα atoms of PtsS, with warmer colors denoting increased mobility.

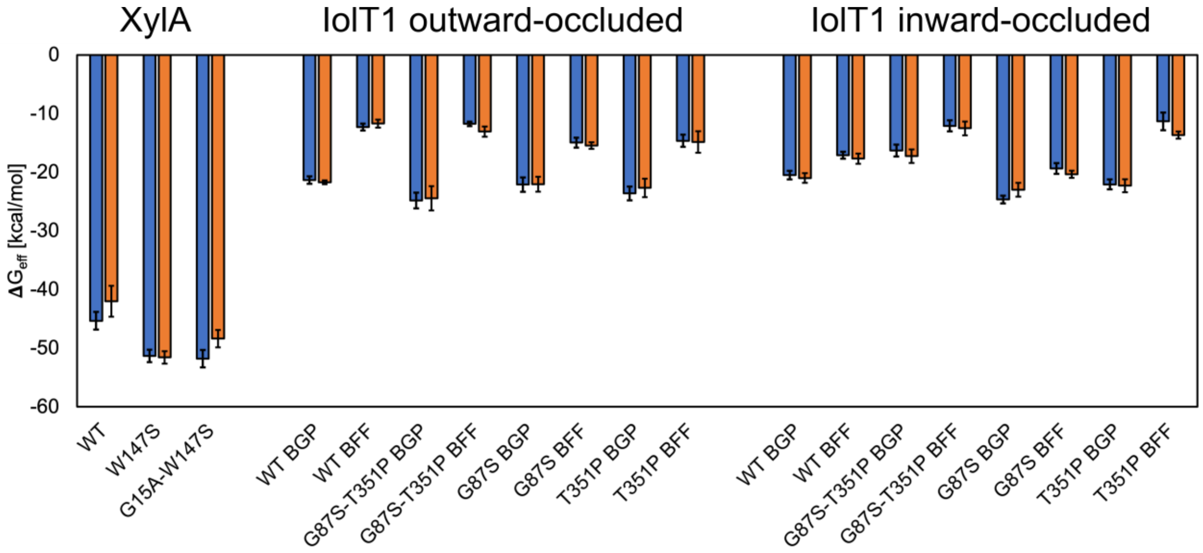

**Figure S29. Molecular Mechanics-Poisson Boltzmann Surface Area (MM-PBSA) binding effective energies computed for β-D-glucopyranose (BGP) in XylAXC and BGP or β-D-fructofuranose (BFF) in IolT1 are converged.** Blue bars describe the binding effective energy for the first half of the replicas, orange bars the results for the second half of the replicas, overall indicating convergence of the results. The error bars represent the propagated SEM.

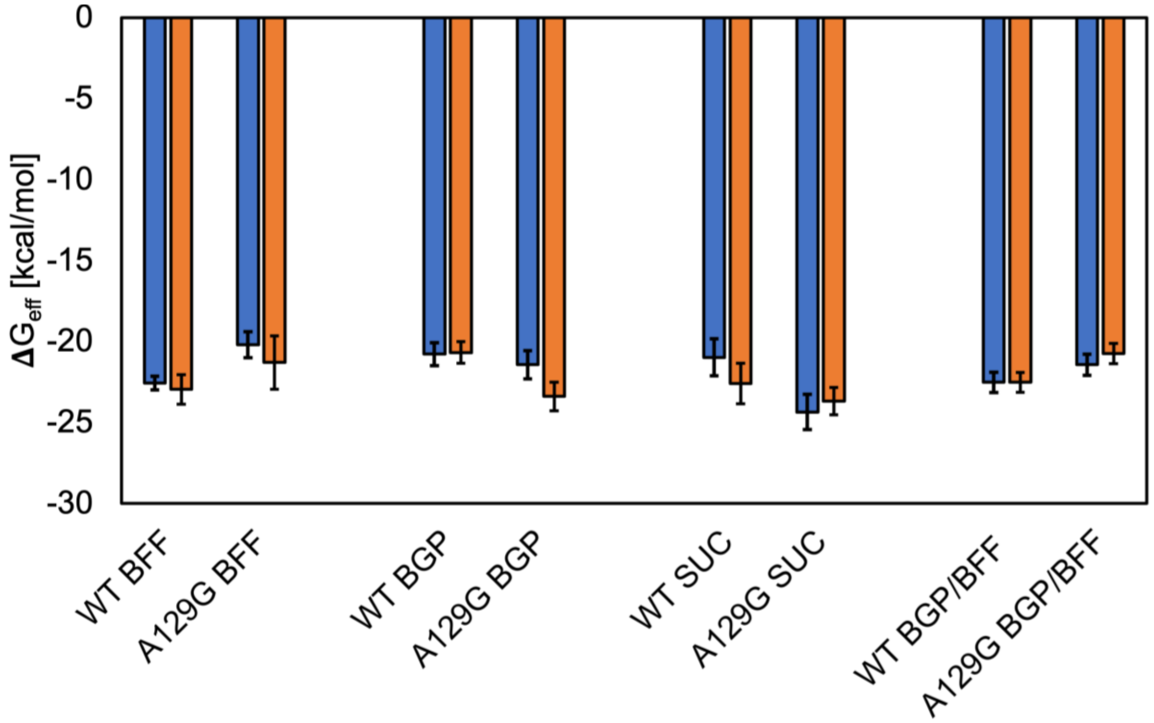

**Figure S30. Molecular Mechanics-Poisson Boltzmann Surface Area (MM-PBSA) binding effective energies computed for β-D-fructofuranose (BFF), β-D-glucopyranose (BGP), D-sucrose (SUC), and BGP/BFF in PtsS are converged.** Blue bars describe the binding effective energy for the first half of the replicas, orange bars the results for the second half of the replicas, overall indicating convergence of the results. The error bars represent the propagated SEM.

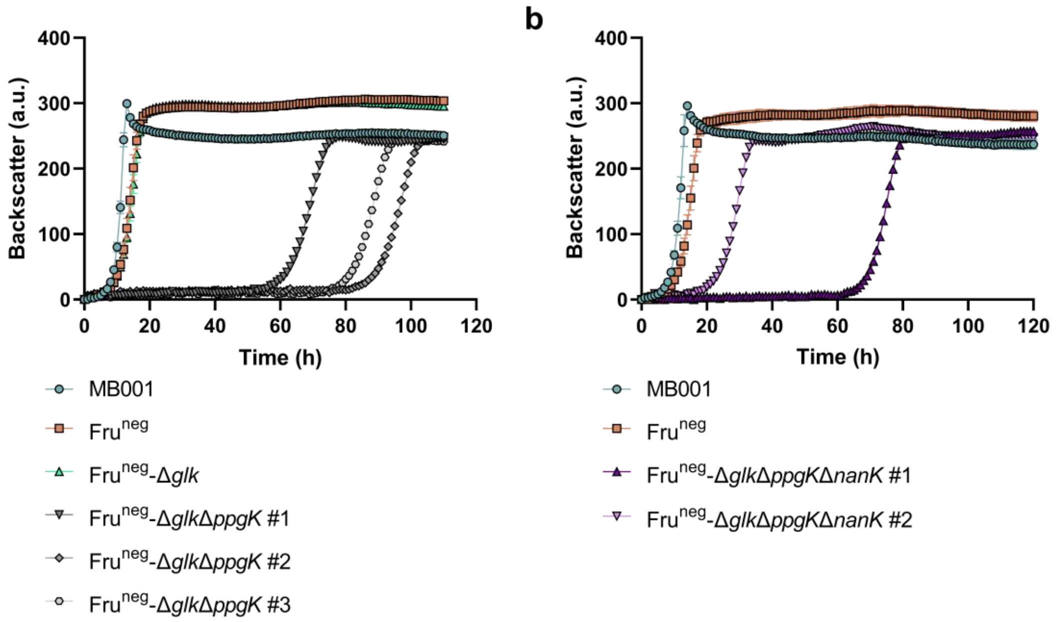

**Figure S31. Growth experiment of *C. glutamicum* strains Fru$^{neg}$-Δ*glk*Δ*ppgK* (a) and Fru$^{neg}$-Δ*glk*Δ*ppgK*Δ*nanK* (b) in CGXII medium with 40 g/L D-glucose.** The experiment was conducted in a BioLector II at 30°C and 1200 rpm. For each cultivation, the strains MB001 and Fru$^{neg}$ were included as controls and for (A) also the strain Fru$^{neg}$-Δ*glk*. Cultivations of MB001, Fru$^{neg}$ and Fru$^{neg}$-Δ*glk* are represented as mean ± S.D. from three biological replicates ($n$ = 3). Data points of Fru$^{neg}$-Δ*glk*Δ*ppgK* and Fru$^{neg}$-Δ*glk*Δ*ppgK*Δ*nanK* show three ($n$ = 3) and two individual cultivations ($n$ = 2), respectively.

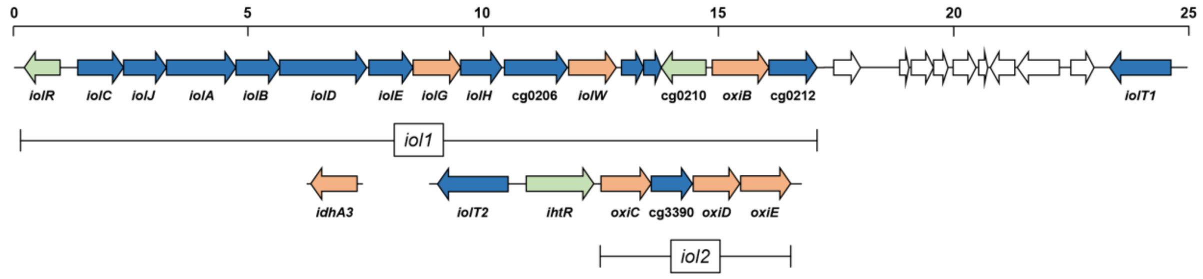

**Figure S32. Gene clusters of *C. glutamicum* involved in the metabolization of inositols.** Genes coding for inositol transporters and for enzymes involved in inositol metabolism are colored in blue, except for genes encoding inositol dehydrogenases, which are marked in orange. Genes encoding transcriptional regulators are shown in green. The regions named *iol1* and *iol2* as well as the *idhA3* gene were deleted in order to prevent oxidation of D-glucose to D-gluconate by a side activity of inositol dehydrogenases. The values of the scale on top are given in kilobases. The illustration was adapted from [4] and subjected to graphical adjustments.

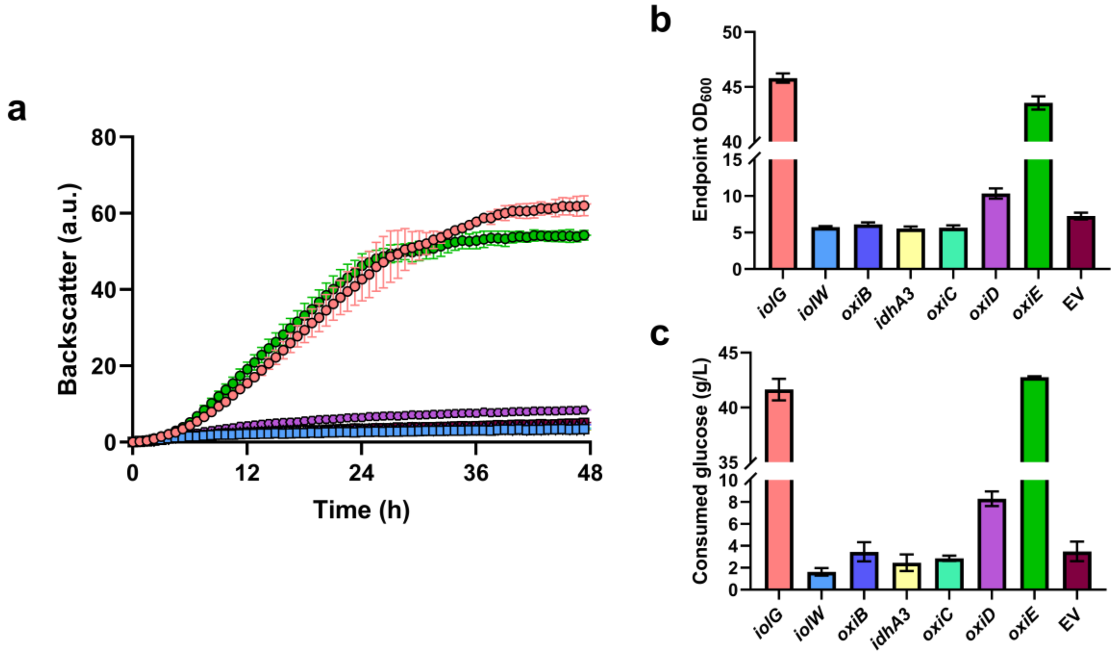

**Figure S33. Inositol dehydrogenase screening for D-glucose utilization in *C. glutamicum*.** The *C. glutamicum* Glu$^{neg}$ strain was transformed with pPREx2 expression plasmids, each encoding one of the seven annotated inositol dehydrogenases of *C. glutamicum* (*iolG*, *iolW*, *oxiB*, *idhA3*, *oxiC*, *oxiD* & *oxiE*) and the empty vector as control. The transformed strains were used for a growth experiment in CGXII medium with 40 g/L of D-glucose, 25 µg/mL kanamycin and 1 mM IPTG, which was performed at 30°C and 1200 rpm for 48 h in a BioLector I (**a**). After termination of the experiment, the endpoint $OD_{600}$ of the respective strains was determined (**b**). Supernatants of the cultures were used for HPLC measurement to determine D-glucose consumption (**c**). All data points represent mean ± S.D. from three biological replicates (*n* = 3), except for *iolG* (*n* = 2).

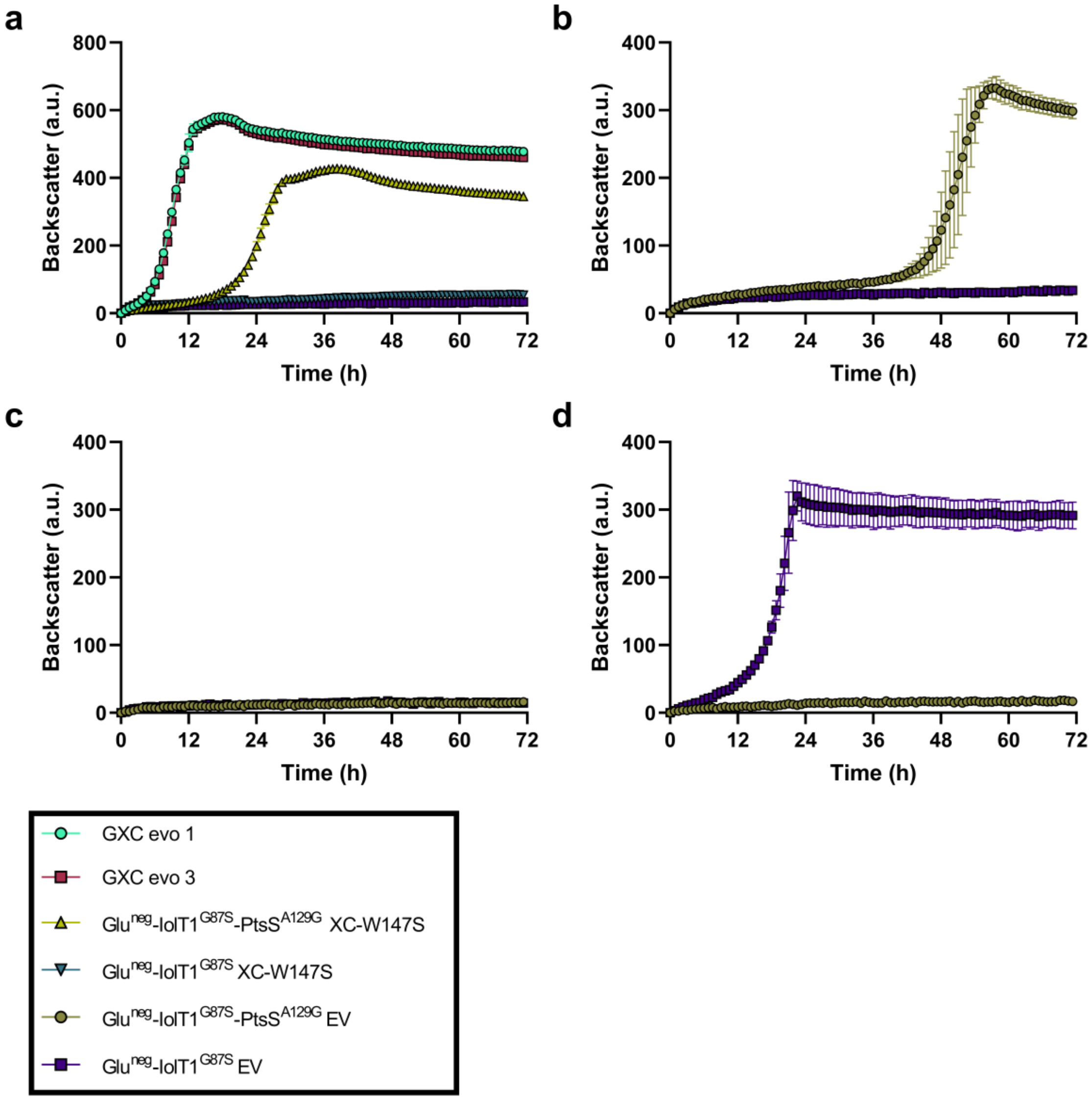

**Figure S34. Evaluation of reverse engineered GXC evo strains with pPREx2-*xylA*$_{XC}$-W147S (XC-W147S) or pPREx2 (EV) in D-glucose, D-fructose or D-sucrose minimal medium. a** GXC evo strains, Glu$^{neg}$-IolT1$^{G87S}$ strain containing PtsS-A129G (Glu$^{neg}$-IolT1$^{G87S}$-PtsS$^{A129G}$), and the unevolved Glu$^{neg}$-IolT1$^{G87S}$ strain carrying XC-W147S or EV were cultivated in CGXII medium with 40 g/L of D-glucose. **b-d** Growth experiments with strains Glu$^{neg}$-IolT1$^{G87S}$-PtsS$^{A129G}$ and Glu$^{neg}$-IolT1$^{G87S}$ carrying pPREx2 (EV) in CGXII medium with 40 g/L of D-glucose (**b**), D-fructose (**c**), or D-sucrose (**d**). All experiments were conducted at 30°C and 1200 rpm for 180 h. All data points represent mean ± S.D. from three biological replicates ($n$ = 3), except for the GXC evo strains ($n$ = 2).

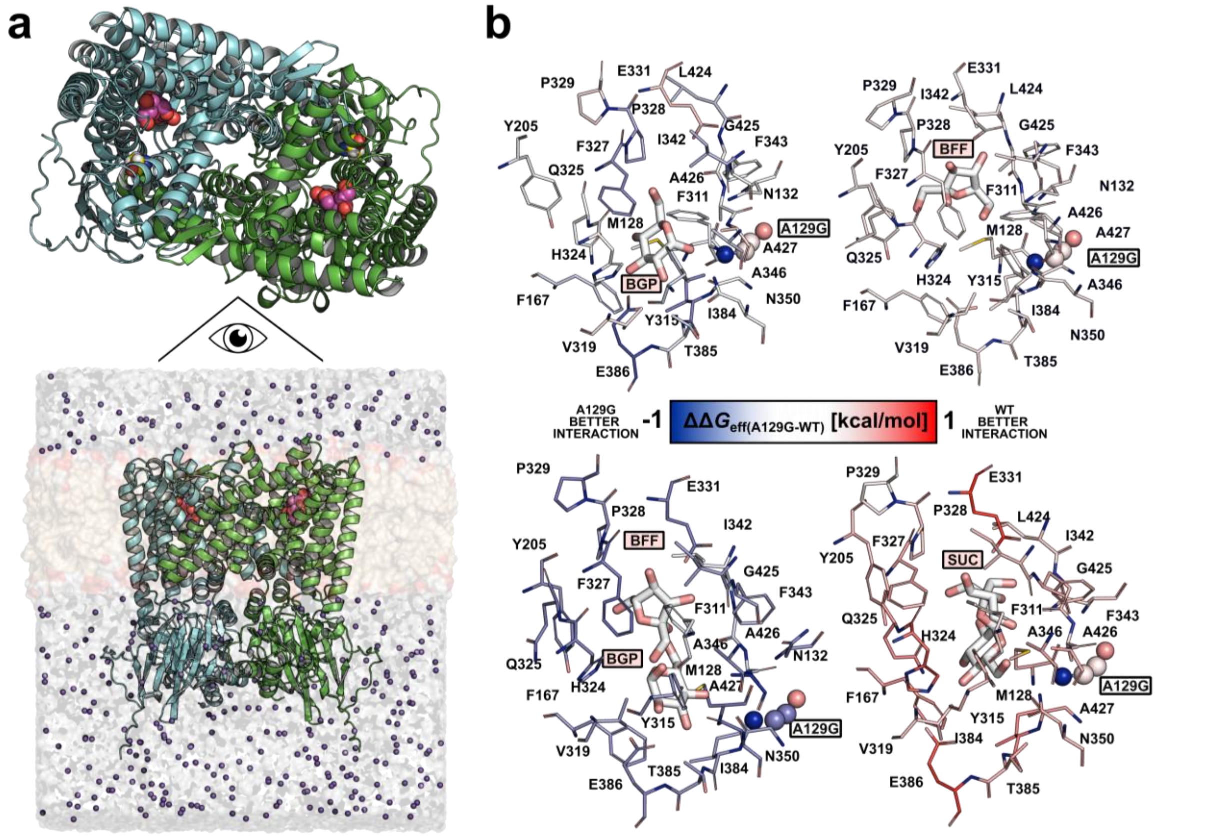

**Figure S35. Mutation A129G impacts carbohydrate binding to the PtsS homodimer. a** The homodimeric structure of *C. glutamicum* PtsS was predicted using AlphaFold3 [5]. The structure contains the carbohydrates β-D-glucopyranose (BGP), β-D-fructofuranose (BFF), BGP/BFF, or D-sucrose (SUC), respectively, in the binding site of each monomer. The carbohydrate is depicted as purple spheres and the mutagenesis site A129G is depicted as yellow spheres in the upper panel. **b** Differential per-residue effective binding energy ($\Delta\Delta G_{eff(A129G\text{-}WT)}$) mapped at the structural level. The energies were averaged across replicas. Larger changes in contributions to the binding effective energy occur near the BGP, SUC, or BGP/BFF binding site, but encompass residues up to 15 Å away from the carbohydrate. Residues for which the largest differences were found are depicted as sticks and colored according to the color scale; red (blue) colors indicate stronger binding by the wild type (variant). The mutagenesis site A129G is depicted as spheres.

**Supplemental Tables**

**Table S1. D-Glucose isomerases with reported activity for D-glucose or D-xylose at 30°C.** Parameters for D-glucose are highlighted in green, while parameters for D-xylose are shown in blue. N.d., not determined.

| Organism (UniProt accession number) | Temperature[a,b] | | $K_m$[a,b] (mM) [condition] | $v_{max}$[a,b] (U/mg) [condition] | $k_{cat}$[a,b] ($s^{-1}$) [condition] | $k_{cat}/K_m$[a,b] ($s^{-1}$ $mM^{-1}$) [condition] | Reference |
|---|---|---|---|---|---|---|---|
| | Optimum (°C) | Activity at 30 °C | | | | | |
| *Anoxybacillus gonensis* (M4HQI7) | 85 | ~ 20 % | 146 [85 °C, pH 6.5] | 43.72 [85 °C, pH 6.5] | 36.47 [85 °C, pH 6.5] | 0.25 [85 °C, pH 6.5] | [6] |
| *Anoxybacillus ayderensis* (A0A0D0HRN9) | 80 | ~ 25 % | 81 [80 °C, pH 7.5] | 2.24 [80 °C, pH 7.5] | n.d. | n.d. | [7] |
| *Arthrobacter* strain N.R.R.L. B3728 (P12070) | 70 | ~ 15 % | 225 [30 °C, pH 7] | 3.2 [30 °C, pH 7] | 2.32 [30 °C, pH 7] | 0.01 [30 °C, pH 7] | [8] |
| *Bacillus* sp. NCIM59 (n.a.) | 80 | ~ 20 % | 142 [80 °C, pH 7.5] | n.d. | 53 [80 °C, pH 7.5] | 0.37 [80 °C, pH 7.5] | [9] |
| *Bifidobacterium adolescentis* (A1A0H0) | 60 | 50% | 398 [45 °C, pH 7] | n.d. | n.d. | n.d. | [10] |
| *Burkholderia cenocepacia* (B4ENA5) | 37 | ~ 30 % | 17 [37 °C, pH 7.2] | 44.97 [37 °C, pH 7.2] | 0.32 [37 °C, pH 7.2] | 0.02 [37 °C, pH 7.2] | [11] |
| *Cereus pterogonus* (n.a.) | 80 | 60 % | 121 [80 °C, pH 7.5] | 7.75 [80 °C, pH 7.5] | n.d. | n.d. | [12] |
| *Escherichia coli* (P00944) | 45 | ~ 5 % | 220 [30 °C, pH 6.8] | 0.06 [30 °C, pH 6.8] | n.d. | n.d. | [13] |
| *Lactobacillus reuteri* (F8KFT7) | 65 | ~ 10 % | 1099 [60 °C, pH 5] | 4.6 [60 °C, pH 5] | 15.8 [60 °C, pH 5] | 0.014 [60 °C, pH 5] | [14] |
| *Opuntia vulgaris* (n.a.) | 90 | ~ 45 % | 260 [90 °C, pH 7.5] | 9.1 [90 °C, pH 7.5] | n.d. | n.d. | [15] |
| *Paenibacillus* sp. R4 (A0A4V8H014) | 60 | ~ 30 % | 17.7 [25 °C, pH 8] | n.d. | 8.5 [25 °C, pH 8] | 0.480 [25 °C, pH 8] | [16] |
| *Piromyces* sp. E2 (Q9P8C9) | n.d. | n.d. | 430 [30 °C, pH 7] | n.d. | 0.1 [30 °C, pH 7] | 0.0002 [30 °C, pH 7] | [17] |
| *Streptomyces albus* (P24299) | 75 | ~ 1-5 % | 86 [70 °C, pH 7.2] | 1.23 [70 °C, pH 7.2] | n.d. | n.d. | [18] |
| *Streptomyces murinus* (P37031) | 85 | ~ 5-10 % | 440 [80 °C, pH 7] | n.d. | 9.93 [80 °C, pH 7] | 0.022 [80 °C, pH 7] | [19] |
| *Streptomyces olivaceoviridis* (Q93RJ9) | 80 | ~ 1-5 % | n.d. | n.d. | n.d. | n.d. | [20] |
| *Streptomyces violaceoruber* (P14405) | 80 | ~ 5-10 % | 150 [35 °C, pH 7.5] | n.d. | 4.9 [35 °C, pH 7.5] | 0.033 [35 °C, pH 7.5] | [21] |
| *Thermoanaerobacterium sacchaorlyticum* (P30435) | 80 | ~ 1-5 % | 120 [65 °C, pH 7] | 6.3 [65 °C, pH 7] | 21 [65 °C, pH 7] | 0.175 [65 °C, pH 7] | [22] |

**Table S2. IolT1-membrane systems generated with Packmol-Memgen and subjected to MD simulations and analyses.** The IolT1 variants G87S, T351P, and G87S-T351P were compared to wild-type IolT1. Ligands used are β-D-glucopyranose (BGP) and β-D-fructofuranose (BFF). The change in the folding free energy with respect to wild-type IolT1 $\Delta\Delta G = \Delta G_{variant} - \Delta G_{wild\ type}$ is measured in kcal $mol^{-1}$ and calculated with FoldX [23].

| IolT1 State | Template | TopScore[a] | Mutation structural stability (Δ*ΔG*) | Ligand |
|---|---|---|---|---|
| Outward Open (I) | 4ZWC *H. sapiens* | 0.25 | WT | - |
| Outward Open (I) | 4ZWC *H. sapiens* | 0.25 | G87S (-0.54) | - |
| Outward Open (I) | 4ZWC *H. sapiens* | 0.25 | T351P (2.47) | - |
| Outward Open (I) | 4ZWC *H. sapiens* | 0.25 | G87S-T351P (0.97) | - |
| Outward Occluded (II) | 4ZW9 *H. sapiens* | 0.27 | WT | BGP |
| Outward Occluded (II) | 4ZW9 *H. sapiens* | 0.27 | G87S (-0.68) | BGP |
| Outward Occluded (II) | 4ZW9 *H. sapiens* | 0.27 | T351P (2.88) | BGP |
| Outward Occluded (II) | 4ZW9 *H. sapiens* | 0.27 | G87S-T351P (1.10) | BGP |
| Outward Occluded (II) | 4ZW9 *H. sapiens* | 0.27 | WT | BFF |
| Outward Occluded (II) | 4ZW9 *H. sapiens* | 0.27 | G87S (-0.67) | BFF |
| Outward Occluded (II) | 4ZW9 *H. sapiens* | 0.27 | T351P (2.85) | BFF |
| Outward Occluded (II) | 4ZW9 *H. sapiens* | 0.27 | G87S-T351P (1.09) | BFF |
| Inward Occluded (III) | 4JA3 *E. coli* | 0.30 | WT | BGP |
| Inward Occluded (III) | 4JA3 *E. coli* | 0.30 | G87S (1.18) | BGP |
| Inward Occluded (III) | 4JA3 *E. coli* | 0.30 | T351P (2.85) | BGP |
| Inward Occluded (III) | 4JA3 *E. coli* | 0.30 | G87S-T351P (2.02) | BGP |
| Inward Occluded (III) | 4JA3 *E. coli* | 0.30 | WT | BFF |
| Inward Occluded (III) | 4JA3 *E. coli* | 0.30 | G87S (1.17) | BFF |
| Inward Occluded (III) | 4JA3 *E. coli* | 0.30 | T351P (2.83) | BFF |
| Inward Occluded (III) | 4JA3 *E. coli* | 0.30 | G87S-T351P (2.02) | BFF |
| Inward Open (IV) | 4YB9 *B. taurus* | 0.36 | WT | - |
| Inward Open (IV) | 4YB9 *B. taurus* | 0.36 | G87S (0.35) | - |
| Inward Open (IV) | 4YB9 *B. taurus* | 0.36 | T351P (2.28) | - |
| Inward Open (IV) | 4YB9 *B. taurus* | 0.36 | G87S-T351P (1.23) | - |

[a] Score to assess the quality of a protein model, bounded between 0 and 1. Lower values indicate a higher quality.

**Table S3. Mutations in plasmid pPREx2-*xylA*$_{XC}$-W147S after random mutagenesis and selection of better growing Fru$^{neg}$-IolT1$^{G87S}$ clones in D-fructose minimal medium.**

| Clone # | Missense mutations in XylA$_{XC}$-W147S | Mutations in pPREx2 backbone |
|---|---|---|
| **5** | V5F | - |
| **6** | A9G | - |
| **7** | T427M | - |
| **8** | - | T → A, 23 bp upstream of *xylA* start codon |
| **9** | V5F | - |
| **12** | - | C → T, 21 bp upstream of *xylA* start codon |
| **13** | S2N<br>A9T<br>A37P<br>A47D | - |
| **14** | V5I | - |
| **15** | S2N<br>W390R | - |
| **16** | S2T | - |
| **17** | S2N<br>L299M | - |
| **18** | S2N<br>E124K | - |
| **19** | - | C → T, 3 bp upstream of *xylA* start codon |
| **20** | S2T | - |
| **21** | G15A | - |
| **23** | A421V | T → G, 18 bp upstream of *xylA* transcription start site |
| **25** | R24A<br>S286T | - |
| **26** | - | T → G, 27 bp upstream of *xylA* start codon |
| **27** | S2T | - |
| **28** | S2T | - |
| **29** | S2N | - |
| **30** | R24A<br>S286T | - |

**Table S4. Mutations identified for evolved strains of *C. glutamicum* Glu$^{neg}$-IolT1$^{G87S}$ pPREx2-*xylA*$_{XC}$-W147S (GXC evo).**

| Strain | Location | Mutation |
|---|---|---|
| GXC evo 1 | *ptsS* (cg2925) | Missense mutation: A129G |
| GXC evo 2 | *ptsS* (cg2925) | Missense mutation: A129G |

**Table S5. List of oligonucleotides used in this study.**

| Oligonucleotide name | Sequence (5'→3') |
|---|---|
| Construction of pK19*mobsacB* plasmids | |
| ALP070_*ptsF*_FW1 | CCTGCAGGTCGACTCTAGAGGACCACAACTTTCAGGTGGTAAC |
| ALP071_*ptsF*_RV1 | GCCGGACAAGCGAGGAATTATTTAC |
| ALP072_*ptsF*_FW2 | GTAAATAATTCCTCGCTTGTCCGGCGCAGCTGTAAACGCATAATC |
| ALP073_*ptsF*_RV2 | AAAACGACGGCCAGTGAATTCTAACGGTGAGCTGCCAACATGAG |
| ALP078_*ptsG*_FW1 | CCTGCAGGTCGACTCTAGAGCGACCCATCCAAGTACGGAATG |
| ALP079_*ptsG*_RV1 | GTCGTCGTCAGTTTGGACGC |
| ALP080_*ptsG*_FW2 | GCGTCCAAACTGACGACGACGTCAACGGCAAGAACGAGTAAC |
| ALP081_*ptsG*_RV2 | AAAACGACGGCCAGTGAATTCAGAAGTACCAGATCAGAG |
| ALP086_*glk*_FW1 | CCTGCAGGTCGACTCTAGAGCAACCATACCGTTCTGCCACC |
| ALP087_*glk*_RV1 | CTGTAGTGGAAGCCAACTAGG |
| ALP088_*glk*_FW2 | CCTAGTTGGCTTCCACTACAGGGCCGGTTTTTGTGGCATAAG |
| ALP089_*glk*_RV2 | AAAACGACGGCCAGTGAATTGGTTATTGCCTCCACTCATGGG |
| ALP094_*ppgK*_FW1 | CCTGCAGGTCGACTCTAGAGCCTCTTGTCATACTGCCAGGTC |
| ALP095_*ppgK*_RV1 | CCACCGATATCAATTCCAAATCC |
| ALP096_*ppgK*_FW2 | GGATTTGGAATTGATATCGGTGGCCAACACCTCACCCCATAAG |
| ALP097_*ppgK*_RV2 | AAAACGACGGCCAGTGAATTCGACGCCGTCTTCATCTTC |
| ALP143_*nanK*_FW1 | CCTGCAGGTCGACTCTAGAGTTCTCACCCGCACTCGTTCC |
| ALP144_*nanK*_RV1 | GGTGCAAGTGGGATCAGTCATGAG |
| ALP145_*nanK*_FW2 | CTCATGACTGATCCCACTTGCACCGCCCGCGATAACGCCTTTTAAGC |
| ALP146_*nanK*_RV2 | AAAACGACGGCCAGTGAATTGTTGGGCTCTTAGAAGCGATTCTG |
| ALP255_T351P_FW | CCTGCAGGTCGACTCTAGAGGTCGCATTTCCGACAACTG |
| ALP256_T351P_RV | AAAACGACGGCCAGTGAATTGCTTGAAGCGCCACAAATGC |
| ALP336_G87S_FW | CCTGCAGGTCGACTCTAGAGATGGCTAGTACCTTCATTCAG |
| ALP337_G87S_RV | AAAACGACGGCCAGTGAATTATCGCATTGATGACAAAAGC |
| ALP478_*ptsS*_FW1 | CCTGCAGGTCGACTCTAGAGCTATACCGCTCGACAGATCTTG |
| ALP479_*ptsS*_RV1 | AATGTCGCGCAGGATGCGTTG |
| ALP480_*ptsS*_FW2 | CAACGCATCCTGCGCGACATTTAAGTTGAAACCTTGAGTGTTCG |
| ALP481_*ptsS*_RV2 | AAAACGACGGCCAGTGAATTCTGCTTTAGCATTTCCGGCACAG |
| ALP486_*ptsS*-A129G_FW | CCTGCAGGTCGACTCTAGAGATGGACCATAAGGACCTCGCGCAACG |
| ALP491_*ptsS*-A129G_RV | AAAACGACGGCCAGTGAATTACTGGAGTGATCAGGAAGTC |
| Construction of pPREx2 plasmids | |
| ALP118_*xylA*(XC)_FW | CCTGCAGAAGGAGATATACAATGAGCAACACCGTTTTC |
| ALP121_*xylA*(XC)_RV | CTGTGGGTGGGACCAGCTAGCTTTCTCCTCTCAACGCGTCAG |
| ALP125_*xylA*(AG)_FW | CCTGCAGAAGGAGATATACAATGGCCTACTTCGAAAACGTGG |
| ALP126_*xylA*(AG)_RV | CTGTGGGTGGGACCAGCTAGTTAGCGAGCCACGCACACTTCC |
| ALP128_*xylA*(PE2)_FW | CCTGCAGAAGGAGATATACAATGGCCAAAGAATACTTCCC |
| ALP129_*xylA*(PE2)_RV | CTGTGGGTGGGACCAGCTAGTTACTGGTACATTGCCACGATTGC |
| ALP131_*xylA*(PR4)_FW | CCTGCAGAAGGAGATATACAATGGGCTACTTCGATCACGTGGG |
| ALP132_*xylA*(PR4)_RV | CTGTGGGTGGGACCAGCTAGTTACACGTTGATGATGTACTGGTTCAGG |

| | |
|---|---|
| ALP134_*xylA*(EC)_FW | CCTGCAGAAGGAGATATACAATGCAAGCCTATTTTGACCAGCTC |
| ALP135_*xylA*(EC)_RV | CTGTGGGTGGGACCAGCTAGTTATTTGTCGAACAGATAATGATTTACC |
| ALP137_*xylA*(AB)_FW | CCTGCAGAAGGAGATATACAATGTCCGTGCAGCCAACTCC |
| ALP138_*xylA*(AB)_RV | CTGTGGGTGGGACCAGCTAGTTAGCGGGAGCCCAGGAGGTGTTCGATTG |
| ALP140_*xylA*(SM)_FW | CCTGCAGAAGGAGATATACAATGTCCTTCCAGCCAACTCCAG |
| ALP141_*xylA*(SM)_RV | CTGTGGGTGGGACCAGCTAGTTAACCGCGTGCACCCAGGAG |
| ALP196_*dae*(AT)_FW | CCTGCAGAAGGAGATATACATATGAAGCACGGCATCTACTACTCC |
| ALP197_*dae*(AT)_RV | CTGTGGGTGGGACCAGCTAGTTAGCCACCGAGCACGAAGC |
| ALP200_*dae*(AG)_FW | CCTGCAGAAGGAGATATACATATGAAGATCGGCTGCCACG |
| ALP201_*dae*(AG)_RV | CTGTGGGTGGGACCAGCTAGTTAGTGCAGTTCGATGGTCTTG |
| ALP204_*dte*(PC)_FW | CCTGCAGAAGGAGATATACATATGAACAAGGTGGGCATGTTCTAC |
| ALP205_*dte*(PC)_RV | CTGTGGGTGGGACCAGCTAGTTATGCCAGCTTATCGCGCACGAAC |
| ALP223_AB(c_t)_FW | AAATGCATGCCGCTTCGCCTTC |
| ALP224_AB(c_t)_RV | GGACATATGTATATCTCCTTCTGCAGGCATGCAAGCTTGG |
| ALP248_AB(S2F)_FW | GGAGATATACATATGTTCGTGCAGCCAACTCC |
| ALP249_AB(S2F)_RV | GGAGTTGGCTGCACGAACATATGTATATCTCC |
| ALP359_RM_XC_FW | GCCTGCAGAAGGAGATATACAT |
| ALP360_RM_XC_RV | GTGGGACCAGCTAGCTTTCTCCTC |
| ALP368_*xylA*(XC)-*dae*(AT)_RV | CAGGTCAGCTGCAAAGGTGCTCAACGCGTCAGGTACTGATTG |
| ALP369_*xylA*(XC)-*dae*(AT)_FW | GCACCTTTGCAGCTGACCTGAAGGAGATATACATATGAAGCACGGCATCTACTAC |
| ALP429_G15A_RV1 | CAGAAGCTGTGCCAGTAGG |
| ALP430_G15A_FW2 | GCCTACTGGCACAGCTTCTG |
| ALP431_E124K_FW | CGGACGACATCGGCGAGTACAAAAACAACCTCAAGCACATGGT |
| ALP432_E124K_RV | ACCATGTGCTTGAGGTTGTTTTTGTACTCGCCGATGTCGTCCG |
| ALP433_S286T_FW | ACGCTGTCGGGCCACACCTTCGAGCACGACCTG |
| ALP434_S286T_RV | CAGGTCGTGCTCGAAGGTGTGGCCCGACAGCGT |
| ALP435_L299M_FW | CAGCGATGCCGGCATGCTCGGCAGCATCG |
| ALP436_L299M_RV | CGATGCTGCCGAGCATGCCGGCATCGCTG |
| ALP437_A421V_RV | CTGGTGCCGTTGGCAAAGTC |
| ALP438_A421V_FW | GACTTTGCCAACGGCACCAG |
| Construction of pPREx5 plasmids | |
| ALP355_His-XC_FW | ACCTGTATTTTCAGGGCCATATGAGCAACACCGTTTTCATC |
| ALP356_His-XC_RV | CTGTCCACCAGTCATGCTAGTCAACGCGTCAGGTACTGATTG |

**Table S6. XylA$_{XC}$ systems generated and subjected to MD simulations and analyses. Ligand used is β-D-glucopyranose (BGP).** The change in the folding free energy with respect to the wild-type XylA$_{XC}$ $\Delta\Delta G = \Delta G_{variant} - \Delta G_{wild\ type}$ is measured in kcal mol$^{-1}$ and calculated with FoldX [23].

| XylA State | Template used to extract BGP coordinates | AF3 pTM score[a] | Mutation structural stability (ΔΔ*G*) | Ligand |
|---|---|---|---|---|
| XylA homodimer (I) | 4LNC *S. rubiginosus* | 0.82 | WT | BGP |
| XylA homodimer (I) | 4LNC *S. rubiginosus* | 0.82 | W147S (-0.94) | BGP |
| XylA homodimer (I) | 4LNC *S. rubiginosus* | 0.82 | G15A-W147S (1.80) | BGP |

[a] Measure of AlphaFold 3 of the prediction accuracy of a complex structure, representing the predicted TM score for a superposition between the predicted complex structure and the assumed true structure, bounded between 0 and 1. Higher values indicate a higher accuracy.

**Table S7. Mutations of *C. glutamicum* Fru$^{neg}$-Δ*glk*Δ*ppgK* and Fru$^{neg}$-Δ*glk*Δ*ppgK*Δ*nanK* suppressor mutants that were able to grow on D-glucose.**

| Strain | Location | Mutation |
|---|---|---|
| Fru$^{neg}$-Δ*glk*Δ*ppgK* #1 | *iolR* (cg0196) | Insertion of cytosine after 483 nt → frameshift after amino acid 161 |
| Fru$^{neg}$-Δ*glk*Δ*ppgK* #2 | *ihtR* (cg3388) | Insertion of guanine after 762 nt → frameshift after amino acid 254 |
| Fru$^{neg}$-Δ*glk*Δ*ppgK* #3 | *ihtR* (cg3388) | Insertion of transposase element (cg2600) |
| Fru$^{neg}$-Δ*glk*Δ*ppgK*Δ*nanK* #1 | *iolR* (cg0196) | Insertion of cytosine after 483 nt → frameshift after amino acid 161 |
| Fru$^{neg}$-Δ*glk*Δ*ppgK*Δ*nanK* #2 | *ihtR* (cg3388) | Missense mutation: L291R |

**Table S8. PtsS systems generated and subjected to MD simulations and analyses.** Ligands used are β-D-glucopyranose (BGP) and β-D-fructofuranose (BFF). The change in the folding free energy of the PtsS-A129G variant with respect to the wild-type PtsS $\Delta\Delta G = \Delta G_{\text{variant}} - \Delta G_{\text{wild type}}$ is measured in kcal $mol^{-1}$ and calculated with FoldX [23].

| XylA state | Template used to extract BGP coordinates | AF3 pTM score[a] | Mutation structural stability ($\Delta\Delta G$) | Ligand |
|---|---|---|---|---|
| PtsS homodimer (I) | 3QNQ *B. cereus* | 0.57 | WT | - |
| PtsS homodimer (I) | 3QNQ *B. cereus* | 0.57 | A129G (2.73) | - |
| PtsS homodimer (I) | 3QNQ *B. cereus* | 0.57 | WT | BGP |
| PtsS homodimer (I) | 3QNQ *B. cereus* | 0.57 | A129G (2.73) | BGP |
| PtsS homodimer (I) | 3QNQ *B. cereus* | 0.57 | WT | BFF |
| PtsS homodimer (I) | 3QNQ *B. cereus* | 0.57 | A129G (2.73) | BFF |
| PtsS homodimer (I) | 3QNQ *B. cereus* | 0.57 | WT | SUC |
| PtsS homodimer (I) | 3QNQ *B. cereus* | 0.57 | A129G (2.73) | SUC |
| PtsS homodimer (I) | 3QNQ *B. cereus* | 0.57 | WT | BGP/BFF |
| PtsS homodimer (I) | 3QNQ *B. cereus* | 0.57 | A129G (2.73) | BGP/BFF |

[a] Measure of AlphaFold 3 of the prediction accuracy of a complex structure, representing the predicted TM score for a superposition between the predicted complex structure and the assumed true structure, bounded between 0 and 1. Higher values indicate a higher accuracy.

**Table S9. Structural influence of the A129G amino acid exchange in PtsS on binding site tunnel characteristics compared to wild-type PtsS calculated with CAVER.** Data was calculated with a probe radius of 1.2 Å.

| PtsS System | Average bottleneck radius [Å] | Average length [Å] |
|---|---|---|
| WT | 2.13 | 11.21 |
| A129G | 2.02 | 11.37 |

### Supplemental Notes

#### Supplemental Note 1. Directed evolution of XylA$_{XC}$-W147S

To further improve D-glucose isomerase activity of XylA$_{XC}$-W147S, the corresponding gene was subjected to two rounds of random mutagenesis, conducted via the GeneMorph II Random Mutagenesis Kit (Agilent Technologies, Santa Clara, USA). The resulting variants were cloned into pPREx2 and used for *E. coli* DH5α transformation. The transformants were used to isolate a plasmid library for the transformation of *C. glutamicum* Fru$^{neg}$-IolT1$^{G87S}$. The transformed cells were plated on D-fructose CGXII agar plates (Figure S15A). In total, 30 clones were obtained with a larger colony size compared to the control with non-mutated pPREx2-*xylA*$_{XC}$-W147S. Of these 30 clones, 22 showed faster growth in liquid D-fructose minimal medium (Figure S17). Plasmid isolation and sequencing revealed mutations upstream of *xylA*$_{XC}$-W147S for five clones and one or multiple missense mutations within the *xylA*$_{XC}$-W147S coding region for 18 clones. In total, 16 different missense mutations were obtained, the most frequent being S2N and S2T exchanges, each of which occurred at least four times, followed by two V5F and one V5I exchange (Table S3).

Excluding the S2 and V5 mutations, we individually tested seven of the missense mutations in combination with W147S in XylA$_{XC}$ for improved growth of Fru$^{neg}$-IolT1$^{G87S}$ in D-fructose minimal medium. Six of the seven mutations (R24A, E124K, S286T, L299M, A421V, T427M) barely affected growth, while the G15A mutation improved growth moderately (Figure S15B-C). Since G15A is located in the N-terminal region, a translation-enhancing effect leading to increased XylA$_{XC}$ protein levels could be responsible for its positive effects on growth, similar to the S2F mutation found in the XylA$_{AB}$ protein (Figure S13A). In a Western blot analysis, the XylA$_{XC}$-G15A-W147S variant revealed a stronger protein signal compared to the XylA$_{XC}$-W147S variant (Figure S18). The protein XylA$_{XC}$-G15A-W147S was overproduced, purified, and kinetically characterized (Table 3).

**Supplemental Note 2. Molecular dynamics simulations of the XylA$_{XC}$ variants**

All-atom unbiased MD simulations were performed of the structural homodimer model of XylA$_{XC}$ bound to D-glucose (as β-D-glucopyranose) (Figure 5A, Figure S25, Table S6). Starting structures of the variants W147S and G15A-W147S were generated using the same protocol as for the IolT1 mutants. For the two systems, five replicas were simulated, each for 2 µs in length. MM-PBSA calculations were performed to provide qualitative insight into the relative effects of mutations on ligand binding and protein stability, calculating an effective binding energy ($\Delta G_{eff}$). As emphasized by Genheden & Ryde (2015) and Roux & Chipot (2024), the end-point free energy approximation is not recommended for absolute binding predictions or lead optimization. Here, it is solely used to evaluate trends across structurally similar wildtype and mutant protein-ligand complexes, with extended trajectories and multiple replicas indicating a robust comparative analysis. Effective binding energy computations revealed that W147S favors the interactions of negatively charged residues with D-glucose-$Mg^{2+}$ and disfavors interactions with polar and positively charged residues (Figure 5B). Here, $Mg^{2+}$ was considered part of the ligand as $Mg^{2+}$ has been described as an activator binding together with the carbohydrate [24]. Tryptophan can be involved in carbohydrate-aromatic interactions [25,26], likely via C-H···π interactions [27,28], whereas serine is less frequently found near carbohydrates [25,26]. The distance of W147 to the carbohydrate atom is ≥ 4 Å. As to position 147 only, the W to S exchange favors interactions with the carbohydrate only slightly (Figure 5B). Yet, overall, the W147S substitution favors the binding of D-glucose-$Mg^{2+}$ compared to wild-type XylA$_{XC}$. The increased affinity of XylA$_{XC}$-W147S to D-glucose probably favors the catalytic efficiency of the reversible conversion to D-fructose. In contrast, the double substitution G15A-W147S does not markedly change the effective binding energy of D-glucose-$Mg^{2+}$ to XylA$_{XC}$ compared to the W147S single mutation (Figure 5C).

**Supplemental Note 3. Construction of a D-glucose-negative *C. glutamicum* strain**

Initial work in the construction of the *C. glutamicum* $Glu^{neg}$ strain focused on the deletion of the glucokinase genes *glk* and *ppgK*, and the *nanK* gene encoding *N*-acetylmannosamine kinase NanK [29], which also accepts D-glucose as substrate [30], in *C. glutamicum* $Fru^{neg}$. A growth experiment in D-glucose minimal medium revealed the appearance of suppressor mutants of the strains $Fru^{neg}$-Δ*glk*Δ*ppgK* and $Fru^{neg}$-Δ*glk*Δ*ppgK*Δ*nanK* that were able to utilize D-glucose after a prolonged lag phase (Figure S31). Genome sequencing of three or two independently grown suppressor clones of both strains revealed that each one contained either a mutation in the *iolR* gene (cg0196) or in the *ihtR* gene (cg3388) (Table S7). In the case of *iolR*, two identical mutations were identified, i.e. a cytosine insertion after nucleotide 483 causing a frameshift after codon 161. In the case of *ihtR*, tree different mutations were found, i.e. a guanine insertion after nucleotide 762 causing a frameshift after codon 254, a transposase insertion, and a missense mutation causing an L291R exchange. Both *iolR* and *ihtR* code for transcriptional regulators and are located in gene clusters that are involved in inositol metabolization (Figure S32). IolR represses the expression of the genes ranging from *iolC* (cg0197) to *iolW* (cg0207) and also *iolT1* (cg0223) in the absence of *myo*-inositol (Figure S32) [31]. IhtR was characterized as a repressor linked to increased indole resistance via the regulation of its upstream gene *rhcM2* (cg3386) [32]. Experiments in our lab indicated that IhtR also represses the gene cluster *oxiC*-cg3390-*oxiD*-*oxiE* located downstream of *ihtR* (unpublished data).

The mutations identified in *iolR* and *ihtR* are supposed to lead to inactive variants of these regulators, causing derepression of their target genes. These include several genes encoding $NAD^+$-dependent inositol dehydrogenases, such as IolG, OxiE, and OxiD [33]. The IolG homolog of *Bacillus subtilis* was shown to oxidize not only *myo*-inositol to 2-keto-*myo*-inositol, but also D-glucose to D-gluconate with reasonable activity [34]. Such a side activity could explain the growth of our suppressor mutants on D-glucose: after oxidation of D-glucose by one or several of the induced inositol dehydrogneases, D-gluconate serves as a good growth substrate for *C. glutamicum* [35]. In order to prevent this type of D-glucose metabolization, we deleted the gene clusters *iol1* and *iol2* and additionally *idhA3* (cg2313), encoding another inositol dehydrogenase [33], in strain $Fru^{neg}$-Δ*glk*Δ*ppgK*Δ*nanK,* resulting in the strain $Glu^{neg}$. During 100 h incubation in D-glucose minimal medium, the $Glu^{neg}$ strain showed no growth (Figure S19). However, growth on D-glucose was enabled by plasmid-based expression of *iolG*, *oxiE*, or *oxiD*, supporting that the corresponding inositol dehydrogenases are able to oxidize D-glucose to D-gluconate (Figure S33).

**Supplemental Note 4. Further studies on PtsS-A129G**

The A129G mutation in PtsS was reconstructed in strain Glu$^{neg}$-IolT1$^{G87S}$ expressing *xylA*$_{XC}$-W147S and investigated for its growth impact via cultivation in the BioLector system. In D-glucose minimal medium, the mutation was confirmed to be beneficial for growth, although it could not fully restore the growth rate of the evolved strains (Figure S34A). Remarkably, the A129G mutation in PtsS also enabled growth of strain Glu$^{neg}$-IolT1$^{G87S}$-PtsS$^{A129G}$ carrying the pPREx2 control in D-glucose minimal medium, while it disabled its growth in D-sucrose minimal medium (Figure S34B and S34D). In D-fructose minimal medium, the PtsS-A129G mutation did not affect growth (Figure S34C).

To study the structural effect of the A129G mutation in PtsS on the geometry of the catalytic tunnel and on carbohydrate binding we performed all-atom unbiased MD simulations [36] of the structural homodimer model of PtsS in the apo state and bound to D-glucose (as β-D-glucopyranose), D-fructose (as β-D-fructofuranose), D-glucose with D-fructose, and D-sucrose (Table S8). The mutation site A129G is ~10 Å away from the center of mass (COM) of the D-sucrose moiety (Figure S35). Starting structures of the variant A129G were generated using the same protocol as for the IolT1 and the XylA$_{XC}$ mutants. For the two obtained systems, five replicas were simulated, each for 2 µs in length. A CAVER analysis [37] revealed that the mutation A129G does not markedly change the geometry of the catalytic tunnel when the tunnel is empty (Table S9). On the other hand, effective binding energy computations with MM-PBSA revealed that A129G favors interactions of the residues of the catalytic tunnel with D-glucose and D-glucose-D-fructose, whereas it does not affect D-fructose binding and disfavors D-sucrose binding (Figure S35B). Again here, MM-PBSA calculations were performed to provide qualitative insight into the relative effects of mutations on ligand binding and protein stability, calculating an effective binding energy ($\Delta G_{eff}$). For further details, see Supplemental Note 2. Based on these results, we assume that wild-type PtsS has a limited ability to cotransport D-glucose with D-fructose, which is improved upon the introduction of A129G. In presence of D-glucose isomerases, taken up D-glucose from D-glucose minimal medium is converted intracellularly to D-fructose, secreted back into the supernatant and taken up again with D-glucose via PtsS, which results into growth. However, since the introduction of A129G in PtsS also improved growth in D-glucose minimal medium without the presence of a D-glucose isomerase, the mutation might also confer the ability to cotransport D-glucose without the necessity of D-fructose.

### Supplemental Methods

#### Random mutagenesis and selection via growth

Random mutagenesis was performed by using the GeneMorph II Random Mutagenesis Kit (Agilent Technologies, Santa Clara, USA) on *xylA*$_{XC}$ containing the W147S mutation. Conditions were framed to obtain a low mutation frequency of 0-4.5 mutations/kb by using 1000 ng of DNA template amplified with 30 PCR cycles. The resulting PCR products were purified using the DNA Clean & Concentrator Kit (Zymo Research, Irvine, USA) and cloned via Gibson assembly into pPREx2 plasmid digested with NdeI and NheI [38]. Assembled plasmids were used for the transformation of *E. coli* DH5α and cells were plated on LB agar with 50 µg/mL kanamycin and incubated at 37°C for 24 h. The colonies formed were jointly resuspended in LB medium, which was then used for plasmid preparation via the Monarch Plasmid Miniprep Kit (New England Biolabs, Ipswich, USA). Purified plasmids were used for transformation of *C. glutamicum* Fru$^{neg}$-IolT1-G87S and cells were plated on CGXII agar plates with 40 g/L D-fructose, 25 µg/mL kanamycin and 25 µM IPTG. After 5 days of incubation at 30°C, larger sized clones were selected for a growth experiment in the BioLector.

#### High-cell-density biotransformation with Suc$^{neg}$-IolT1$^{G87S}$

The high-cell-density biotransformation was performed in 2 L DASGIP bioreactors (Eppendorf SE, Hamburg, Germany). Precultures of the Suc$^{neg}$-IolT1$^{G87S}$ strain carrying pPREx2-*xylA*$_{XC}$-G15A-W147S-*dae*$_{AT}$ were prepared first in BHI medium with 25 µg/mL kanamycin and then in CGXII medium containing 40 g/L of D-gluconate and 25 µg/mL kanamycin. 1 L of CGXII medium with 60 g/L of D-gluconate, 25 µg/mL kanamycin and 1 mM IPTG was used for the main culture, which was prepared to an initial $OD_{600}$ of 1. Temperature and pH of the fermentation were maintained at 30°C and 7, respectively. Shifts in pH were controlled with a 3 M sulfuric acid and a 1 M sodium hydroxide solution. Dissolved oxygen (DO) was set to a minimum value of 30% and controlled via agitation speed (400-1200 rpm) and air flow (60-80 sL/h). After D-gluconate was completely consumed, D-glucose was manually added to the medium to a final concentration of 58 g/L.

**Preparation of XylA$_{XC}$ starting structures**

A resolved 3D structure of the XylA homodimer of *X. campestris* is not available. The starting structure for MD simulations of the homodimeric XylA$_{XC}$ state was thus modeled using AlphaFold3 [39]. The target sequence for XylA$_{XC}$ has Uniprot ID Q4UTU6 [40]. A fasta file with the two monomeric target sequences was prepared and uploaded to the server. The multiple sequence alignments were generated with the default AlphaFold3 workflow. Alphafold3 provides confidence scores in the form of a weighted linear combination of the predicted TM-score (pTM) and the interface predicted TM-score (ipTM). The five resulting models all have a pTM score of 0.82 and an ipTM score of 0.70. The less prevalent α-D-glucopyranose (AGP) structure bound to *S. rubiginosus* D-glucose isomerase in the monomeric configuration together with the coordinating $Mg^{2+}$ ion is available in the Protein Data Bank (PDB ID 4LNC) [41]. We used PDB ID 2XIS to extract the information regarding the position of the second coordinating $Mg^{2+}$ rather than the $Mn^{2+}$ found in PDB ID 4LNC [42]. After the superposition of the experimental protein structures with our generated model (RMSD = 1.9 Å), the bound pose of AGP – $Mg^{2+}$ and the single $Mg^{2+}$ were extracted and merged with the obtained model. Parallely, β-D-glucopyranose (BGP) was docked into the two monomers of the model, using the same protocol as for BFF. The AGP / BGP structures have a ligand dRMSD $<$ 1.5 Å between the ligands with respect to both monomers. We decided to use the docked BGP pose and the $Mg^{2+}$ from the superpositioning step to generate the starting configuration.

The substitutions W147S and G15A-W147S were introduced in XylA following the same protocol as described for the IolT1 mutations. The partial atomic charges of the ligands were taken from the IolT1 setup. The obtained systems were solvated using PACKMOL-Memgen [43,44]. To obtain a neutral system, counter ions were added that replaced solvent molecules (KCl 0.15 M). Two independent starting configurations were generated and are listed in Table S4.

**Preparation of PtsS starting structures**

To our knowledge, there is also no experimentally resolved 3D structure of the D-sucrose phosphotransferase PtsS from *C. glutamicum*. The starting structure for MD simulations of the homodimeric PtsS state was thus modeled using AlphaFold3 [39]. The target sequence for PtsS has Uniprot ID Q8NMD6 [45]. A fasta file with the two monomeric target sequences was prepared and uploaded to the server. The multiple sequence alignments were generated with the default AlphaFold3 workflow. The five resulting models all have a pTM score of 0.57 and an ipTM score of 0.59. The structure of 2-acetamido-2-deoxy-β-D-glucopyranosyl-(1→4)-2-acetamido-2-deoxy-β-D-glucopyranose is available in the Protein Data Bank (PDB ID 3QNQ) bound to the *B. cereus* phosphorylation-coupled saccharide transporter in the homodimeric configuration [46]. After the superposition of the experimental protein structure with our generated model (RMSD = 2.7 Å), the carbohydrate pose was extracted and merged with the obtained model. Additionally, β-D-glucopyranose (BGP), β-D-fructofuranose (BFF), D-sucrose (SUC), and BGP and BFF together (BGP/BFF) were docked into the two monomers of the model, using the same protocol as before and the bound carbohydrate pose to generate the docking grid. BGP/BFF corresponds to the SUC moiety but without a bond between between the two monosaccharides. The carbohydrate structures have a ligand dRMSD < 1.5 Å with respect to both monomers.

The substitution A129G was introduced in PtsS following the same protocol as described for the IolT1 mutations. The partial atomic charges of the ligands were taken from the IolT1 setup. The obtained systems were embedded into a membrane composed of phosphatidylglycerol 16:0-18:1, cardiolipin 16:0-18.1, and phosphatidylinositol 16:0-18:1 in a ratio of 6:2:1 using the PPM server [47] and solvated using PACKMOL-Memgen [43,44] as for IolT1. Two independent starting configurations were generated and are listed in Table S7**.** To check if the proposed substitutions would modify the geometry of the binding site tunnel, the bottleneck radius and the length of the wild type and the variant tunnels were calculated using CAVER 3.0 [37] (Table S8). The COMs of the residues in a distance < 3.0 Å from the original 2-acetamido-2-deoxy-β-D-glucopyranosyl-(1→4)-2-acetamido-2-deoxy-β-D-glucopyranose were defined as the starting points of the search. The probe radius was set to 1.2 Å.

**Relaxation, thermalization, and product runs of the obtained systems**

An initial minimization step was performed with the CPU code of pmemd [48]. Each minimization was organized in four steps of 10000 cycles each, for a total of 40000 cycles of minimization. Afterward, each minimized system was thermalized in one stage from 0 to 300 K over 250 ps using the NVT ensemble and the Langevin thermostat [49], and the density was adapted to 1.0 g $cm^{-3}$ over 750 ps using the NPT ensemble with a semi-isotropic Berendsen barostat [50], with the pressure set to 1 bar. The thermalization and equilibration were performed with the GPU code of pmemd [48]. There are four density adaptation steps with a total time of 1 ns. The sum of thermalization, density adaptation, and equilibration took 50 ns. For each replica, 2 µs of production run using the GPU code of pmemd was performed in the NPT ensemble at a temperature of 300 K using the Langevin thermostat [49] and a collision frequency of 1 $ps^{-1}$. To avoid noticeable distortions in the simulation box size, semi-isotropic pressure scaling using the Berendsen barostat [50] and a pressure relaxation of 1 ps was employed by coupling the box size changes along the membrane plane [51].

**Constraint network analysis**

To detect changes in structural rigidity between IolT1 wildtype and variants, we analyzed ensembles of constraint network topologies based on conformational ensembles saved every 1200 ps from the last 1.5 µs that were considered as production phase of the unbiased MD simulations. For each configuration, we generated 400 conformations. Overall, we investigated 1,600 conformations in this study.

Structural rigidity was analyzed with the CNA software package [52,53], which is a front and back end to the FIRST software [54]. It was used to construct networks of nodes (atoms) and covalent and noncovalent (hydrogen bonds, salt bridges, and hydrophobic tethers) constraints. The hydrogen bond energy (including salt bridges) is determined from an empirical function [55], while hydrophobic tethers between carbon and sulfur atoms were considered if the distance between these atoms was less than the sum of their van der Waals radii plus a cutoff of 0.25 Å [56]. Biomolecules generally display a hierarchy of rigidity that reflects the modular structure of biomolecules in terms of secondary, tertiary, and supertertiary structure [52]. This hierarchy can be identified by gradually removing noncovalent constraints from an initial network representation of a biomolecule, which generates a succession of network states $\sigma$, resulting in a "constraint dilution trajectory". For that, hydrogen bonds and salt bridges are removed in the order of increasing strength such that for network state $\sigma$, only those hydrogen bonds are kept that have an energy $E_{HB} \leq E_{cut}(\sigma)$. In our analysis, we used energy cutoffs between $E_{start}$ = -0.1 kcal/mol and $E_{stop}$ = -10.0 kcal/mol, with $E_{step}$ = 0.1 kcal/mol. We also assigned the cutoff for finding native contacts between pairs of residues (ncd = 4.5).

For all MD-generated snapshots, we calculated a rigidity index $r_i$ [57] (eq. 1).

$$r_i = min\{E_{cut} | \exists c \in C^{E_{cut}} : A_{i1} \wedge A_{i2} \in c\} \qquad \text{(eq. 1)}$$

The index is defined for each covalent bond $i$ between two atoms $A_i$, [58] as the $E_{cut}$ value during a constraint dilution simulation at which the bond changes from rigid to flexible. Phrased differently, this index monitors when a bond segregates from any rigid cluster $c$ of the set of rigid clusters $C^{E_{cut}}$. It reflects structural stability on a per-residue basis and, thus, can be used to identify the location and distribution of structurally weak or strong parts in IolT1.

The same analysis was performed for the five independent replicas of each IolT1 configuration for the WT, the G87S and T351P single variants, and the G87S-T351P double variant. The per-residue $r_i$ values were averaged on a per-residue basis across the five replicas and the SEM was calculated. Afterward, the per-residue difference $\Delta r_{i,(Mut-WT)}$ (eq. 2) was calculated, including an error propagation analysis for the SEM. Values of $\Delta r_{i,(Mut-WT)} < 0$ (> 0) indicate that the variant is more rigid (flexible) than the wildtype.

$$\Delta r_{i,(Mut-WT)} = r_{i,(Mut)} - r_{i,(WT)} \qquad \text{(eq. 2)}$$

Finally, we obtained the cumulative difference $\Delta r_{r,(Mut-WT)}$ of a specific region of interest, $r_r$ (residues 290-306 and residues 390-401) (eq. 3) by summing the $m$ single per-residue $r_{i,(Mut-WT)}$ contributions, including an error propagation analysis.

$$\Delta r_r = \sum_i^m \Delta r_{i,(Mut-WT)} \qquad \text{(eq. 3)}$$
